# Highly diverse flavobacterial phages as mortality factor during North Sea spring blooms

**DOI:** 10.1101/2021.05.20.444936

**Authors:** Nina Bartlau, Antje Wichels, Georg Krohne, Evelien M. Adriaenssens, Anneke Heins, Bernhard M. Fuchs, Rudolf Amann, Cristina Moraru

## Abstract

It is generally recognized that phages have a modulating role in the marine environment. Therefore, we hypothesized that phages can be a mortality factor for the dense heterotrophic bacterial population succeeding in phytoplankton blooms. For the marine carbon cycle, spring phytoplankton blooms are important recurring events. In this study, we focused on *Flavobacteriia*, because they are main responders during these blooms and have an important role in the degradation of polysaccharides. A cultivation-based approach was used, obtaining 44 lytic flavobacterial phages (flavophages), representing twelve new species from two viral realms – *Duplodnaviria* and *Monodnaviria*. Taxonomic analysis allowed us to delineate ten new phage genera and seven new families, from which nine and four, respectively, had no previously cultivated representatives. Genomic analysis predicted various life styles and genomic replication strategies. A likely eukaryote-associated host habitat was reflected in the gene content of some of the flavophages. Detection in cellular metagenomes and by direct-plating indicated that part of these phages were actively replicating in the environment during the 2018 spring bloom. Furthermore, CRISPR/Cas spacers and re-isolation during two consecutive years indicated that, at least part of the new flavophages are stable components of the microbial community in the North Sea. Together, our results indicate that these diverse flavophages have the potential to modulate their respective host populations.

## Introduction

Marine bacteriophages outnumber their hosts by one order of magnitude in surface seawater and infect 10-45% of the bacterial cells at any given time (1–3). They have a major impact on bacterioplankton dynamics. This impact can be density dependent (4) and take many forms. By lysing infected cells, viruses decrease the abundance of their host population, shifting the dominant bacterial population, and recycling the intracellular nutrients inside the same trophic level (5). By expressing auxiliary metabolic genes, phages can enhance the metabolic capabilities of the virocells (6, 7). By transferring pieces of host DNA, they can drive bacterial evolution (8). By blocking superinfections with other phages (9), they can protect from immediate lysis. Potentially phages even influence carbon export to the deep ocean due to aggregation of cell debris resulted from cell lysis, called the viral shuttle (10, 11). Marine phages modulate not only their hosts, but also the diversity and function of whole ecosystems. This global impact is reflected in a high phage abundance (12) and diversity (13, 14).

Phages are also well known for modulating bacterial communities in temperate coastal oceans. Here, the increase in temperature and solar radiation in spring induces the formation of phytoplankton blooms (15), which are globally important components of the marine carbon cycle. These ephemeral events release high amounts of organic matter, which fuels subsequent blooms of heterotrophic bacteria. *Flavobacteriia* and *Gammaproteobacteria* are the main responders. Recurrent genera like *Polaribacter, Maribacter*, and *Pseudoalteromonas* succeed each other in a highly dynamic fashion. This bacterial succession is likely triggered by the availability and quality of substrates such as polysaccharides (16, 17), yet it cannot be fully understood without considering mortality factors such as grazing by protists, and viral lysis. Grazing by protists is mostly size dependent (18), whereas viral lysis is highly host-specific (19).

Based on the availability of suitable host bacteria, marine phages can be obtained with standard techniques. Over the years notable numbers of phages infecting marine *Alphapoteobacteria* (e.g. (20, 21)), *Gammproteobacteria* (e.g. (22)), and *Cyanobacteria* (e.g. (23, 24)) have been isolated. Despite the importance of *Flavobacteriia* as primary degraders of high molecular weight algal derived matter only few marine flavobacterial phages, to which we refer in the following as flavophages, have been characterized. This includes several *Cellulophaga* phage isolates from the Baltic Sea, covering all morphological types in *Duplodnaviria* and also two different phage groups in *Monodnaviria* (19, 25). In addition, eight tailed phages, one having a myoviral and the rest a siphoviral morphology, were isolated for members of the genera *Polaribacter* (26), *Flavobacterium (27, 28), Croceibacter* (29) or *Nonlabens* (30). However, still the coverage of the class *Flavobacteriia* and the diversity of marine flavophages remains low. With the exception of the *Cellulophaga* phages most of the other flavophages have only been briefly characterized in genome announcements.

In the context of a large project investigating bacterioplankton successions during North Sea spring bloom season, we isolated and characterized new flavophages, with the purpose of assessing their ecological impact and diversity. In total, more than 100 phage isolates were obtained, sequenced, annotated, and classified. This diverse collection is here presented in the context of virus and bacterioplankton abundances. All newly isolated flavophage genomes were mapped to metagenomes obtained for Helgoland waters of different size fractions, testing the environmental relevance of the flavophage isolates. This study indicates that flavophages are indeed a mortality factor during spring blooms in temperate coastal seas. Furthermore it provides twelve novel phage-host systems of six genera of *Flavobacteriia*, doubling the number of known hosts.

## Material and methods

### Sampling campaigns

Surface water samples were taken off the island Helgoland at the long term ecological research station Kabeltonne (54° 11.3’ N, 7° 54.0’ E). The water depth was fluctuating from 7 to 10 m over the tidal cycle. In 2017, a weekly sampling was conducted over five weeks starting on March 14 (Julian days 73-106) and covered the beginning of a spring phytoplankton bloom. In 2018, sampling was conducted over eight weeks starting on March 29 and ending on May 24 (Julian days 88-145). It covered the full phytoplankton bloom. Additional measurements were performed to determine the chlorophyll concentration, total bacterial cell counts and total virus abundances (SI file 1 text). Viruses were counted both by epifluorescence microscopy of Sybr Gold stained samples and by transmission electron microscopy (TEM) of uranyl acetate stained samples (SI file 1 text).

### Phage isolation

Phage isolates were obtained either after an intermediate liquid enrichment step or by direct plating on host lawns (SI file 1 Table1). In both cases, seawater serially filtered through 10 μm, 3 μm, 0.2 μm polycarbonate membranes served as phage source. To ensure purity, single phage plaques were transferred three times and then a phage stock was prepared. For more details, see SI file 1 text. These phage stocks were then used for assessing phage morphology, host range and genome size, and for DNA extraction (SI file 1 text).

### Determination of phage genomes

Phage DNA was extracted using the Wizard resin kit (Promega, Madison, USA) and eluted in TE buffer (after (31)). The DNA was sequenced on a Illumina HiSeq3000 applying the paired-end 2 x 150 bp read mode. For most *Cellulophaga* phages (except Ingeline) and *Maribacter* phages a ChIPSeq library was prepared. For the other phages a DNA FS library was prepared. The raw reads were quality trimmed and checked, then assembled using SPAdes (v3.13.0, (32)) and Tadpole (v35.14, sourceforge.net/projects/bbmap/). Assembly quality was checked with Bandage (32). The genome ends were determined using PhageTerm (33). For more details about all these procedures, see SI file 1 text.

### Retrieval of related phage genomes

Several publicly available datasets of environmental phage contigs (13, 33-38) were queried for sequences related with the flavophages isolated in this study, in a multistep procedure (SI file 1 text). All environmental phage contigs and the reference genomes that formed a cluster in vConTact2 (39) with the flavophage isolates were selected for further analysis. Only environmental contigs with a length >80% compared to their related flavophage isolate were kept.

### Phage genome annotation

All phage genomes analysed in this study were annotated using a custom bioinformatics pipeline described elsewhere (20), with modifications (SI file 1 text). The final protein annotations were evaluated manually.

### Phage genome taxonomic assignment

The dataset used for phylogenetic analysis comprised of i) all flavophage isolates from this study; ii) related phage genomes (see above); and iii) all *Caudovirales* phage genomes from the Master Species List release 35 (March 2020) (40) from the International Committee of Taxonomy of Viruses (ICTV).

To determine the family level classification of the flavophages we used ViPTree (41) and VICTOR (42) to calculate whole proteome phage trees. First, the command line ViPTree was applied on the full phage genome dataset and different clades were assigned to families, taking as reference the phylogenetic distances for the *Herelleviridae* family (43). Then, a subset of phages comprising all family level clades with the new flavophages was selected for analysis with VICTOR. The parameters for VICTOR were “amino acid” data type and the “d6” intergenomic distance formula. In addition to phylogenetic trees, VICTOR used the following predetermined distance thresholds to suggest taxon boundaries at subfamily (0.888940) and family (0.985225) level (42).

To calculate the sequence relatedness of the ssDNA virus Omtje, the web service of GRAViTy (http://gravity.cvr.gla.ac.uk, (44)) was used with the “Obscuriviridae” and the Baltimore Group II - ssDNA viruses + *Papillomaviridae* and *Polyomaviridae* (VMRv34).

To determine the intra-familial structure and relationships, smaller phage genome datasets corresponding to each family were analysed using i) nucleic acid based intergenomic similarities calculated with VIRIDIC (viridic.icbm.de) and ii) core protein phylogeny. The thresholds used for species and genus definition were 95% and 70% intergenomic similarity, respectively. The core genome analysis was conducted as follows: i) core genes were calculated with the CoGePh web tool (cogeph.icbm.de); ii) a multiple alignment of all concatenated core proteins was constructed with MUSCLE (v3.8.425,(45)) and iiia) used for the calculation of an approximately maximum-likelihood phylogenetic tree using FastTree v 2.1.11 (46), integrated as plugin in Geneious v 11.1.5 (http://www.geneious.com, (47). In addition, iiib) IQ-Trees with SH-aLRT (48) and ultrafast boostrap values (49) using ModelFinder (50) were calculated. All phylogenetic trees were visualized using FigTree v1.4.4. (51), available at http://tree.bio.ed.ac.uk/software/figtree/).

### Host assignment for environmental phage genomes

To determine potential hosts for the environmental phage genomes, a BLASTN (52) search (standard parameters) was performed against the nucleotide collection (nr/nt, taxid:2, bacteria), for all phages and environmental contigs belonging to the newly defined viral families. The hit with the highest bitscore and annotated genes was chosen to indicate the host.

### Detection of flavophages and their hosts in Helgoland metagenomes by read mapping

The presence of flavophages, flavobacterial hosts and environmental phages in unassembled metagenomes from the North Sea and their relative abundances were determined by read mapping, using a custom bioinformatics pipeline (20). Two datasets were analyzed: i) cellular metagenomes (0.2 – 3 μm fraction) from the spring 2016 algal bloom (SI file 1 Table 2) and ii) cellular metagenomes (0.2 – 3 μm, 3 - 10 μm and >10 μm fractions) from the spring 2018 algal bloom (SI file 1 Table 2).

A bacterium was considered present when > 60% of the genome was covered by reads with at least 95% identity. A phage was present when > 75% of its genome was covered by reads with at least 90% identity. Relatives of a phage were present when > 60% of its genome was covered by reads with at least 70% identity. The relative abundance of a phage/host genome in a sample was calculated by the following formula: “number of bases at ≥x% identity aligning to the genome / genome size in bases / library size in gigabases (Gb)”.

### Host 16S rRNA analysis

The genomes of those bacteria which gave phages were sequenced with Sequel I technology (Pacific Biosciences, Menlo Park, USA) (SI file 1 text). After genome assembly the quality was checked and 16S rRNA operons were retrieved using the MiGA online platform (53). Additionally, the 16S rRNA gene from all the other bacterial strains was amplified and sequenced using the Sanger technology (see SI file 1).

For phylogenetic analysis, a neighbour joining tree with Jukes-Cantor correction and a RaxML tree (version 8, (54)) were calculated using ARB (55). The reference data set Ref132 was used, with the termini filter and Capnocytophaga as outgroup (56). Afterwards, a consensus tree was calculated.

### CRISPR spacer search

CRISPR spacers and cas systems were identified in the host genomes by CRISPRCasFinder (57). Extracted spacers were mapped with Geneious to the flavophage genomes.

The IMG/VR (36) web service was used to search for spacers targeting the flavophage isolates and the related environmental genomes. A BlastN against the viral spacer database and the metagenome spacer database were run with standard parameters.

### Data availability

Sequencing data and phage genomes are available at the National Center for Biotechnology Information (NCBI) with the accession number PRJNA639310. Phages (DSM111231-111236, DSM111238-111241, DSM111252, and DSM111256-111257) and bacterial hosts (DSM111037-111041, DSM111044, DSM111047-111048, DSM111061, and DSM111152) were deposited at the German Collection of Microorganisms and Cell Cultures GmbH, Braunschweig, Germany.

## Results

Spring phytoplankton blooms were monitored by chlorophyll *a* measurements (Figure 1, SI file 1 Figure 1). In 2018, the bloom had two chlorophyll *a* peaks, and it was more prominent than in 2017. Diatoms and green algae dominated the 2018 blooms (SI file 1 Figure 2). During both blooms, bacterial cell numbers almost tripled, from ~ 6.5×10^5^ cells ml^-1^ to ~ 2×10^6^ cells ml^-1^. The *Bacteroidetes* population showed a similar trend, as revealed by 16S rRNA data (Figure 1).

**Figure 1:**
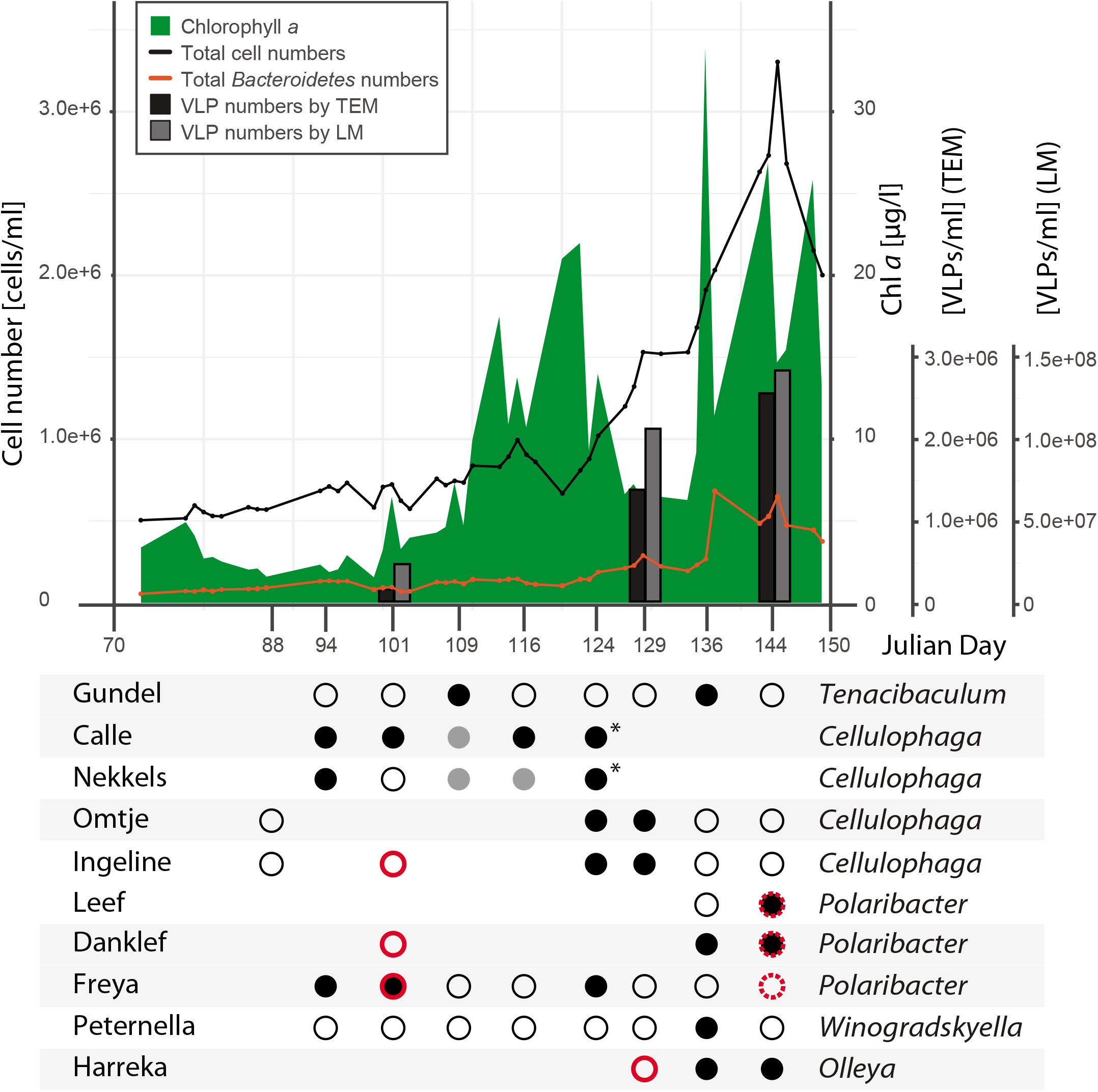
Flavophage detection during the 2018 spring phytoplankton bloom, as inferred from phage isolation and metagenome read mapping. Upper panel: Results are presented in the context of chlorophyll a concentration (green), total cell counts (black line), Bacteroidetes counts (orange line), phage counts by transmission electron microscopy (TEM, black bar), and phage counts by epifluorescence light microscopy (LM, grey bar). Lower panel: Phage isolation is shown for different time points (x axis, Julian days), with the phage identity verified by sequencing (black dots) or not determined (gray dots). Most of the isolations were done by enrichment, and just a few by direct plating (asterisks). Phage detection in metagenomes was performed by read mapping, with a 90% read identity threshold for phages in the same species (full red circles) and 70% read identity threshold for related phages (dashed red circles). Alternating gray shading of isolates indicates phages belonging to the same family.

### Viral counts

Viral particles were counted at three time points during the 2018 bloom, both by SYBR Gold staining and TEM. Numbers determined by TEM were almost two orders of magnitude lower than those determined by SYBR Gold (Figure 1), a typical phenomenon when comparing these two methods (58). However, both methods showed a strong increase of viral particle numbers over the course of the bloom. All viruses counted by TEM were tailed, a strong indication that they were infecting bacteria or archaea, but not algae (59) (SI file 1 Figure 3). The capsid size ranged between 54 and 61 nm, without any significant differences between the three time points (SI file 1 Figure 4). The virus to bacteria ratio increased throughout the bloom, almost doubling (Figure 1, SI file 1 Table 3).

### Flavophage isolation and classification

For phage enrichment, 23 bacterial strains previously isolated from algal blooms in the North Sea were used as bait (SI file 1 Table 1). A total of 108 phage isolates were obtained for 10 of the bacterial strains. These were affiliated with the bacterial genera *Polaribacter, Cellulophaga, Olleya, Tenacibaculum, Winogradskyella*, and *Maribacter* (Figure 2, SI file 1 Table 4).

**Figure 2:**
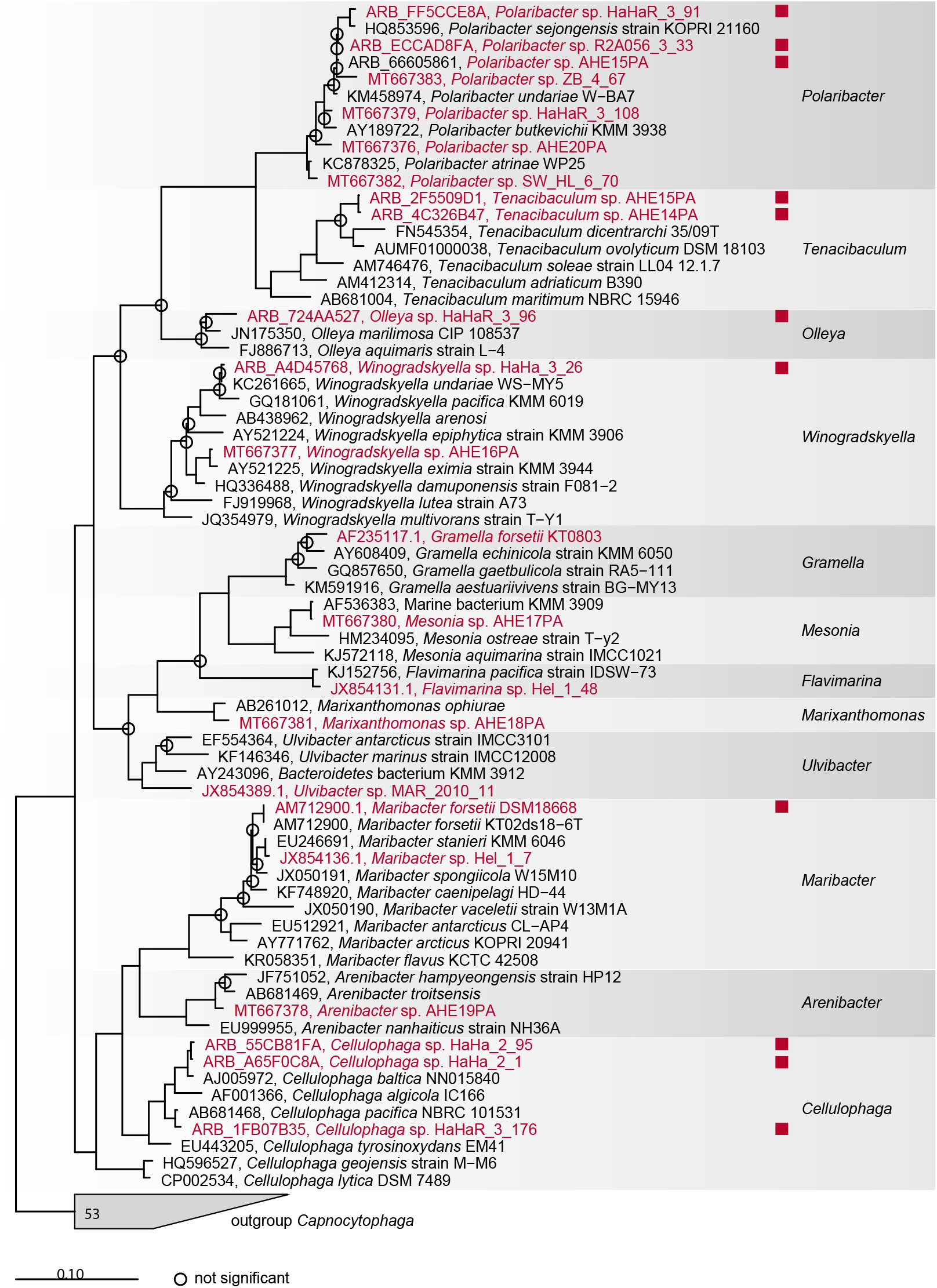
Phylogenetic tree (consensus between raxml and the NJ) of the 16S rRNA gene from all bacterial strains used to enrich for phages in 2017 and 2018 (red), plus reference genomes. Red squares indicate successful phage isolation.

Intergenomic similarities at the nucleic acid level allowed the grouping of the 108 flavophages into 44 strains (100% similarity threshold) and 12 species (95% similarity threshold) (SI file Figure 5). The species were tentatively named “Olleya virus Harreka” (Harreka), “Tenacibaculum virus Gundel” (Gundel), “Maribacter virus Molly” (Molly), “Maribacter virus Colly” (Colly), “Winogradskyella virus Peternella” (Peternella), “Polaribacter virus Danklef” (Danklef), “Polaribacter virus Freya” (Freya), “Polaribacter virus Leef” (Leef), “Cellulophaga virus Nekkels” (Nekkels), “Cellulophaga virus Ingeline” (Ingeline), “Cellulophga virus Calle” (Calle), and “Cellulophaga virus Omtje” (Omtje). The names have a Frisian origin, to reflect the flavophage place of isolation. Molly and Colly were isolated only in 2017, Ingeline and Omtje both in 2017 and 2018, and the remaining in 2018 (SI file Figure 1).

Virion morphology as determined by TEM (Figure 3) showed that 11 of the phages were tailed. According to the new megataxonomy of viruses, tailed phages belong to the realm *Duplodnaviria*, kingdom *Heunggongvirae*, phylum *Uroviricota*, class *Caudoviricetes*, order *Caudovirales (60)*. The only non-tailed phage was Omtje. Further digestion experiments with different nucleases showed that Omtje has a ssDNA genome (SI file 1, Figure 6). Therefore, this phage belongs to the realm *Monodnaviria*. The non-tailed, icosahedral capsid morphology (Figure 3) places Omtje in the *Sangevirae* kingdom.

**Figure 3:**
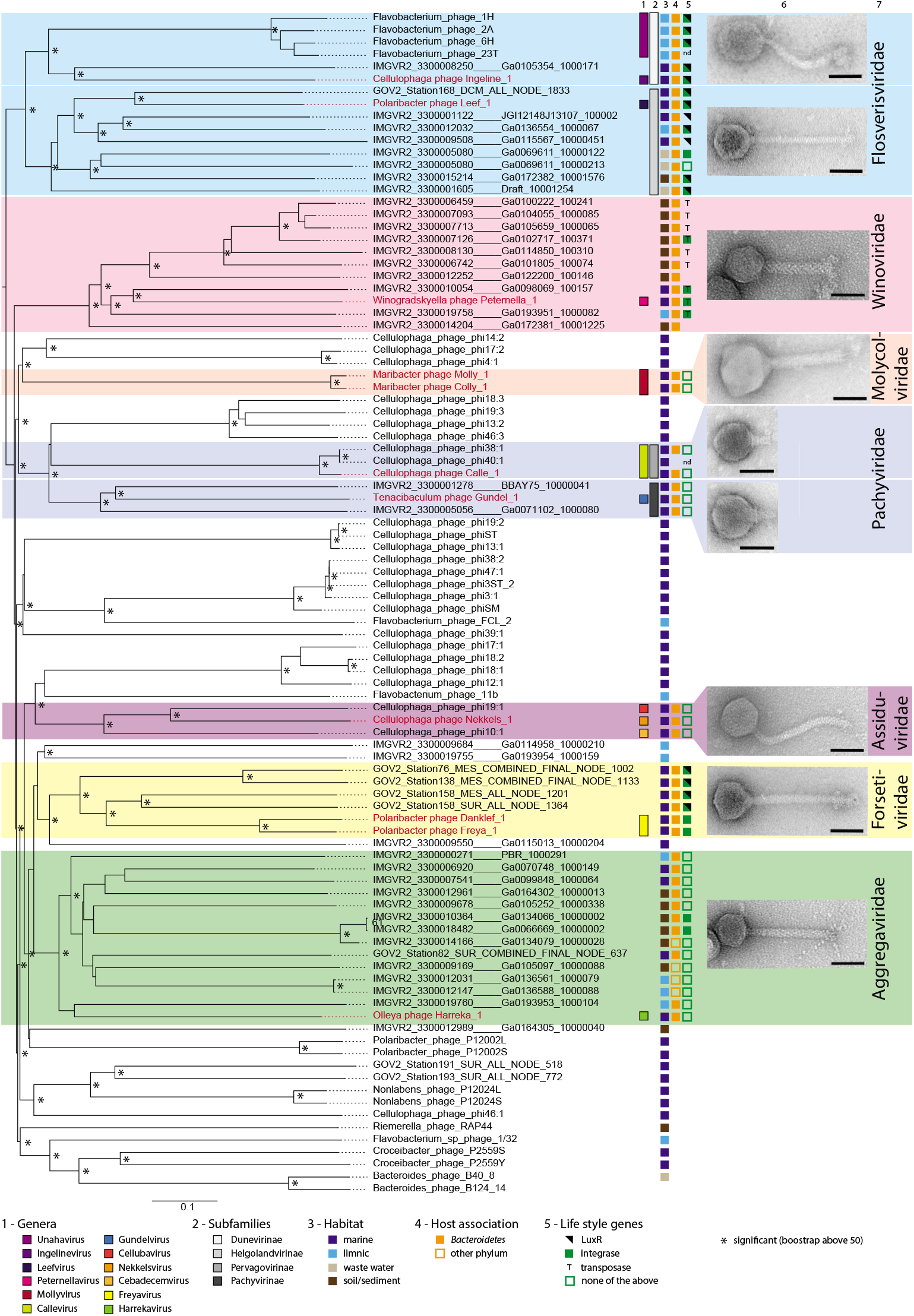
Whole genome phylogeny determined with VICTOR (amino-acid based) for the dsDNA flavophages, including environmental sequences and reference phage genomes. Our new phage isolates are depicted in red. Pseudo-bootstrap values are indicated at branches. Column 6 shows the TEM image of the negative stained new flavophage (scale bar in each TEM image 50 nm) and column 7 gives the new family name.

Phylogenetic comparison using ViPTree of the 11 tailed flavophages with all *Caudoviricetes* phages in the ICTV35 database showed that they fall into 7 monophyletic clades of similar rank with the newly defined *Herelleviridae* family (61) (SI file Figure 7). In a further analysis of selected phage genomes using VICTOR, the same monophyletic clades were obtained (Figure 3). All were maximally supported, with pseudo boostrap values of 100, with the exception of the clade containing the Harreka_1 phage, which had a pseudo bootstrap value of 67. To most of these clades VICTOR assigned the rank of subfamily (SI file 1 Figure 7). In the light of the current changes in the ICTV phage classification, this corresponds to the family rank (61). Therefore, taking both the ViPTree and the VICTOR results into consideration, we are proposing that the 7 clades correspond to 7 new families, which we tentatively named as follows: “Flosverisviridae”, “Winoviridae”, “Molycolviridae”, “Pompuviridae”, “Assiduviridae”, “Forsetiviridae” and “Aggregaviridae” (Figure 3).

Four of the new families (“Aggregaviridae”, “Forsetiviridae”, “Molycolviridae” and “Winoviridae”) were formed only from new flavophages and from environmental phage genomes. The remaining three also contained previously cultivated phages, infecting bacteria from the genus *Cellulophaga* (“Pompuviridae” and “Assiduviridae”) and *Flavobacterium* (“Flosverisviridae”).

Using a 70% threshold for the intergenomic similarities at nucleotide level indicated that Harreka, Nekkels, Gundel, Peternella, Leef and Ingeline phages form genera on their own, tentatively named here “Harrekavirus”, “Nekkelsvirus”, “Gundelvirus”, “Peternellavirus”, “Leefvirus” and “Ingelinevirus”. The other new flavophages formed genera together with isolates from this study or with previously isolated flavophages, as follows: the genus “Freyavirus” formed by Danklef and Freya, the genus “Callevirus” formed by Calle, Cellulophaga phage phi38:1, Cellulophaga phage phi40:1, and the genus “Mollyvirus” formed by Molly and Colly (SI file Figure 9). For each family containing more than 3 phages, a phylogenetic analysis of the core proteins, based on the determined core genes, was performed (SI file 1 Figure 10 - 20). The trees obtained supported the intra-familial structure obtained with VICTOR, and allowed us to define two subfamilies in “Pompuviridae” (“Pachyvirinae” and “Pervagovirinae”) and two in “Flosverisviridae” (“Helgolandvirinae” and “Duenevirinae”), see Figure 3 and SI file 1 Figure 10, 11, 14, 15.

Phylogenetic analysis using both ViPTree and VICTOR of a dataset including the genome of Omtje and all ssDNA phages from the ICTV35 release, and of Omtje using GRAViTy, showed that Omtje and previously isolated *Cellulophaga* phages form a clade separated from the known *Microviridae* family (Figure 4, SI file 1 Figure 21 - 23). We propose here that this clade represents a new family, tentatively called here “Obscuriviridae”. The placement of this family into higher taxonomic ranks remains to be determined in the future. Intergenomic similarity calculations indicate that Omtje forms a genus by itself (SI file 1 Figure 24).

**Figure 4:**
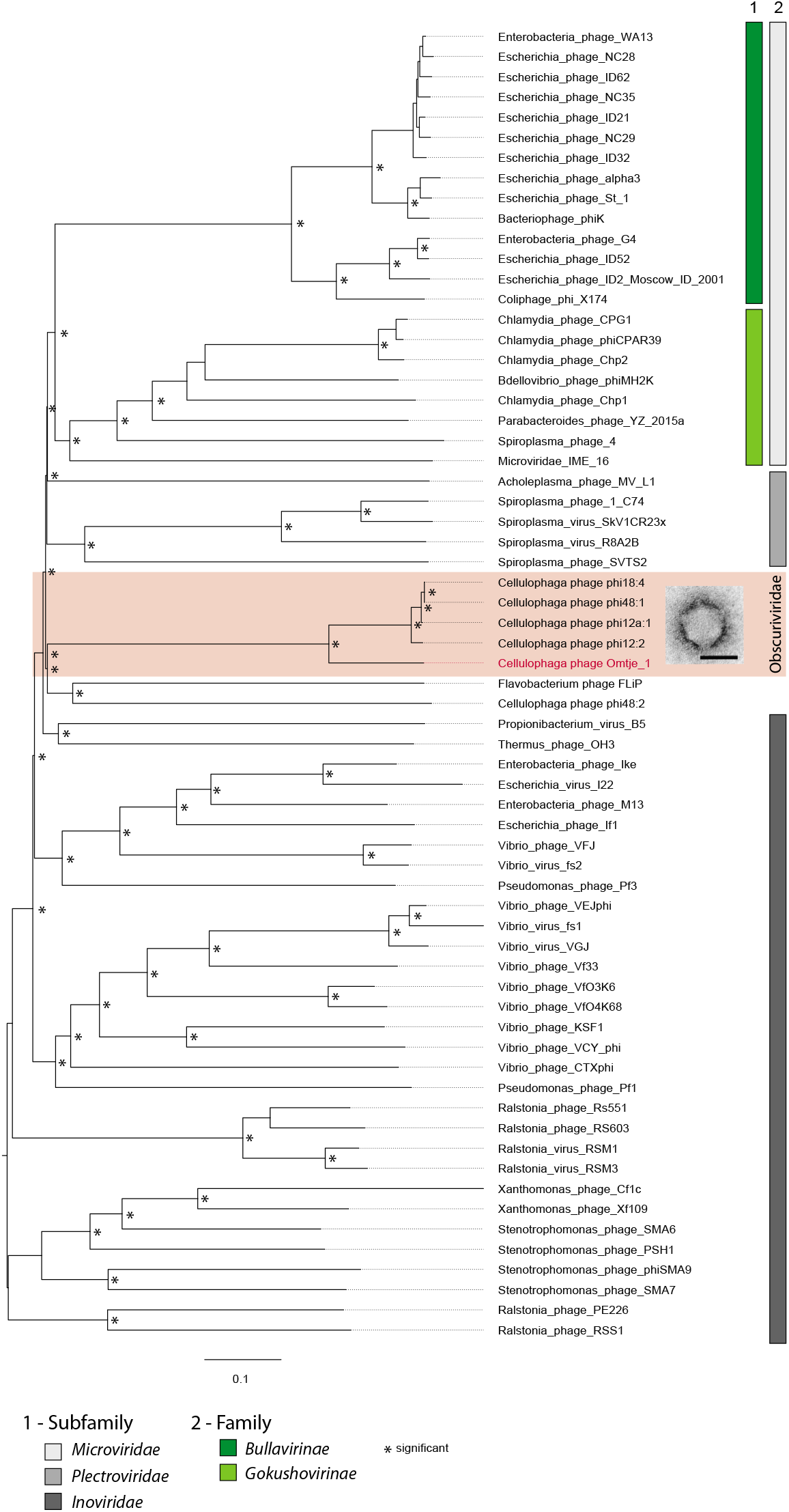
Whole genome phylogeny determined with VICTOR (amino-acid based) for the ssDNA phages. Our new phage isolate is depicted in red. Pseudo-bootstrap values are indicated at branches. Column 6 shows the TEM image of the negative stained new flavophage (scale bar in each TEM image 50 nm) and column 7 gives the new family name.

During the phylogenetic analysis we have worked closely with ICTV members, to ensure a good quality of the phage taxonomic affiliations. Two taxonomic proposals for the new defined taxons are being prepared, one for flavophages in *Duplodnaviria* and one for *Monodnaviria*.

### Features of the new flavophage isolates

#### Phages of *Polaribacter*

Three *Polaribacter* phages were obtained: i) Danklef and Freya, part of the family “Forsetiviridae”, infected *Polaribacter* sp. R2A056_3_33 and HaHaR_3_91, respectively; and ii) Leef, part of the family “Flosverisviridae”, infected *Polaribacter* sp. AHE13PA (SI file Figure 1). These phages infected only their isolation host (SI file 1 Figure 18). They all had a siphoviral morphology (Figure 3 and Table 1).

**Table 1:** Phage characteristics of each phage group. As Colly has the same characteristics as Molly, it is not especially mentionied in the table. Lysis genes are stated as N-acetylmuramidase (M), N-aceylmuramoyl-L-alanine-amidase (A), Glycosid Hydrolase 19(GH), L-Alanine-D-glutamine-peptidase (P), spanin (S), holin(H).

| Phage genus | host genus | genome size [kbp] | %GC | ORFs | # tRNAs | # tmRNAs | # integrases | lysis genes | genome ends | circularly closed | capsid diameter [nm] | tail length [nm] | tail width [nm] | morphology | # strains |
| --- | --- | --- | --- | --- | --- | --- | --- | --- | --- | --- | --- | --- | --- | --- | --- |
| Molly | *Maribacter* | 124.2 | 36.2 | 193-201 | 0 | 0 | 0 | A | nd | yes | 74.9±3.6 | 101.5±6.3 | 18.1±1.6 | myoviridal | 7 |
| Gundel | *Tenacibaculum* | 78.5 | 30.4 | 118 | 10 | 0 | 0 | P | short DTRs | yes | 60.5±5.2 | 22.7±3.2 | - | podoviridal | 1 |
| Calle | *Cellulophaga* | 73.0 | 38.1 | 85-86 | 20 | 1 | 0 | A | nd | yes | 60.3±3.0 | 23.0±5.5 | 13.5±2.4 | podoviridal | 3 |
| Nekkels | *Cellulophaga* | 53.3-54.3 | 31.5 | 93-94 | 0 | 0 | 0 | GH, S | nd | yes | 57.8±5.0 | 141.0±7.9 | 13.0±1.5 | siphoviridal | 2 |
| Omtje | *Cellulophaga* | 6.6 | 31.2 | 13 | 0 | 0 | 0 | A | nd | yes | 52.3±4.6 | - | - | non tailed | 5 |
| Ingeline | *Cellulophaga* | 42.6 | 32.2 | 49-50 | 0 | 0 | 2 | S | nd | yes | 59.0±5.3 | 132.9±19.3 | 11.2±1.7 | siphoviridal | 8 |
| Leef | *Polaribacter* | 37.5 | 29.7 | 48 | 0 | 0 | 2 | M | Cos 3' | yes | 49.2±3.6 | 138.7±9.6 | 11.1±2.0 | siphoviridal | 1 |
| Danklef | *Polaribacter* | 47.2-48.2 | 28.9 | 76-78 | 1 | 0 | 1 | M | nd | yes | 46.1±2.2 | 157.4±4.6 | 12.1±1.8 | siphoviridal | 5 |
| Freya | *Polaribacter* | 44.0-48.9 | 28.9 | 66-78 | 0 | 0 | 1 | 0 | nd | yes | 53.9±4.7 | 151.0±8.2 | 13.1±2.0 | siphoviridal | 10 |
| Peternella | *Winogradskyella* | 39.6 | 35.3 | 62 | 0 | 0 | 1 | A, H x | Mu-like | no | 52.3±4.2 | 105.8±7.4 | 16.4±2.4 | myoviridal | 1 |
| Harreka | *Olleya* | 43.2 | 32.0 | 80 | 0 | 0 | 0 | GH | nd | yes | 44.3±3.6 | 123.8±8.0 | 14.3±2.0 | myoviridal | 1 |

The genome size ranged between ~ 38 and ~ 49 kbp. The GC content was very low (28.9-29.7%). For Leef, PhageTerm predicted genome ends of type Cos 3’ (Table 1). Three types of structural proteins were identified in all phages, a major capsid protein, a tail tape measure protein and a portal protein. Several genes for DNA replication, modification and nucleotide metabolism genes were found (Figure 5 and SI file 2). An N-acetylmuramidase was detected in Danklef and Leef, surrounded by transmembrane domains (TMDs) containing proteins. All three phages encoded an integrase. Leef had also a LuxR protein and a pectin lyase.

**Figure 5:**
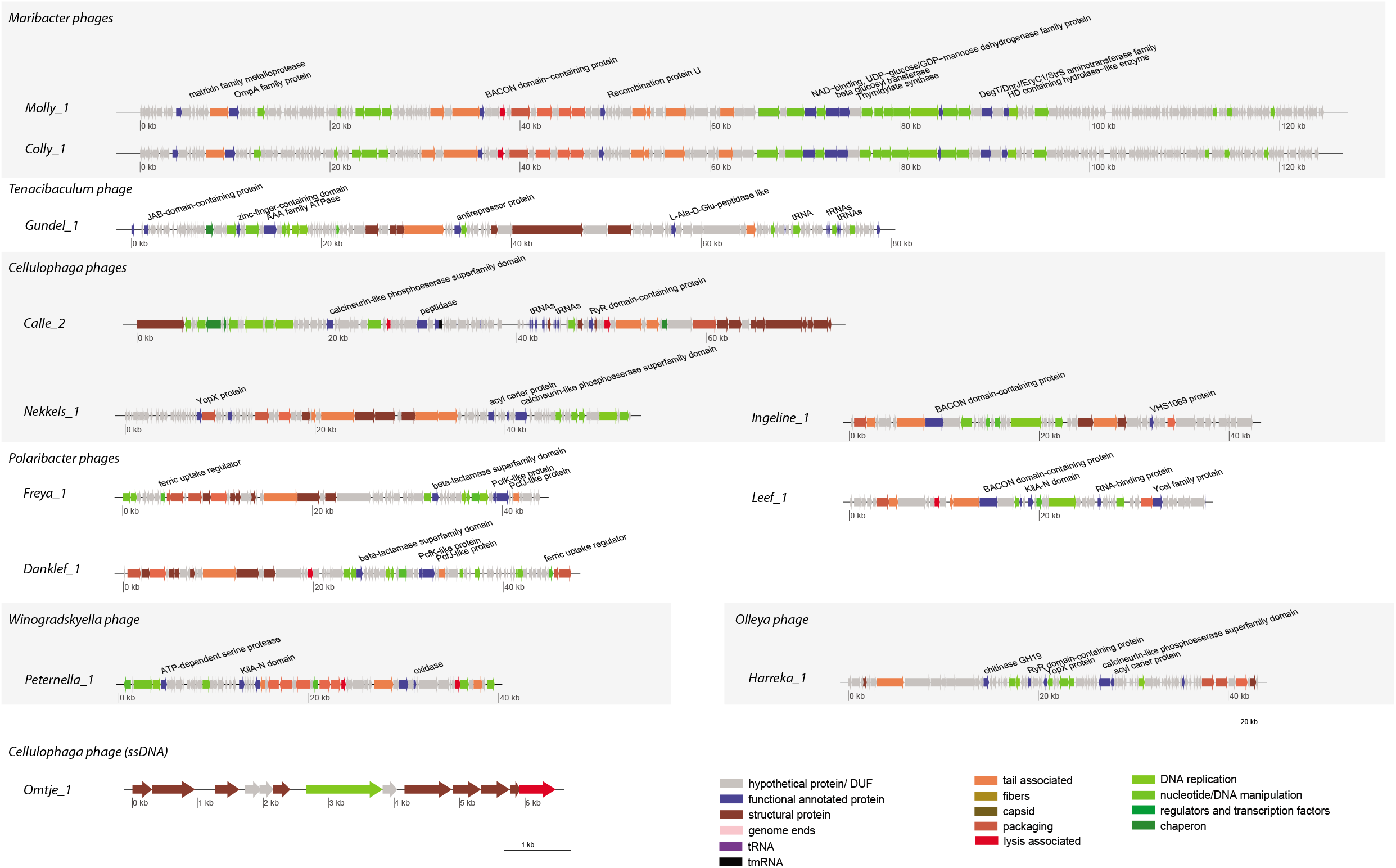
Genome maps of phage isolates with color-coded gene annotations.

#### Phages of *Cellulophaga*

Four Cellulophaga phages were obtained: i) Calle, part of the family “Pompuviridae”, infected *Cellulophaga* sp. HaHa_2_95; ii) Nekkels, part of the family “Assiduviridae”, infected *Cellulophaga* sp. HaHa_2_1; iii) Ingeline, part of the family “Flosverisviridae”, and Omtje, part of the family “Obscuriviridae”, infected *Cellulophaga* sp. HaHaR_3_176 (SI file 1 Figure 1). Nekkels and Calle also infected other bacterial strains than their isolation host, although at a low efficiency: *Polaribacter* sp. AHE13PA (Nekkels) and Cellulophaga sp. HaHa_2_1 (Calle) (SI file 1 Figure 25). The virion morphology varied from icosahedral, non-tailed, microvirus-like for Omtje, to tailed, podovirus-like for Calle and siphovirus-like for Nekkels and Ingeline (Figure 3 and Table 1).

The genome size ranged from ~ 6.5 kb (Omtje) to ~ 73 kbp (Calle). The G+C content varied between 31.2 and 38.1% (Table 1). The few structural genes recognized were in agreement with the virion morphology (Figure 5). For example, Ingeline and Nekkels had a tape measure protein, and a portal protein was found in all three tailed phages. From the replication genes, we detected in Omtje a replication initiation protein specific for ssDNA phages, and in Calle a DNA polymerase A (Figure 5).

Potential endolysins found were the mannosyl-glycoprotein endo-beta-N-acetylglucosaminidase in Omtje and the N-aceylmuramoyl-L-alanine-amidase in Calle. The latter had in its vicinity two proteins with a TMD. In Ingeline we annotated a unimolecular spanin. Nekkels encoded a glucoside-hydrolase of the GH19 family, with a peptidoglycan-binding domain. In its vicinity there were three proteins with three to six TMDs and one potential unimolecular spanin (Table 1, SI file 2).

Pectin and chondroitin/alginate lyase domains were found in Ingeline and Nekkels, as part of bigger proteins, also carrying signaling peptides. Nekkels also encoded a Yersinia outer protein X (YopX), a potential virulence factor against eukaryotes. Ingeline encoded an integrase and a LuxR gene. Calle had 20 tRNA and one tmRNA genes (Figure 5, SI file 2).

#### Phages isolated from other *Flavobacteriaceae*

Five phages were obtained from four other flavobacterial hosts: i) Harreka, part of “Aggregaviridae”, infected *Olleya* sp. HaHaR_3_96; ii) Peternella, part of “Winoviridae”, infected *Winogradskyella* sp. HaHa_3_26; iii) Gundel, part of “Pompuviridae”, infected *Tenacibaculum* sp. AHE14PA and AHE15PA; and iv) Molly and Colly, part of “Molycolviridae”, infected *Maribacter forsetii* DSM18668 (Figure 3). Harreka infected also *Tenacibaculum* sp. AHE14PA and AHE15PA with a significantly lower infection efficiency (SI file 1 Figure 25). All virions were tailed, with a podoviral morphology for Gundel, and a myoviral morphology for Molly, Peternella and Harreka (Table 1). The genome size ranged from ~ 40 kbp to ~ 125 kbp and the G+C content between 30.4 and 36.2%. Gundel had short direct terminal repeats (DTRs) as genome ends (Table 1).

In these phages we recognized several structural, DNA replication and modification, and nucleotide metabolism genes (Figure 5). With respect to replication, Molly and Colly had a DNA polymerase I gene, plus a helicase and a primase. In Gundel and Harreka we only found a DNA replication protein. In Peternella we found a MuA transposase, several structural genes similar to the Mu phage (62) and hybrid phage/host genome reads, indicating that Peternella is likely a transducing phage.

Potential lysins were a N-acetylmuramoyl-L-alanine-amidase in Molly and Peternella and a L-Alanine-D-glutamine-peptidase in Gundel. Harreka had a glycoside-hydrolase of the GH19 family. In the genomic vicinity of the potential lysins, several proteins having 1-4 TMDs were found, including a holin in Peternella (Table 1, SI file annots).

Additional features of these phages were: i) 10 tRNA genes in Gundel; ii) a relatively short (199 aa) zinc-dependent metallopeptidase, formed from a lipoprotein domain and the peptidase domain in Molly, and iii) a YopX protein in Harreka (Figure 5).

### Environmental phage genomes

Five of the 7 proposed new families, five (“Aggregaviridae”, “Forsetiviridae”, “Pompuviridae”, “Winoviridae” and “Flosverisviridae”) had members for which the genomes were assembled from environmental metagenomes (Figure 3, SI file 1 Table 5). We have briefly investigated which bacterial groups are potential hosts for these phages. Most of them gave BLASTN hits with a length between 190 and 5845 bases with bacterial genomes from the *Bacteroidetes* phylum (Figure 3 SI file 1 Tab 4). Furthermore, two of the marine environmental genomes from “Aggregaviridae” and “Flosverisviridae” had hits with viral spacers from CRISPR arrays in *Bacteroidetes* genomes (SI file 1 Table 5). Additionally, several marine contigs from the “Flosverisviridae” contained Bacteroidetes Associated Carbohydrate-binding Often N-terminal (BACON) domains (SI file Table 6). Together, these results suggest that at least part of the environmental phage contigs in the new families are also infecting members of the phylum *Bacteroidetes*.

Integrase encoding genes were found on all environmental phages from the “Forsetiviridae”, and part of the genomes from “Winoviridae” and “Flosverisviridae”. LuxR encoding genes were found in many genomes from “Flosverisviridae”, and one had also a transporter for the auto-inducer 2 (Figure 3). In the “Winoviridae”, all phages encoded Mu-like structural proteins, including the MuD terminase, and two environmental phages also encoded a MuA transposase.

### Flavophages in the environment

#### CRISPR/Cas spacers indicate flavophage presence in the environment

CRISPR/Cas systems were identified in *Polaribacter* sp. HaHaR_3_91 and *Polaribacter* sp. R2A056_3_33 genomes. Spacers from the first strain matched Freya genomes. From the second strains, several spacers matched Danklef genomes, one matched Freya and another Leef (SI file 1 Table 7). This shows that Freya, Danklef and Leef, or their relatives, have infected *Polaribacter* strains in the Helgoland sampling site before 2016, when the host *Polaribacter* strains were isolated. Spacers matching Nekkels, were found in a metagenome of a *Rhodophyta* associated bacterial community (SI file 1 Table 8), showing the presence of Nekkels or its relatives in this habitat.

#### Read mapping for phages and hosts show presence in North Sea waters

To assess the presence of flavophages in the North Sea, we mined by read mapping cellular metagenomes from the 2016 and 2018 spring blooms. We found five of the new flavophages in the cellular metagenomes from the 2018 spring phytoplankton bloom, at three different time points (Table 2). The complete genomes of Freya, Harreka and Ingeline were covered by reads with 100% identity, signifying that exactly these phage isolates were present in the environment. About 85% from Danklef’s genome was covered with reads having 100% identity, indicating that close relatives of this phage (e.g. same species) were present. The genome of Leef was covered only 62% with reads of >70% identity, suggesting that more distant relatives (e.g. genus level) were detected. All phages and their relatives were exclusively found in the >3 μm and >10 μm metagenomes. The most abundant flavophages were Freya and Danklef, reaching 53.8 and 10.3 normalized genome coverage.

**Table 2:** Read mapping results from 2018 metagenomes for isolated flavophages and their hosts.

| Julian day<br>/Greorian<br>date | Fraction | Phage |  | Host |  | Phage/Host<br>** |
| --- | --- | --- | --- | --- | --- | --- |
|  |  | Name | Norm.<br>genome<br>coverage | Name | Norm.<br>genome<br>coverage |  |
| 102 /<br>12.04.2018 | 10 µm | Freya* | 53.8 | *Polaribacter*<br>sp.<br>(HaHaR_3_91<br>and<br>R2A056_3_33) | 0.14 | 384 |
|  |  | Freya relatives | 63.3 |  |  | 454 |
|  |  | Danklef | 10.4 |  |  | 74 |
|  |  | Danklef relatives | 45.8 |  |  | 327 |
|  |  | Ingeline* | 0.7 |  |  |  |
|  |  | Ingeline relatives | 0.8 |  |  |  |
|  |  |  |  | *Olleya* sp.<br>HaHaR_3_96 | 0.05 |  |
| 116 /<br>26.04.2018 | 10 µm |  |  | *Polaribacter*<br>sp.<br>HaHaR_3_91<br>and<br>R2A056_3_33 | 0.06 |  |
|  |  |  |  | *Olleya* sp.<br>HaHaR_3_96 | 0.03 |  |
| 122 /<br>02.05.2018 | 0.2 µm |  |  | *Polaribacter*<br>sp.<br>HaHaR_3_91<br>and | 0.07 |  |
|  |  |  |  | *Olleya* sp.<br>HaHaR_3_96 | 0.05 |  |
| 128 /<br>08.05.2018 | 3 µm | Harreka* | 1.8 |  |  |  |
|  |  | Harreka relatives | 5.7 |  |  |  |
| 142 /<br>22.05.2018 | 10 µm | Freya relatives | 0.04 | *Polaribacter*<br>sp.<br>(HaHaR_3_91,<br>R2A056_3_33<br>and AHE53.11) | 0.46 | 0.09 |
|  |  | Danklef relatives | 0.03 |  |  | 0.07 |
|  |  | Leef relatives | 0.02 |  |  | 0.04 |

Further, we searched for the presence of the five flavophage hosts in the 2018 spring bloom (Tab 2). *Polaribacter* sp. was found in the >10 μm and >0.2 μm fractions, at different time points during the bloom, including in the two same samples where its phages (Freya, Danklef and Leef) and their relatives were detected. At the beginning of the bloom, in April, there were significantly more phages than hosts. The phage/host genome ratio was 384 for Freya, and 74 for Danklef. One and a half months later, only relatives of Freya, Danklef, and Leef were found, the phage/host ratio being lower than 0.1. Discrimination between the different *Polaribacter* strains was not possible, due to their high similarity (>98% ANI). *Olleya* sp. was found in the > 0.2 μm and > 10 μm fractions, at low abundance and dates preceding the detection of its phage, Harreka. *Cellulophaga* sp., the host for Ingeline, was not found (SI file 1 Table 9-13).

## Discussion

We have performed a cultivation-based assessment of the diversity of flavophages potentially modulating *Flavobacteriia*, a key group of heterotrophic bacteria. With our results, we are able to connect phages with their host species and even strain, in contrast to high-throughput viromics-based studies. In addition, these phages were isolated in the ecological context of spring bloom events in two consecutive years. In 2017, we implemented a method of enriching flavophages on six host bacteria. A much larger and more diverse collection of 21 mostly recently isolated *Flavobacteriia* was used in 2018.

Highly diverse flavophages were isolated. Their taxonomic diversity is evident immediately by having representatives in two out of four of the newly delineated viral realms that is in *Duplodnaviria* and *Monodnaviria*. As a point of reference, a viral realm is the equivalent of a domain in the cellular world (60). Furthermore, inside *Duplodnaviria*, the new flavophages fall into 7 new families, four of which have no previously cultivated representatives. At the genus level, out of ten new genera, only “Callevirus” includes previously cultivated phages. The newly delineated families are relevant not only for the marine environment. Besides cultivated flavophages, five of the families also include environmental phages from marine, freshwater, wastewater and soil samples.

Genomic analysis indicates that the new flavophages have various life styles. The presence of integrases in Danklef, Freya, Peternella, Leef and Ingeline, as well as in many of the other phages in their families, points toward the potential to perform a temperate life cycle. However, the absence of recognizable gene markers for lysogeny in the genomes of Harreka, Nekkels, Gundel, Calle, Molly, Colly suggests that most likely these phages are strictly lytic. With respect to genome replication, several strategies can be recognized. First, the genome ending in DTRs of Gundel and the Cos-3’ ends of Leef are indicators for replication through long concatemers (63). Second, the MuA-transposase of Peternella, as well as of other environmental phages in this family, indicate a replicative transposition strategy (64). And third, Omtje, a ssDNA phage, most likely replicates through a ssDNA rolling circle mechanism (65). At the translation level, Calle and Gundel likely increase the protein synthesis efficiency by encoding a high number of tRNAs and even a tmRNA (Table 1), similar to other cultivated phages of the “Pompuviridae” family.

With regard to lysis, flavophages encode various peptidoglycan degrading proteins potentially functioning as endolysins. These genes are usually surrounded by genes coding for proteins with several TMDs, likely holins and anti-holins, with the exception of the endo-beta-N-acetylglucosaminidase of Omtje. Harreka and Nekkels do not encode easily recognizable lysis enzymes. Instead, they encode each a GH19. Usually, this hydrolase family is known for chitin degradation yet also peptidoglycan may be degraded (66). A phage GH19 expressed in *Escherichia coli* caused cellular lysis (67). Furthermore, in Harreka and Nekkels, the vicinity with potential holins, antiholins and spanin, and the peptidoglycan-binding domain in Nekkels, suggest that the GH19 proteins of these two phages likely function as endolysins and degrade bacterial peptidoglycan. It cannot be excluded though that these enzymes have a dual function. Once released extracellularly due to lysis, they could degrade chitin, an abundant polysacharide in the marine environment, produced for example by green-algae or copepods (68). The lysis mechanism in the new dsDNA flavophages likely follows the canonical holin/endolysins model, as suggested by the lack of membrane binding domains in the potential endolysins.

The presence of lyases in the genomes of Leef (pectin lyase), Ingeline (pectin lyase) and Nekkels (pectin and alginate lyases) suggests that their bacterial hosts, *Polaribacter* sp. and *Cellulophaga* sp., are surrounded by polysaccharides. In the marine environment, both alginate and pectin, are produced in large quantities by micro and macro algae (69–71), which serve as substrate for marine *Flavobacteriia*, especially *Polaribacter* (72–74). Bacteria are not only able to degrade these polysaccharides, but also to produce them and form capsules or an extracellular matrix of biofilms. In phages, such polysaccharide degrading enzymes are usually located on the tails, as part of proteins with multiple domains to reach the bacterial cell. Another enzyme potentially degrading proteins in the extracellular matrix is the zinc-dependent metallopeptidase of Molly, which infects a *Maribacter* strain. By depolymerizing the extracellular matrix surrounding the cells, lyases and peptidases help the phage to reach the bacterial membranes for infection or allow the new progeny to escape the cell debris and the extracellular matrix (75, 76). It remains to be proven if phages carrying these enzymes contribute significantly to the degradation of algal excreted polysaccharides, as a byproduct of their quest to infect new bacterial cells.

Previous studies indicate that flavobacteriia can have a surface-associated life style (77). Our results paint a similar picture. For example, we detected phages for *Polaribacter, Cellulophaga* and *Olleya*, as well as the *Polaribacter* and *Olleya* genomes themselves in the particulate fraction of the cellular metagenomes. Therefore, it is likely that these bacteria are associated with particles, protists or zooplankton. Viral spacers matching Nekkels, a *Cellulophaga* phage, with English Channel metagenomes suggest an association with red macro-algae (SI file 1 Table 8). An association with eukaryotes is also supported by the presence of YopX proteins in Harreka, infecting *Olleya*, and in all three “Assiduviridae” phages infecting *Cellulophaga*. Scanning electron microscopy images of *Cellulophaga* sp. HaHaR_3_176 cells show high amounts of extracellular material (SI file 1 Figure 26), indicating the ability for an attached life style.

In the “Flosverisviridae” family, Leef and Ingeline, as well as all other cultivated flavophages and most of the environmental genomes, encode an integrase gene, which is a marker for a temperate life style, and a LuxR gene, which is a quorum sensing dependent transcriptional activator (Figure 3 and SI file 1 Table 6). This suggests that “Flosverisviridae” phages potentially respond to changes in the hosts cell densities, for example by entering the lytic cycle, as recently observed in *Vibrio cholera* (pro)-phages (78, 79). This type of quorum-sensing response would make sense in a habitat in which the host cells can achieve high densities, as for example in association with eukaryotes or other particles.

All of our flavophages, with the exception of Molly and Colly, were actively produced at different times throughout the 2018 phytoplankton bloom, as shown by their isolation by enrichment or direct-plating. Several of these flavophages were also found with 100% read identity covering the whole genome in the cellular metagenomes from the 2018 bloom, indicating not only an intracellular presence, but also higher abundances. Additionally, the genus “Callevirus” is related to the 5^th^ most abundant cluster in the Global Ocean Virome (14). Calle and Nekkels, infecting *Cellulophaga*, although not detected in the cellular metagenomes, were in high abundance (at least 100 PFU ml^-1^) in the viral fraction of our samples, as revealed by their isolation by direct-plating without previous enrichment. This apparent discrepancy between the presence in either the viral or cellular fraction, could be explained either by the phage lysis preceding the metagenome collection, or by their host habitat not being sampled for metagenomics, because *Cellulophaga* are known to grow on macro-algae (80). The presence of Omtje, Peternella and Gundel in enrichments, but not in direct-plating or in the cellular metagenomes, indicate their presence in the environment in low abundances. Further investigations of phage and host habitat are necessary to confirm these findings.

From the five flavophages detected in the 2018 spring bloom metagenomes, Freya, Danklef, Leef and Ingeline have a temperate potential, and Harreka is most likely strictly lytic. The ratio between phage and host normalized read abundance can be used to determine their lytic or temperate state in environmental samples. *Cellulophaga*, the host of Ingeline, was not detected throughout the spring bloom. However, because Ingeline was detected presumably in the intracellular fraction, we can hypothesize that either its host was in a low abundance, or its genome was degraded under the phage influence. Either way, it points toward Ingeline being in a lytic cycle at the time of detection. For Freya, Danklef and their relatives, the high phage to host genome ratios (as high as 454 x) from the 12^th^ of April suggest that these phages were actively replicating and lysing their cells. In contrast, when they reappeared on the 22^nd^ of May, these phages were not only three orders of magnitude less abundant than in April, but also ~ 10 times less abundant then their host. This indicates that in May, only a small proportion of the *Polaribacter* cells were infected with relatives of Freya and Danklef, either in a lytic or a temperate state.

A temporal dynamics in the phage populations is also visible from the isolation results for Freya and Gundel and it is reminiscent of the boom and bust behavior previously observed with T4-like marine phages (81). Furthermore, the read mapping results indicate a change in the dominant phage genotypes, from the same species and even the same strain with Danklef and Freya in April, towards more distant relatives at the level of genus or even family in May. This suggests a selection of resistant *Polaribacter* strains after the April infection and differs from the genotypic changes previously observed for marine phages, which were at the species level (82). Persistence over longer times in the environment was be shown for *Polaribacter* phages by the CRISPR/Cas spacers, and for Omtje and Ingeline, by their isolation both in 2017 and 2018.

The North Sea flavophages isolated in this study have a high taxonomic, genomic and life style diversity. They are a stable and active part of the microbial community in Helgoland waters. With their influence on the dynamics of their host populations by lysis or on other bacterial population by substrate facilitation, they are modulating the carbon cycling in coastal shelf seas. The increase in bacterial numbers, mirrored by the phage numbers and the phage to bacterial cell ratio, indicate that phages actively replicate through the 2018 phytoplankton bloom, matching previous observations of the North Sea microbial community (83). Our read mapping data indicate a complex dynamics, which can now be further investigated with the novel phage-host systems obtained in this study.

## Acknowledgements

The authors would like to thank Carlota Alejandre-Colomo, and Jens Harder for providing several bacterial isolates. For sampling and 16S rRNA analysis we would like to thank Lilly Franzmeier and Mirja Meiners. Many thanks to the Biologische Anstalt Helgoland (BAH), the Aade Crew for providing the infrastructure for the sampling campaign, and Karen Wiltshire for providing the chlorophyll *a* and algae fluorescence data. The authors acknowledge the help in the lab of Jan Brüwer, Jörg Wulf and Sabine Kühn, the funding of the German Research Foundation (DFG) project FOR2406 - “Proteogenomic of Marine Polysaccaride Utilisation (POMPU)” project and the Max Planck Society. Part of the bioinformatics analyses were performed on the High Performance Computing Cluster CARL, located at the University of Oldenburg (Germany) and funded by the DFG through its Major Research Instrumentation Programme (INST 184/157-1 FUGG) and the Ministry of Science and Culture (MWK) of the Lower Saxony State. EMA was funded by the Biotechnology and Biological Sciences Research Council (BBSRC); this research was funded by the BBSRC Institute Strategic Programme Gut Microbes and Health BB/R012490/1.

## Conflict of Interest

E.M.A. is the current Chair of the Bacterial and Archaeal Viruses Subcommittee of the International Committee on Taxonomy of Viruses, and member of its Executive Committee. The other authors declare no conflict of interest.

Supplementary information is available at ISME Journal’s website.

## Notes

### Competing Interest Statement

The authors have declared no competing interest.

