## Supplementary material for "Highly diverse flavobacterial phages as mortality factor during North Sea spring blooms": SI_Fig_5

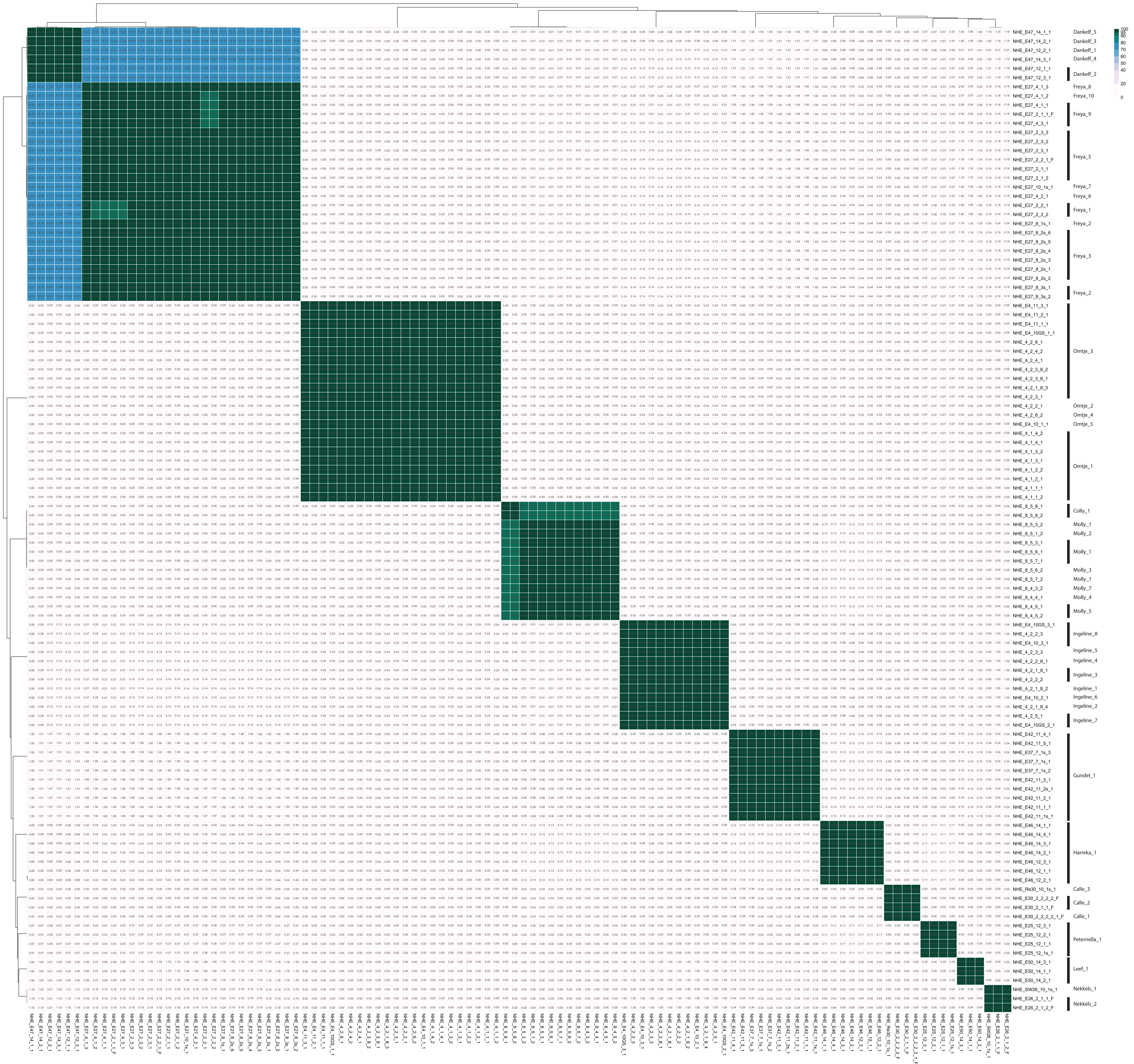


**Figure 5:** Intergenomic nucleotide similarity matrix of all obtained isolates. Phage strains are indicated on the right side.
