## Supplementary figures and images for "Highly diverse flavobacterial phages as mortality factor during North Sea spring blooms"

### SI_Fig_7

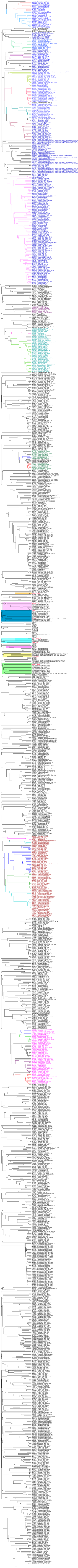
