## Supplementary material for "Highly diverse flavobacterial phages as mortality factor during North Sea spring blooms": SI_Fig_9

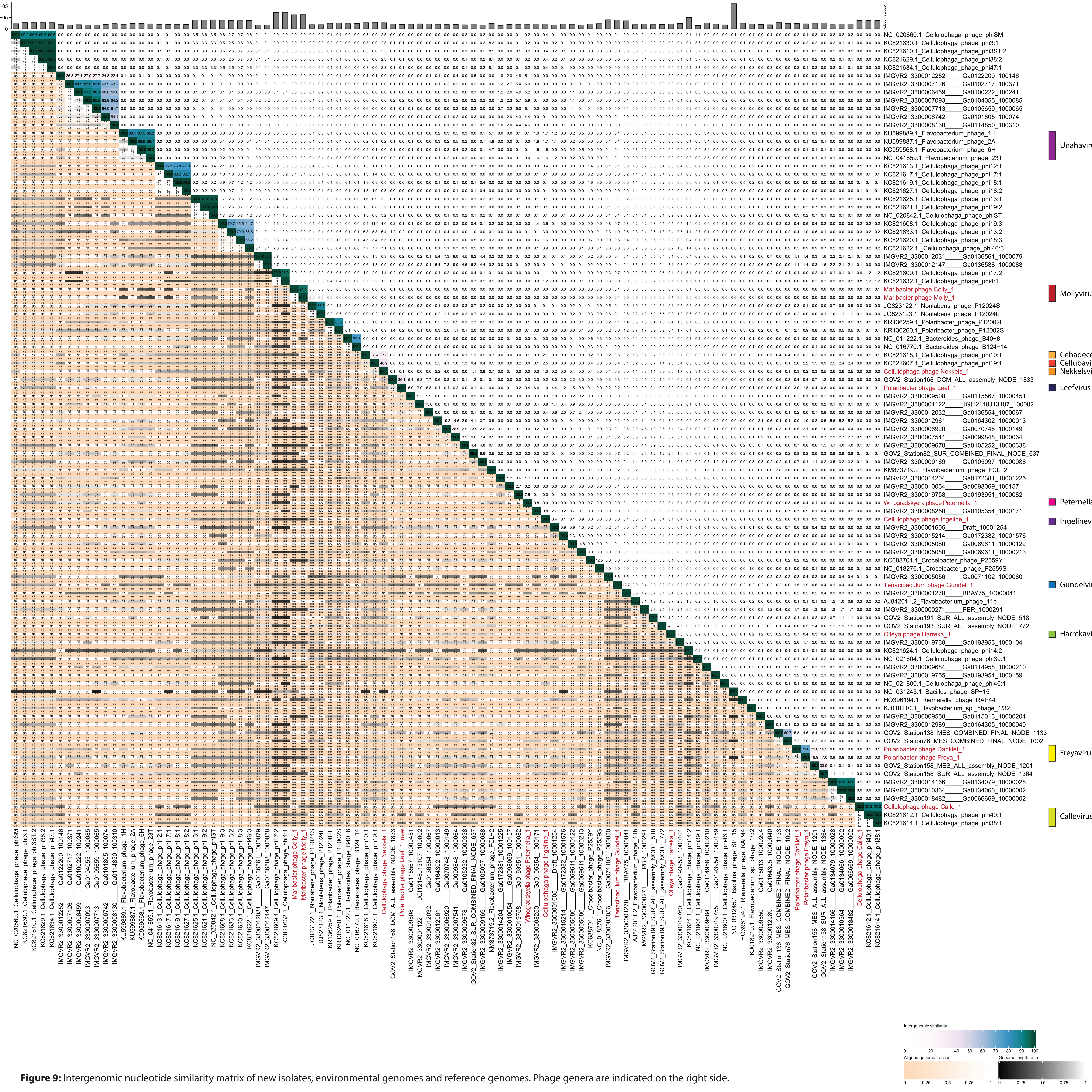


**Figure 9:** Intergenomic nucleotide similarity matrix of new isolates, environmental genomes and reference genomes. Phage genera are indicated on the right side.
