## Supplementary material for "Highly diverse flavobacterial phages as mortality factor during North Sea spring blooms": SI_Fig_23

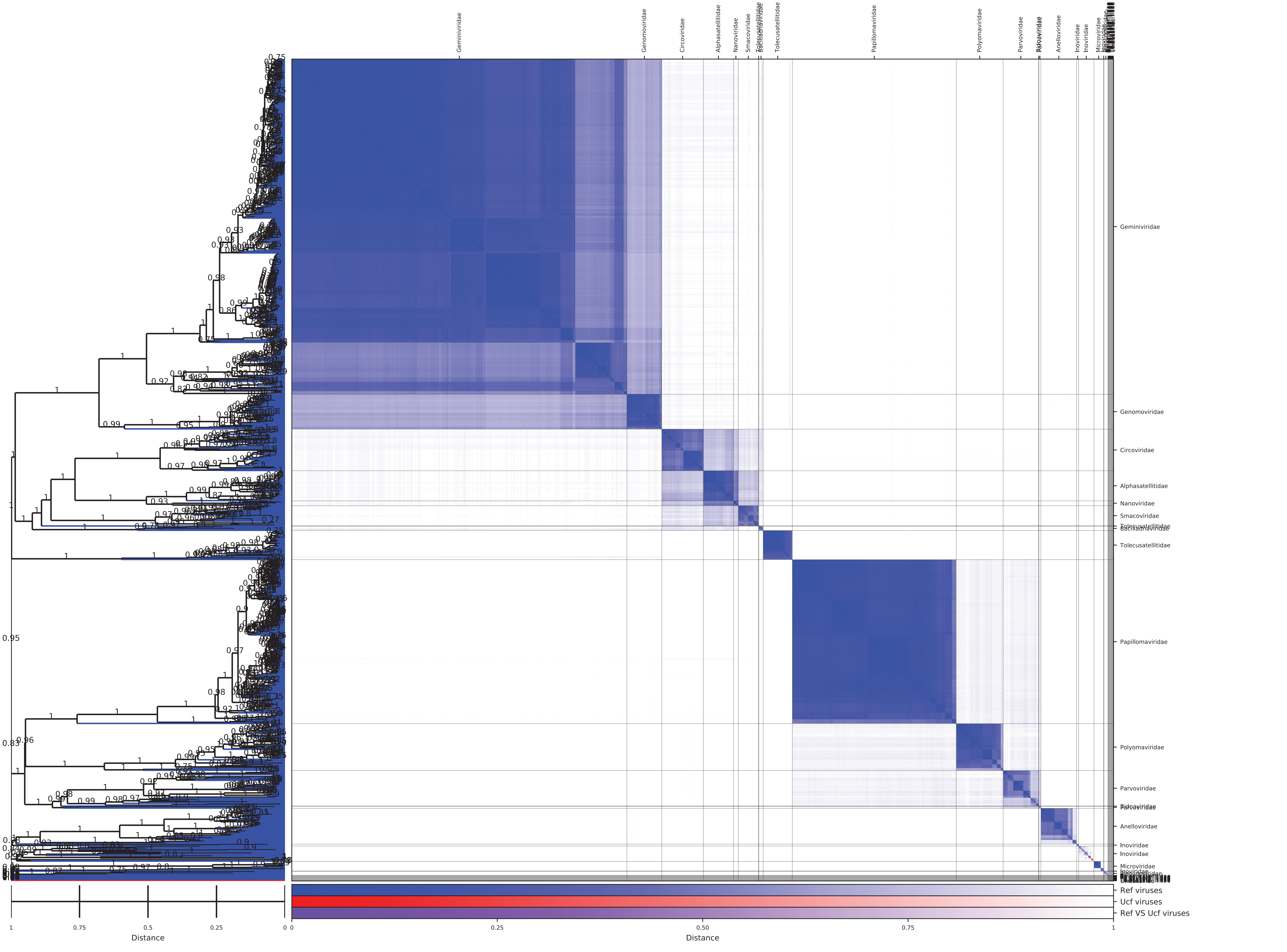


**Figure 23:** Heatmap from GRAViTy with new ssDNA phage isolate, ssDNA Cellulophaga phages and the Baltimore Group II - ssDNA viruses + Papillomaviridae and Polyomaviridae (VMRv34) database. Red squares indicate the position in the heat map of the "Obsuriviridae".
