## Supplementary material for "Highly diverse flavobacterial phages as mortality factor during North Sea spring blooms": SI_file_1

### Supplementary Information_file_1

#### Materials and Methods

#### Environmental samples

##### Chlorophyll measurements

Chlorophyll *a,* green algae, and diatoms were measured via fluorescence using an algal group analyser (bbe moldaenke, Kiel-Kronshagen, Germany).

##### Total bacterial counts and *Bacteroidetes* counts

Samples for total cell counts were fixed with a final concentration of 1% formaldehyde for 1 h at room temperature. Aliquots of 10 ml were filtered on a 0.2 µm polycarbonate filters (Merck Millipore, Burlington, USA), and stained with 4',6-diamidino-2-phenylindole (DAPI, 1µg/ml) for 20 min at room temperature and shortly washed with MilliQ water and 80% ethanol. Dried filters were automatically counted with the Zeiss Axio Imager.Z2 (Carl Zeiss MicroImaging GmbH, Jena, Germany) and quantified with the modified ACMEtool3.0 (Zeder, M. 2005-2010, Software for Biology, http://www.technobiology.ch) after (1).

On the same filter, catalysed reporter deposition – fluorescence *in situ* hybridization (CARD-FISH) was performed with the CF319a probe (TGGTCCGTGTCTCAGTAC) (2) according to (3) with modifications. Hybridization was done with horseradish peroxidase (HRP)-labeled oligonucleotide probes at 35% formamide concentration.

##### Determination of virus-like particle (VLP) numbers using epifluorescence microscopy

Epifluorescence microscopy of fluorescently stained samples was used to count virus like particles (VLPs). For this purpose 0.5 ml of the of 0.2 µm filtrate were filtered through a 0.02 µm pore size Anodisc filter (GE Healthcare Life Sciences, Maidstone, UK), and stained with SYBR Gold (Invitrogen, Carlsbad, USA) (25x final concentration) for 14 min at room temperature. At least 10 fields of view were counted (4, 5) using the Zeiss AxioImager.D2 (Carl Zeiss MicroImaging GmbH).

##### Determination of phage numbers using transmission electron microscopy (TEM)

Phages were also counted by transmission electron microscopy (TEM) with the JEM-2100 (JEOL, Tokyo, Japan). Samples were filtered through a 3 µm filter to remove big aggregates, fixed with a final concentration of 2% glutaraldehyde, and stored at 4°C. For electron microscope grid preparation, 10 ml were centrifuged in a SW40TI (Beckman Coulter, Brea, USA) at 20,000 rpm for 1 hour at 4°C on two carbon coated copper grids with a polyvinyl butyral support film. Samples were negative stained for 1 min with 0.5% uranyl acetate and images of 2-3 meshes were taken, with a TemCam F416 (TVIPS, Gauting, Germany) camera, a beam of 44 µA and

80 kV, and 12,000x magnification. A detailed description of the method can be found in Brum et al. 2005 (6).

#### Phage isolation

##### Media preparation

*2216 Difco Marine Broth (MB)*

Medium was prepared following the manufacturer's recommendation. In short, 37.4 g Difco Marine Broth (BD Biosciences, San Jose, CA, USA) were dissolved in 1 l MilliQ water and autoclaved at 121°C for 20 min. This medium was used in 2017.

*Marine Broth (MB) from native sea water*

North Sea surface water was filtered through 0.2 µm membrane (Merck Millipore) diluted with MilliQ water to a final concentration of 75%. Both, 1 g $l^{-1}$ Bacto Yeast Extract (BD Biosciences) and 5 g $l^{-1}$ Bacto Tryptone (BD Biosciences) were dissolved, the pH was adjusted to 7.6, and the medium was autoclaved at 121°C for 20 min. This medium was used in 2018, because the bacteria grew better in this medium.

*Agar for Plaque Assay and Spot Assay*

For the bottom agar, 15 g $l^{-1}$ Bacto Agar (BD Biosciences) was added to the above described media before autoclaving, but after pH adjustment. 10 ml were poured in 92 mm vented Petri dish (Sarstedt) and dried for one day at room temperature. Plates were stored at 4°C and again dried for at least one hour before usage.

For the top agar, 6 g $l^{-1}$ Bacto Agar (BD Biosciences) was added to the above described media before autoclaving, but after pH adjustment. 3 ml were sterile poured into 15 ml tubes (Sarstedt). Tubes were stored at 4°C and melted in boiling water prior usage. In order to avoid killing the bacterium it was cooled down to 42°C in a water bath.

.

##### Spot Assay

After transferring 500 µl of densely grown host culture to the molten top agar (42°C), the mixture is poured on top of the bottom agar and spread by swirling. After solidification 10 µl of phage stock was transferred to the upper third of the plate. If several samples were tested on a single plate, those drops were in a horizontal line. Plates were tilted in order to start a slow flow of the drops which was stopped before a drop was reaching the rim of the Petri dish. Plates were incubated at 18°C for one to two days, depending on the host growth.

##### Plaque Assay

To obtain single plaques, a decadal dilution series of the phage stock was performed with saline magnesium (SM) buffer (5.8 g $l^{-1}$ NaCl, 2.0 g $l^{-1}$ $MgSO_4$ * 7 $H_2O$, and 50 mM Tris-HCl (pH 7.5), dissolved in MilliQ water, and autoclaved). For each dilution a single plate was used following the same procedure. 100 µl of phage liquid was transferred to the middle of the bottom agar plate. 500 µl of densely grown host culture was transferred to molten top agar. This mixture was poured on the bottom agar and spread to cover the whole plate. After solidification, plates were incubated at 18°C for one to two days, depending on the host growth.

#### Phage isolation from enrichments and direct-plating

Phages were enriched by mixing 740 ml seawater consecutively pre-filtered through 10 µm, 3 µm, and 0.2 µm pore-size polycarbonate filters (Merck Millipore, Burlington, USA) in 1 L glass bottles (Schott AG, Mainz, Germany) with 27 ml 10x MB medium (7) in a final concentration of 0.3 x 2216 marine both medium (see above). To this mix 27 ml of flavobacterial culture was added pre-grown at 18°C to logarithmic phase MB medium. The enrichments were incubated for four days at 18°C with 100 rpm shaking. In total, 23 bacterial strains were tested (6 in 2017, 21 in 2018) (Tab.1). Afterwards, the enriched phage fraction was obtained by filtering 10 ml through a 0.2 µm syringe filter (Merck Millipore). Successful enrichments were detected by a spot test (8) with the host used for enrichment. If clearing zones were obtained, indicating phage lysis, phage dilutions were plated. Three plaques each were picked and transferred three timesbefore a phage stock was prepared.

#### Phage stock preparation

At least three single plaques were picked from the plates of the plaque assay with a 1 µl plastic inoculation tube. Those plaques were dissolved in modified SM buffer, diluted, and plated again. This procedure was repeated three times to ensure the purity of the final phage stock. With a fourth dilution series the stock was prepared by adding 10 ml SM buffer to a plate with confluent lysis (9). After 1 h at room temperature, the liquid was filtered through a 0.2 µm syringe filter (Merck Millipore) and stored in glass tubes at 4°C

#### Host range determination

For host range determination all bacterial isolates used for the enrichment cultures were tested with each obtained phage group. An agar overlay was performed and 10 µl of phage stock were spotted on the plate (see spot test). Four to five stocks were tested per plate. If lysis was observed an additional overlay was done with only one phage stock per plate. Lysis showing combinations were tested again with a plaque assay.

#### Determination of phage morphology using TEM

Phage stocks obtained from lysates were fixed with glutaraldehyde at a final concentration of 0.5% (2.5% glutaraldehyde, 2.5 mM $MgCl_2$, 50 mM KCl, 50 mM cacodylic acid, pH 7.2). Approximately 10 µl of fixed phages were transferred onto glow discharged carbon coated copper grids. After 2 min the solution was removed from the grids with filter paper, grids were washed for 10 sec with deionized $H_2O$ and then negative stained with 0.5% uranyl acetate for 1 min. The samples were visualized with a Zeiss EM900 TEM (Zeiss, Oberkochen, Germany). TEM negatives were digitalized with an Epson Perfection V700 Photo scanner.

#### Determination of phage genomes

##### Phage DNA extraction

First, free nucleic acids were digested with DNase 1 (Ambion, 0.004 U/µl) and RNase 1 (Ambion, 0.1 U/µl). Then, the capsid was opened with Proteinase K (0.05 U/ml) and SDS (0.5%). Finally, the phage DNA was loaded on Wizard columns (Promega, Madison, USA) by mixing 1 ml of phage lysate with the Wizard resin. A vacuum manifold was used for the extraction. DNA was eluted in TE Buffer (after (10)).

##### Phage genome sequencing

Both types of phages were sequenced with the Illumina HiSeq3000 (paired-end read 2 x 150 bp). For potentially ssDNA phages and Maribacter phages a ChIPSeq library was prepared with the DNA SMART ChIPSeq Kit (Takara Bio Europe S.A., Saint-Germain-en-Laye, France) and 12-18 PCR cycles. For dsDNA phages a DNA FS library was prepared using the DNA Ultra II kit (New England Biolabs GmbH, Frankfurt am Main, Germany) with fragmentation by S2 tubes (Covaris, Woburn, MA, USA) or with chemical fragmentation for 9 min and 3-12 PCR cycles.

##### Phage genome assembly

The raw reads were quality trimmed with BBDuk (v35.14, sourceforge.net/projects/bbmap/), using the parameters “qtrim=rl trimq=20, maq=20 minlen=30 ordered t=8". The cleaned reads were then assembled both with SPAdes (v3.13.0, (11)) and Tadpole (v35.14, sourceforge.net/projects/bbmap/). The parameters for Tadpole were "k=50 t=8". For the first batch of genomes the parameters for SPAdes were "-k 35,55,75 --sc". In a second batch the parameters for SPAdes were “-k 31, 41, 51, 61, 71, 81, 91, 101, 111, 121, 127 -m 500”. If more than one contig was obtained, a normalization of the reads with BBNorm (v35.14, sourceforge.net/projects/bbmap/) was performed with the parameters "target=100 min=5", followed by an assembly as above. BBDuk, Tadpole and BBNorm are part of the BBTools package (https://jgi.doe.gov/data-and-tools/bbtools/).

##### Determination of phage genome ends

Bandage (12) was used to assess the quality of the assembly. If the phage genome had several nodes, or if the genome was linear, primers were designed and amplified fragments were sequenced using the Sanger technology (see below). In order to validate the genome size of the assembly, Pulsed Field Gel Electrophoresis was performed with agarose plugs containing phage stock solution (see below).

##### Gap closure of phage genomes

Primers were designed with SnapGene (GSL Biotech LLC), which covered the region. The amplified DNA was sequenced after Sanger with a 3130xl Genetic Analyzer (ABI PRISM), manually trimmed with FinchTV (v1.4.0, Geospiza Inc.) and mapped on the genome with the Geneious mapper.

#### Determination of phage genome size by Pulsed Field Gel Electrophoresis (PFGE)

For plug preparation 50 µl of 2% low melting agarose (invitrogen, UltraPure™ LMP Agarose) were mixed with 50 µl of phage stock solution and transferred into a plug mold (BIO-RAD, CHEF® Mapper XA System). Plugs were incubated in modified SM buffer with sodium dodecyl sulphate (SDS) (0.5%) and Proteinase K (Macherey-Nagel, 0.26 U/ml) at 56°C for 24 hours. Plugs were washed two times in TE Buffer (10 mM Tris-HCl pH 8, 0.1 mM EDTA) at room temperature for 24 hours. Samples were run in a 1% Pulsed Field Gel (BIO-RAD, Pulsed Field Certified Agarose) in 0.5 x TBE Buffer (0.045 M Tris, 0.045 Boric Acid, 0.001M EDTA) with a CHEF-DR® III System and cooling module (BIO-RAD). The initial switch time was 0.3 sec and was increased by a linear ramp to 11.5 sec. The run time was 19.5 hours with 6 V/cm, an included angle of 120°, in 0.5 x TBE buffer in a 14°C cooled system. Gels were with ethidium bromide and images were taken with an Intas UV-System Gel iX Box (13).

#### Phage genome enzymatic digestion

DNA of Omtje_3 was extracted with the Wizard DNA extraction resin (1,08 µg/µl), and digested at room temperature for 40 min with the Exonuclease III (Thermo Fisher, final concentration 25 U/µl) and DNase I (Ambion, final concentration 0.5 U/µl). Digestion products were analysed with a fragment analyzer system 5200 (Agilent Technologies, Santa Clara, CA, USA) with the HS Genomic DNA Kit (Agilent Technologies).

#### Retrieval of related phage genomes

First, the datasets were prepared. For this, contigs smaller than 10 kb were removed. Open reading frames (ORFs) were predicted with MetaGeneAnnotator (14). The ORFs were then translated into proteins with a custom R script using the seqinr package and the translation code 11. All predicted proteins were pooled into a single BLAST database, named here ENV_DB, using makeblastdb tool from the BLAST+ 2.6.0 package (15). Second, flavophage related contigs were found as follows. The proteins of the isolated flavobacterial phages were used as query for a BLASTP search against ENV_DB, with the parameters "-evalue 0.001 -max_target_seqs 10000". Protein hits with a bitscore lower than 50 were removed. All contigs from ENV_DB with at least four protein hits with a flavophage query were selected. And third, all flavophage genomes and the above selected environmental genomes were clustered using vConTact2 (16, 17) with the standard parameters and cultivated phages of the ProkaryoticViralRefSeq85-ICTV. All environmental phage contigs and the reference genomes that formed one cluster with the flavophage isolates were selected for further analysis. Only environmental contigs with a length >80% compared to their related flavophage isolate were kept.

#### Phage genome annotation

First, the ORFs were predicted using MetaGeneAnnotator and translated using a custom R script, with the seqinr package and translation code 11. Second, tRNAs and tmRNAs where predicted with using tRNAscan 2.0 (parameters "-q -B –D") (18, 19) and Aragorn v1.2.38 (parameters "-m -fo - gcbact –fon") (20), integrated in a custom R script. The InterProScan (21) plugin from Geneious v 11.1.5 (http://www.geneious.com, (22) was used to predict the cellular localization of the protein domains and presence of signal peptides.

#### Isolation and cultivation of particle-associated heterotrophic bacterial strains

Bacterial strains were isolated spring 2017 from particle-enriched surface-enriched seawater samples on 2216 plates and an artificial seawater plate medium with 2 gL-1 laminarin as major carbon source at 12°C in the dark (modified from (23)). Further details are shown in Table 14.

#### Host imaging by Scanning Electron Microscopy (SEM)

Bacterial colonies grown on agar plates were excised together with the underlying agar and fixed for 12-24 hours at 4°C with 6.25% glutaraldehyde (Merck, Darmstadt, Germany), 50 mM Soerensen phosphate buffer, pH 7.4. Samples were then washed three times with 66 mM Soerensen phosphate buffer, pH 7.4.

Samples were stepwise dehydrated with acetone, critical point dried (critical point dryer: BAL-TEC CPD 030) and metal coated (sputter coater BAL-TEC SCD 005) with gold-palladium. Specimens were inspected with a field emission scanning electron microscope (JEOL JSM-7500F) at 5 kV using a detector for secondary electrons (LEI detector).

#### Sequence analysis of host 16S rRNA genes

Host strains were grown on 2216 medium plates. Single colonies were picked and transferred to 20 µl PCR grade water. After three freeze-thaw cycles a PCR was performed with GM3_F and GM4_R as primers to amplify the 16S rRNA gene. Products were purified and the sequencing reaction with the primers GM3_4, GM4_R, GM1_F, and GM1_R was performed. After purification, the products were sequenced with a Sanger machine 3130xl Genetic Analyzer (ABI PRISM), manually trimmed with FinchTV (v1.4.0, Geospiza Inc.), and assembled with Geneious.

#### Host genome sequencing

Isolates from phage – host systems were additionally genome sequenced with a Sequel I (Pacific Biosciences, Menlo Park, USA) using 16mer barcodes. The library was prepared according to 20 kb template preparation for Sequel Systems using the SMRT bell Template Prep Kit 1.0 SPv3 (Pacific Biosciences). After using the Covaris g-tube fragmentation for 9 kb fragments (Woburn, USA), a size selection on a Blue Pippin (Sage Science, Beverly, USA) was done to enrich for fragments above 8 kb in a 0.75% cassette.

#### Host genome assembly and analysis

Reads were assembled with the HGAP4 (24) or CANU (25) assembler. Assemblies were manually edited to remove duplicated overlapping regions. The 16S rRNA genes were retrieved using the MiGA online platform (26). The average nucleotide identity (ANI) was calculated with the enveomics command line package (27).

### Polaribacter Phages

#### “Polaribacter virus Freya" species

The “Polaribacter virus Freya” was isolated very early in the bloom with its host *Polaribacter* sp. HaHaR_3_91 (DSM111048). The ten strains obtained share a similarity of 94.68-99.05%. All genomes are circular, ranging from 43,978 up to 48,920 bp and had a GC content of 28.9%. The diameter of the capsid is 53.9 ± 4.7 nm, the tail is 151.0 ± 8.2 nm long, and 13.1 ± 2.0 nm wide. The morphology was siphoviral-like. Genomes varied in size by 11 genes, which were all annotate as hypothetical proteins. Genes encoding capsid, tail, tape measure, terminase, and portal proteins were annotated. Closely related sequences are also from the Norwegian Sea and are 16.2% and 13.7% similar.

#### “Polaribacter virus Danklef” species

“Polaribacter virus Danklef” with its five strains Danklef_1-5 (similarity > 95.6%) were isolated late in the 2018 bloom, infecting *Polaribacter* sp. R2A056_3_33 (DSM111047). The capsid of Danklef was 46.1 ± 2.2 nm in diameter, the tail was 157.4 ± 4.6 nm long, and 12.1 ± 1.8 nm wide. Danklef had a siphoviral morphology. Its circular genome had a size of 47,177-47,426 bp with a GC content of 28.9%. Danklef’s major capsid protein was HK97-like. It had genes coding for a portal, tape measure, terminase, integrase, and N-acetylmuramidase protein. Interestingly, Danklef had a ferric uptake regulator family related gene. Danklef’s closest environmental relatives are coming from the Norwegian Sea and are 16.8 and 15.14% similar.

Freya and Danklef were closely related (71.38%) and belong to the same genus. Comparing the two phages, Freya had a peptidase and a DNA replication protein, whereas Danklef had two endonucleases, a methylase, and an endolysin, which is not found in Freya.

#### “Polaribacter virus Leef” species

“Polaribacter virus Leef” infected *Polaribacter* sp. AHE13PA (DSM111061). Leef was isolated at the peak of the bacterial bloom. It had a capsid size of 49.2 ± 3.6 nm, a tail length of 138.7 ± 9.6 nm, and tail width of 11.1 ± 2.0 nm. The appearance was siphoviral. Its circular genome had Cos 3’ ends, a size of 37,547 bp and a GC content of 29.7%. Leef encoded proteins related to the HK97 Phage, a LuxR, a BACON (Bacteroidetes-Associated Carbohydrate-binding Often N-terminal) domain, a pectin lyase, and two integrases. Genes encoding capsid, tail tape measure, and neck proteins were annotated along with a terminase and a portal protein. An N-acetylmuramidase was detected in Leef, surrounded by transmembrane domains (TMDs) containing proteins. The closest relative to Leef was node 1833 from the GOV2 dataset, which was sampled in the Barents Sea. They were 30.4% similar.

### Cellulophaga Phages

#### “Cellulophaga virus Omtje” species

With *Cellulophaga* sp. HaHaR_3_176 (DSM111152) a set of five closely related phages were isolated, to which we refer as Omtje_1-5 (sequence identity > 99.9%). They are all belonging to the species “Cellulophaga virus Omtje”. Only Omtje_3 was isolated in 2017 and 2018. Omtje had a capsid diameter of 52.3 ± 4.6 nm, and lack a tail. Thin sections of the phages show a potential lipid layer inside the capsid. DNA digestion revealed that they are ssDNA viruses. Their small circular genome of 6,558 bp with 31.2% GC content also suggests that they are belonging to the tail-less ssDNA phages. ORF prediction revealed 13 genes, which mostly encoded for structural proteins. In addition, a replication initiation factor and a lysis protein (N-aceylmuramoyl-L-alanine-amidase) were identified. Omtje were 57.6% similar to the *Cellulophaga phage phi12:2* (NC_021797.1), which is a ssDNA microviridal phage isolated from the Baltic Sea in 2000 (28, 29).

#### “Cellulophaga virus Ingeline” species

The “Cellulophaga virus Ingeline” infected *Cellulophaga* sp. HaHaR_3_176 (DSM111152). This phage group is very diverse, as indicated by eight isolates, Ingeline 1-8, of a high nucleotide similarity above 99.9%. Ingeline_7 and Ingeline_8 were isolated in 2017 and 2018. Ingeline had a capsid diameter of 59.0 ± 5.3 nm and a tail, which is 132.9 ± 19.3 nm long, and 11.2 ± 1.7 nm wide. It had a circular genome ranging between 42,624 and 42,797 bp and a GC content of 32.2%. We annotated genes encoding a capsid, tail, tail tape measure, adaptor, portal, and a potential spanin protein. The morphology observed by TEM was that of a Siphovirus. Interestingly, the genome also encodes a LuxR gene and a BACON domain-containing protein. Its closest environmental relative is Ga0105354_1000171 from the Norwegian Sea with 12.4% similarity. Although Ingeline is lytic to its original *Cellulophaga* host, it contains two integrases, indicating the potential for lysogeny.

#### “Cellulophaga virus Calle” species

With *Cellulophaga* sp. HaHa_2_95 (DSM111037) the podoviral “Cellulophaga virus Calle” was isolated. Three strains Calle_1-3, which were 99.95-99.98% similar, belong to this species. This species was present throughout the 2018 phytoplankton bloom. The capsid had a diameter of 60.3 ± 3.0 nm. The tail is 23.0 ± 5.5 nm long, and 13.5 ± 2.4 nm wide. The circular genome length of the strains ranged from 72,979 to 72,980 bp and the GC content was 38.1%. The genome encoded a capsid, tail, and DNA polymerase protein. In addition, it contained 20 tRNAs and a tRNA-splicing ligase RtcB. This phage is also encoded a tmRNA. Both types of RNA are suggesting a more efficient phage replication and might increase the host range. Indeed, Calle is able to infect another *Cellulophaga* strain (HaHa_2_1). Two chaperonin proteins, which are associated with the GroEL system were also encoded in the genome. Its closest relative was the *Cellulophaga phage phi38:1* (KC821614.1) with 92.2% similarity, which was isolated in the Baltic Sea in 2005 (28) with a closely related host (99.5% 16S rRNA sequence identity and 94.2% ANI). Phi38:1 and Calle belong to an abundant cluster of marine phages (30).

#### “Cellulophaga virus Nekkels” species

The “Cellulophaga virus Nekkels” was isolated by direct plating of sea surface water with the host *Cellulophaga* sp. HaHa_2_1 (DSM111038). Spot tests indicated that this phage group is present during most of the sampling period. The second strain, Nekkels_2, had a nucleotide similarity of 97.1%. Nekkels had a capsid diameter of 54.8 ± 5.0 nm, its tail was 141.0 ± 7.9 nm long, and 13.0 ± 1.5 nm wide. The two genomes varied in length. Nekkels_2 encoded in its 54,332 bp genome two ORFs more than Nekkels_1 (53,385 bp). Both circular genomes have a 31.5% GC content. The genomes encoded a major capsid, tail tape measure, neck, terminase, pectate lyase, a lysozyme (GH19), and a potential spanin protein. Additional genes were the acyl carrier protein and a Yersinia outer protein X (YopX). Although Nekkels had a 40.5% nucleotide similarity with the *Cellulophaga phage phi19:1* (KC821607.1), which is a podovirus. However, Nekkels had a long tail and was morphological a siphovirus. Nekkels was able to infect another flavobacterial host: AHE13PA, a *Polaribacter* sp., but with a lower efficiency.

### Other flavophages

#### "Olleya virus Harreka" species

The "Olleya virus Harreka" was isolated during the bacterial bloom in 2018 with the host *Olleya* sp. HaHaR_3_96 (DSM). The capsid diameter was 44.3 ± 3.6 nm, the tail was 123.8 ± 8.0 nm long and 14.3 ± 2.0 nm wide. Harreka had a myoviral morphology. In total seven stocks were obtained and all of them revealed the 100% identical 43,175 bp circular genome with a GC content of 32%. Genes for capsid, tail, portal, replication proteins, GH19 and YopX were identified. Harreka was able to infect the two closely related *Tenacibaculum* strains AHE14PA and AHE15PA.

#### "Winogradskyella virus Peternella" species

From *Winogradskyella* sp. HaHa_3_26 (DSM111041) "Winogradskyella virus Peternella" was isolated once in 2018 during the bacterial bloom. It has a capsid diameter of 52.3 ± 4.2 nm, a tail length of 105.8 ± 7.4 nm, and a tail width of 16.4 ± 2.4 nm. The morphology was myoviral. From four phage stocks, the same 39,649 bp long linear genome with 35.3% GC content was retrieved. This phage belongs to the Mu-like phages, due to its overlapping reads with the host genome and the typical genes like the MuA transposase and several structural proteins like the capsid, tail, tail fiber, baseplate, neck. This phage also encodes a portal, a holing, and a L-Alanine-D-glutamine-peptidase protein.

#### "Maribacter virus Molly" species

Using the type strain *Maribacter forsetii* T (DSM 18668) the "Maribacter virus Molly" was isolated from two time points in 2017. Six highly similar 99.56 to 99.99% strains (Molly1-7) were obtained. The phage has a capsid diameter of 74.9 ± 3.6 nm, a tail length of 101.5 ± 6.3 nm, and a tail width of 18.1 ± 1.6 nm. The morphology was myoviral-like. The circular genome had 124,169 to 124,898 bp with a GC% of 36.2. This phage group was very difficult to sequence, which might be due to a high degree of DNA modifications indicated by the respective genes. Furthermore, genes encoding a baseplate, tail fiber, tail, tail sheath, tape measure, neck, portal, major capsid protein, and a DNA polymerase I were identified. Additionally, it had a ribonucleotide reductase with two subunits A and B and a relatively short (199 aa) zinc-dependent metallopeptidase, formed from a lipoprotein domain and the peptidase domain.

#### “Maribacter virus Colly” species

“Maribacter virus Colly” is 94% similar to “Maribacter virus Molly”. Following the ICTV guidelines it is a different species in the genus *Mollyvirus*. Colly had the same functional genes as Molly. The difference is due to genes encoding hypothetical genes.

#### “Tenacibaculum virus Gundel” species

The “Tenacibaculum virus Gunde”l was isolated twice, before and after the phytoplankton peak in 2018, with AHE14PA (DSM111040) and AHE15PA (DSM111039). It had a podoviral morphology with a capsid diameter of 60.5 ± 5.2 nm and a tail length of 22.7 ± 3.2 nm. Ten isolates were obtained, all having a genome size of 78,511 bp and a GC content of 30.4%. The circular genome had short direct terminal repeats (DTRs) at the ends. Gundel can only infect its two hosts of isolation, AHE14PA and AHE15PA, which had 99.87% similar 16S rRNA and an ANI of 99.99%. Gundel had genes coding for tail fiber, portal, L-alanine-D-glutamine-peptidase, and a phage antirepressor protein. In addition, Gundel had 10 tRNAs.

### Tables and Figures

#### Figures

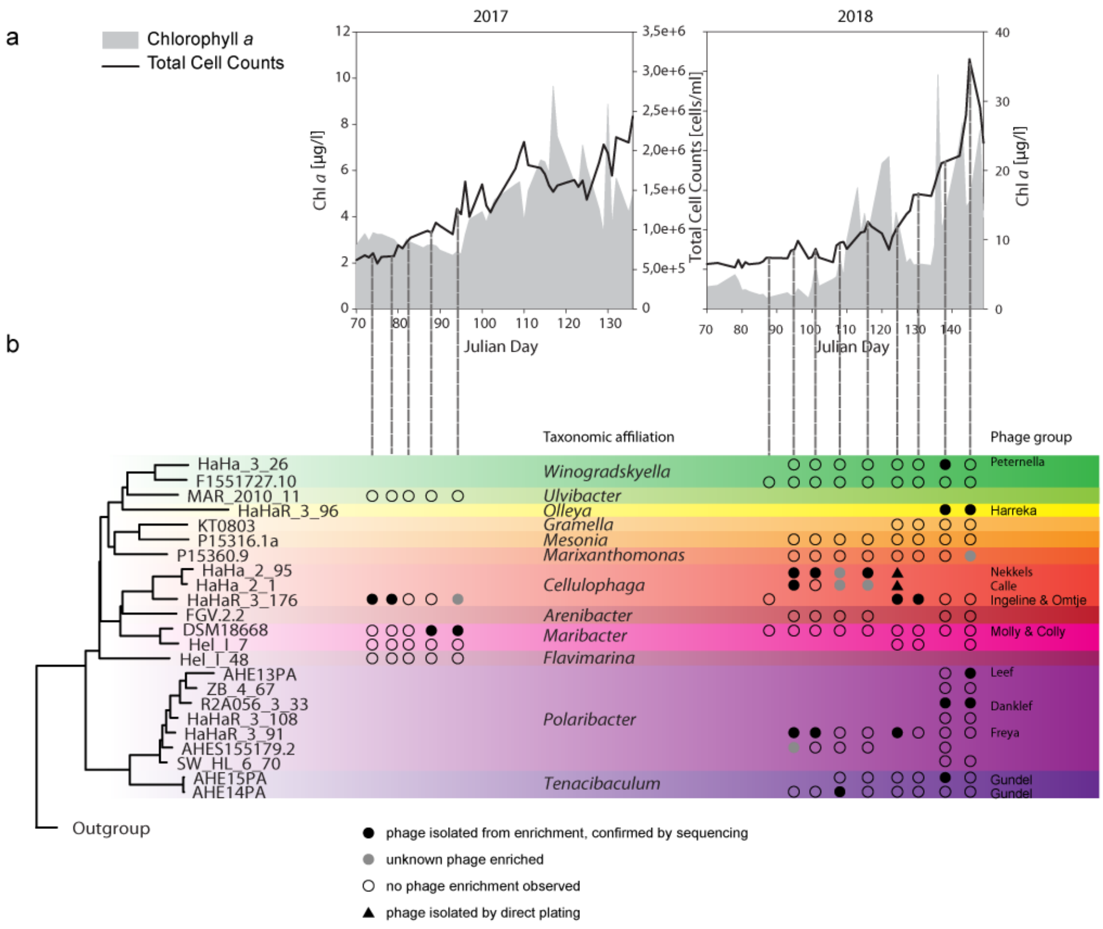


**Figure 1:** Chlorophyll *a* concentration and total cell counts during the sampling and phage enumeration with TEM and epifluorescence microscopy (a). Neighbor – joining tree with bacterial isolates and type strains on the basis of 16S rRNA and the isolation success of the corresponding phages (b).

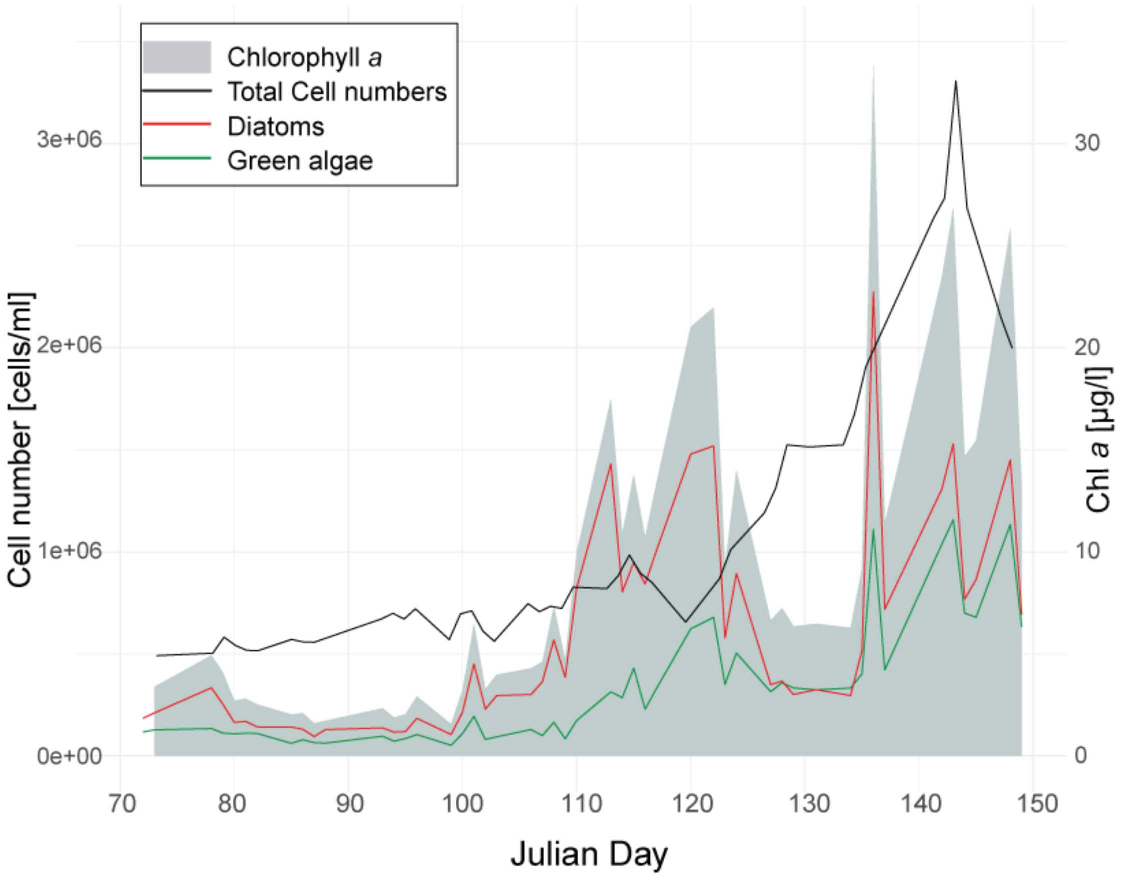


**Figure 2:** Pigment concentration of green algae and diatoms measured by fluorescence over the course of the bloom 2018.

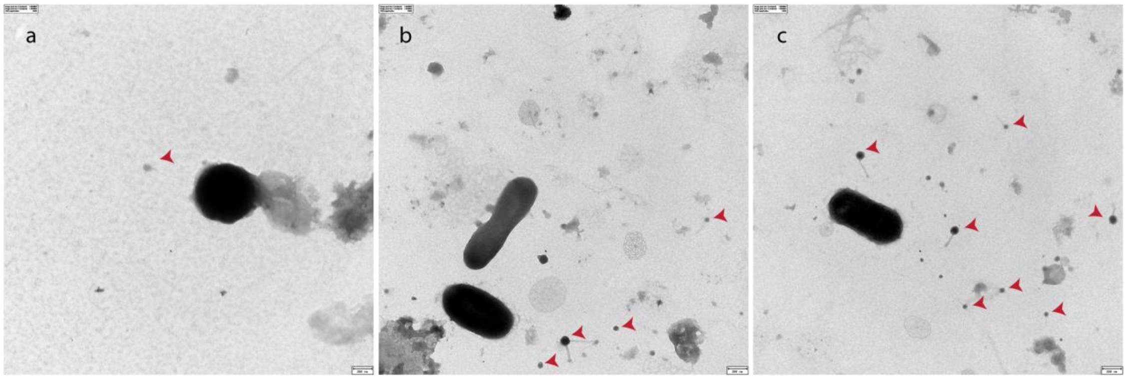


**Figure 3:** Example images of TEM virus counts from Julian Day 102 (a), 128 (b), and 144 (c). Phage particles are marked by arrowheads.

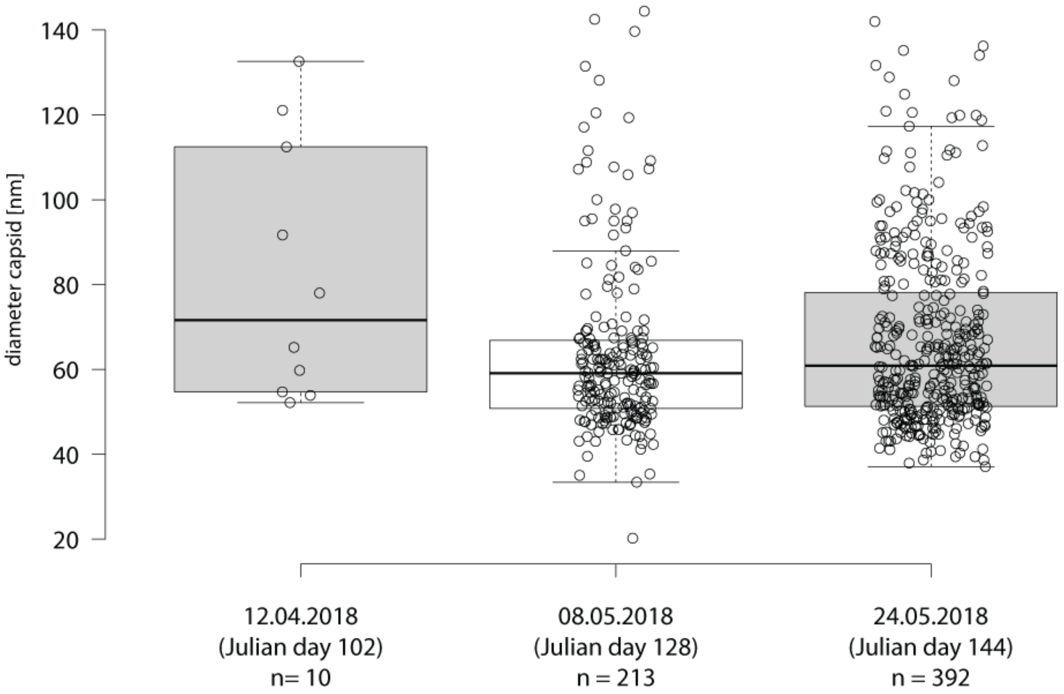


**Figure 4:** Capsid size distribution during the spring bloom 2018 from TEM images taken for virus counts.

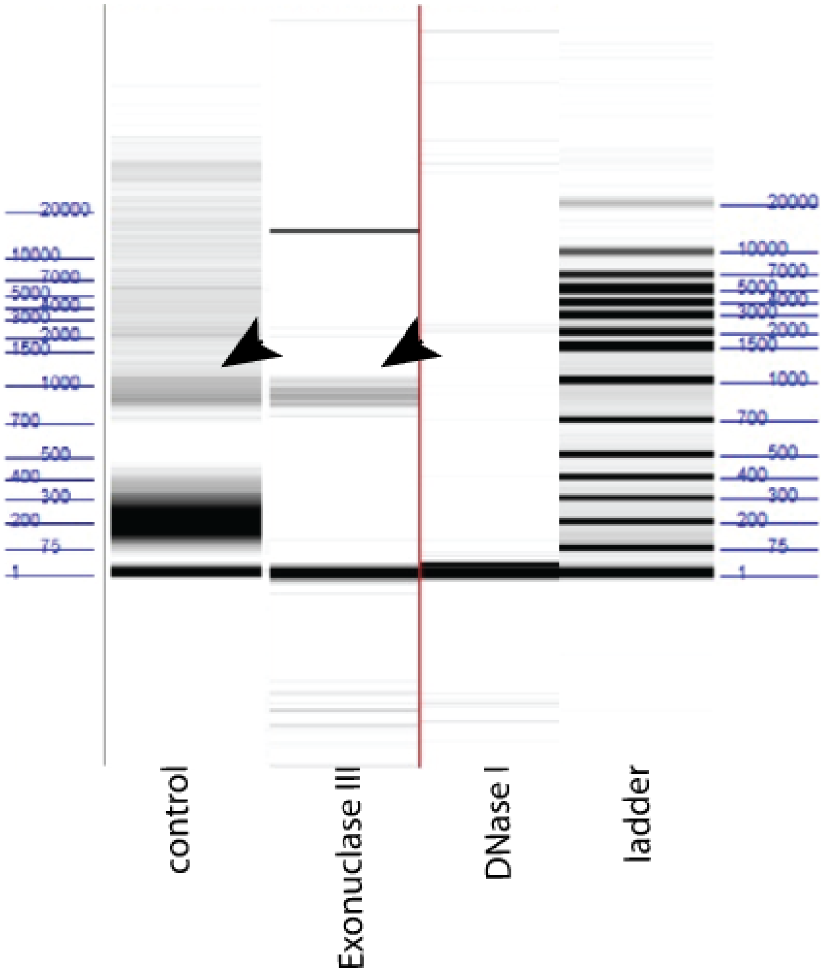


**Figure 6:** DNA digestion of *Cellulophaga* phage Omtje_1 visualized with Fragment Analyzer. Arrows indicate phage DNA band.

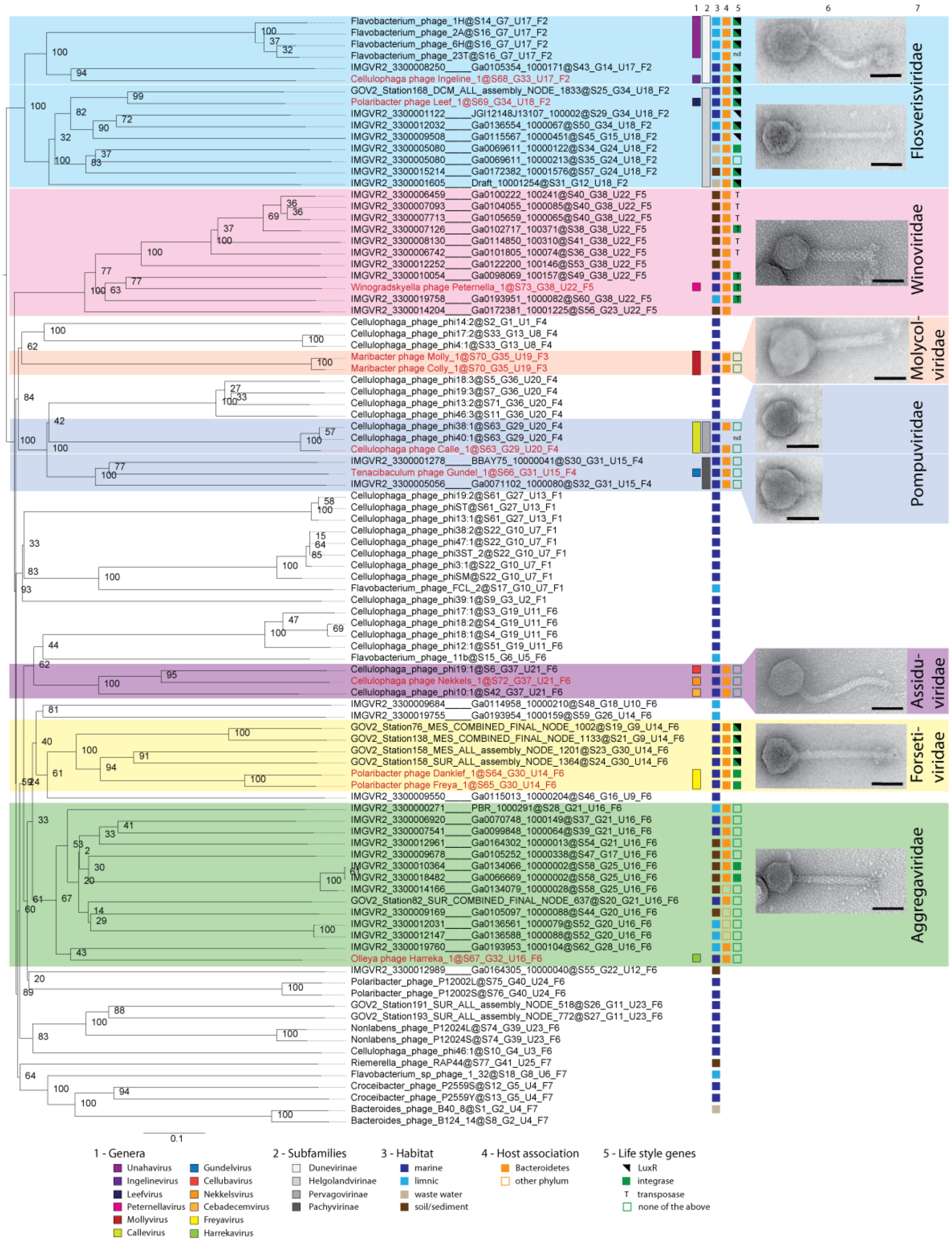


**Figure 8:** Whole genome phylogeny determined with VICTOR (amino-acid based) for the dsDNA flavophages, including environmental sequences and reference phage genomes. Our new phage isolates are depicted in red. Pseudo-bootstrap values are indicated at branches. Column 6 shows the TEM image of the negative stained new flavophage (scale bar in each TEM image 50 nm) and column 7 gives the new family name. Family and subfamily clustering is indicated at the end of the genome names.

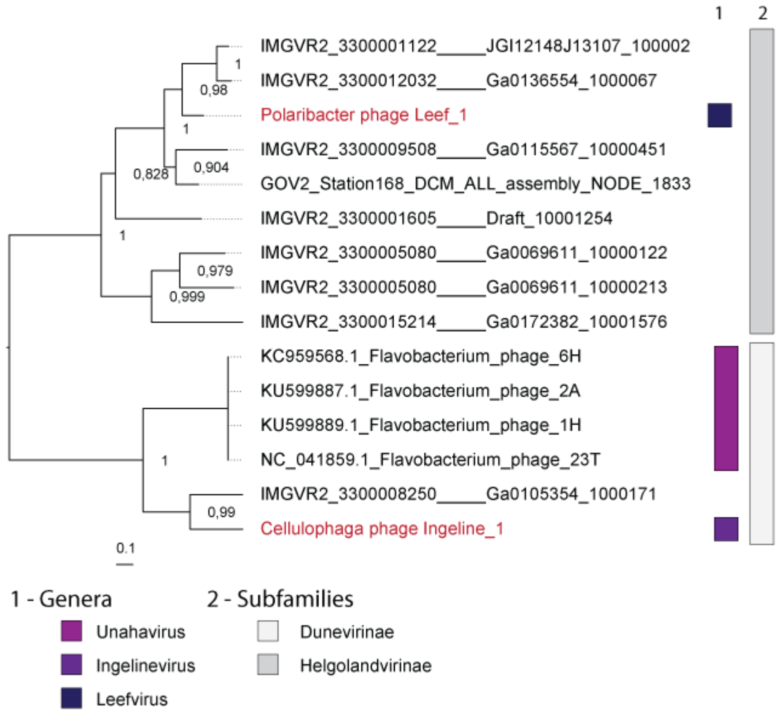


**Figure 10:** Core gene phylogeny of "Flosverisviridae". This phylogeny is based on one core gene which encodes a hypothetical protein and can be found in the annotation file with the protein cluster 11 and 5 for Ingeline and Leef, respectively.

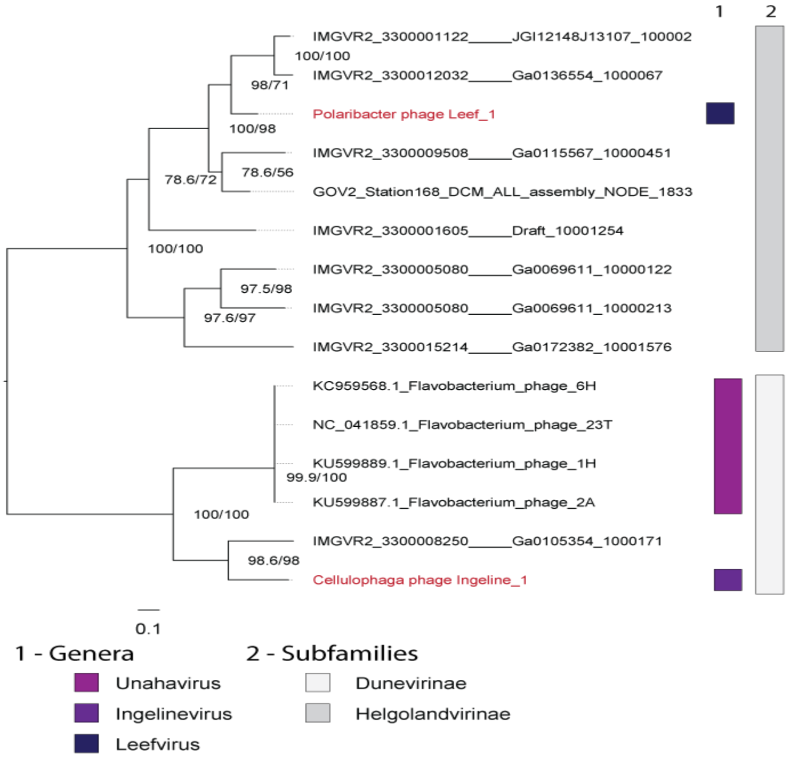

**Figure 11:** Determination of intra-family structure of the" Flosverisviridae" using MUSCLE aligned core proteins and IQ-Tree. The first branch support value is the SH-aLRT support in %, the second value is the ultrafast bootstrap support. Using the model finder the VT+F+G4 substitution model was determined as best fitting substitution model and used for the tree calculation.

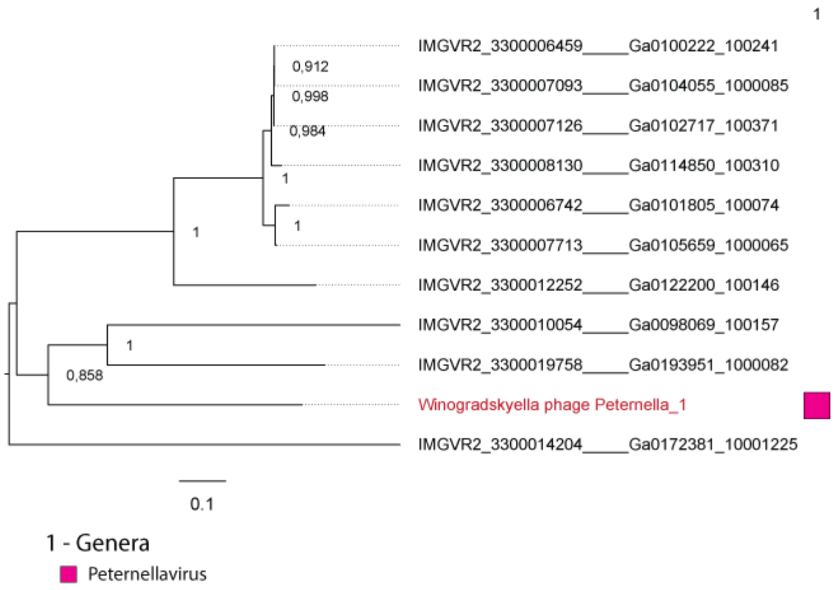


**Figure 12;** Core gene phylogeny of "Winoviridae". This phylogeny is based on 13 core genes, which can be found in the annotation file of Peternella in the following protein clusters: 2 (Head assembly cofactor/terminase), 3 (major capsid protein), 4 (Clp protease), 5 (DNA-binding transcriptional regulator), 6 (DUF2586/sheath), 8 (hp), 10 (Mu-like prophage protein gpG/neck), 11 (hp), 12 (hp/tail completion), 13 (phage protein D), 14 (hp), 15 (oxidase), 18 (nucleotidyltransferase).

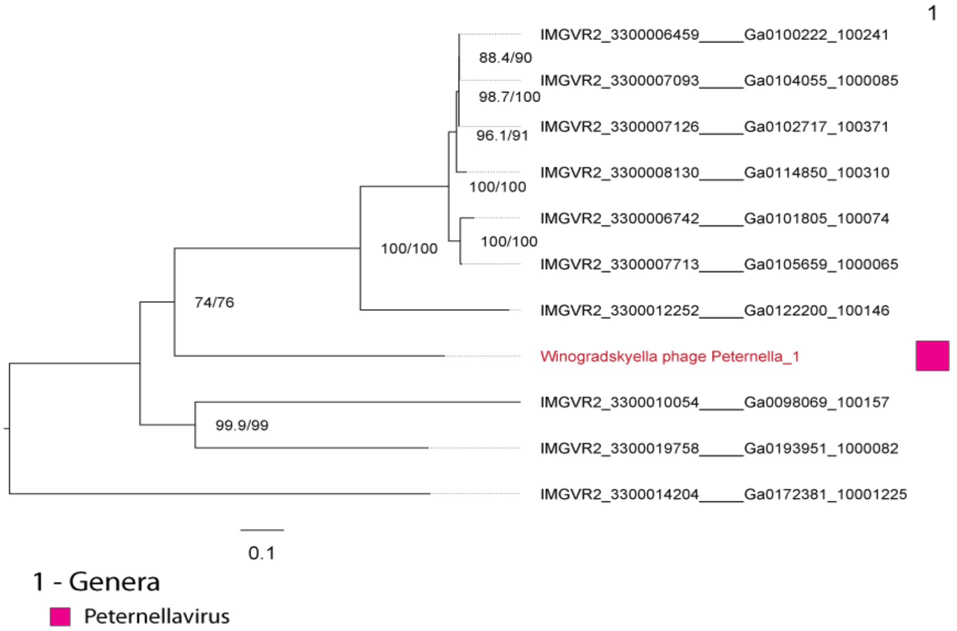


**Figure 13:** Determination of intra-family structure of the" Winoviridae" using MUSCLE aligned core proteins and IQ-Tree. The first branch support value is the SH-aLRT support in %, the second value is the ultrafast bootstrap support. Using the model finder the LG+F+I+G4 substitution model was determined as best fitting substitution model and used for the tree calculation.

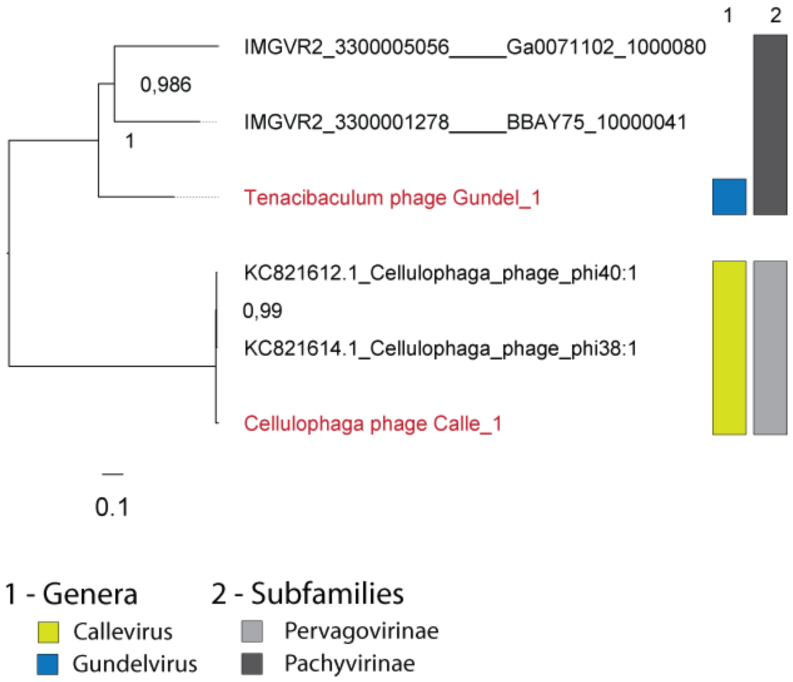


**Figure 14:** Core gene phylogeny of “Pompuviridae”. This phylogeny is based on seven core genes, which can be found in the annotation file for Calle with the protein clusters 40, 64, 65, 67, 68, 69, 71 all encoding hypothetical proteins. The protein clusters 1, 5, 6, 7, 8, 9, 10 for Gundel encoded structural proteins with protein cluster 8 encoding a portal protein and protein cluster 7 a hypothetical protein.

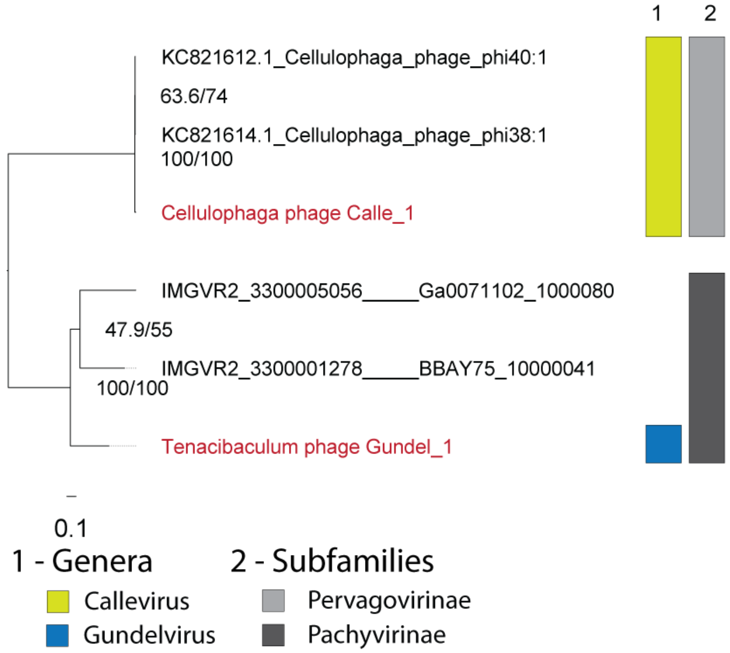


**Figure 15:** Determination of intra-family structure of the “Pompuviridae” using MUSCLE aligned core proteins and IQ-Tree. The first branch support value is the SH-aLRT support in %, the second value is the ultrafast bootstrap support. Using the model finder the LG+F+G4 substitution model was determined as best fitting substitution model and used for the tree calculation.

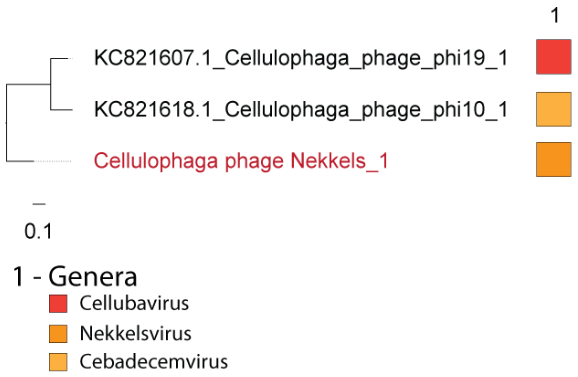


**Figure 16:** Core gene phylogeny of "Assiduviridae". This phylogeny is based on 32 core genes, which can be found in the annotation file of Nekkels with the following protein clusters and annotations: 6 (NinG recombinase protein), 13 (calcineurin-like phosphoesterase superfamily domain protein), 14 (hp), 17 (pectate lyase), 21 (hp), 22 (DNA primase/helicase, TOPRIM), 23 (structural protein), 28 (hp), 29 (hp), 30 (pectate lyase), 41 (hp), 47 (hp), 48 (hp), 49 (hp), 50 (hp), 51 (hp), 52 (hp), 53 (hp), 54 (hp), 55 (hp), 57 (hp), 58 (hp), 59 (hp), 61(hp), 62 (hp), 63 (hp), 64 (hp), 65 (hp),66 (hp), 67 (hp), 68 (hp), 85 (hp).

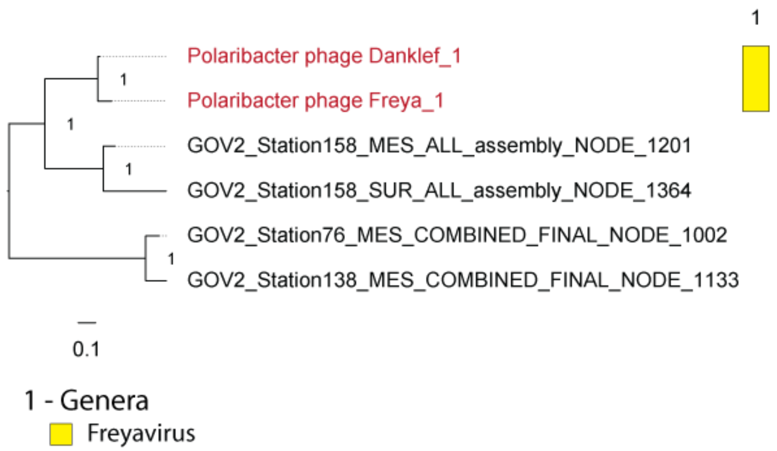


**Figure 17:** Core gene phylogeny of" Forsetiviridae". This phylogeny is based on 12 core genes, which can be found in the annotation file of Danklef and Freya with the following protein clusters and annotations: 1 (tape measure protein), 3 (structural protein), 4 (portal protein), 5 (structural protein), 6 (structural protein), 7(structural protein), 8 (hp), 9 (hp), 10 (terminase large subunit), 11 (structural protein), 13 (terminase small subunit), 18 (integrase).

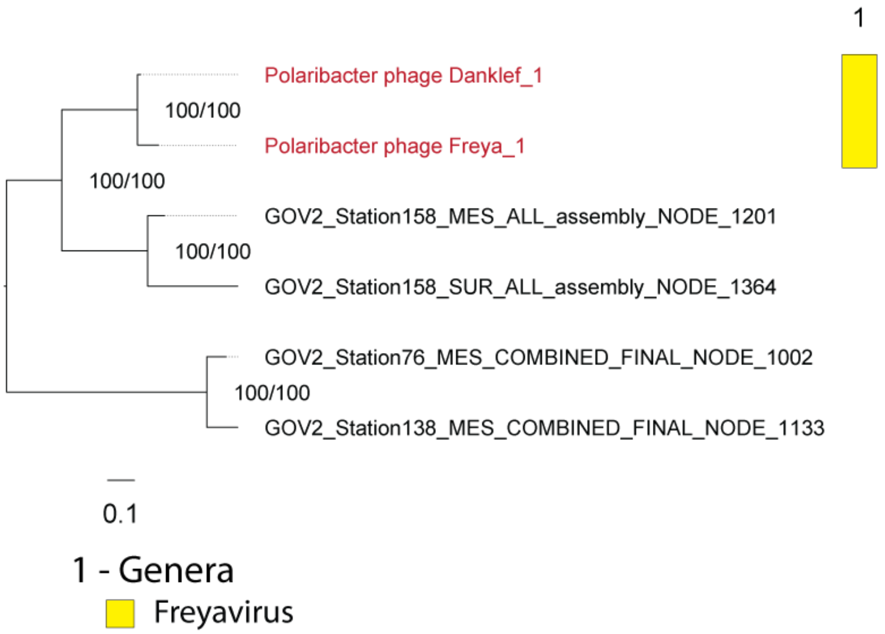


**Figure 18:** Determination of intra-family structure of the "Forsetiviridae" using MUSCLE aligned core proteins and IQ-Tree. The first branch support value is the SH-aLRT support in %, the second value is the ultrafast bootstrap support. Using the model finder the VT+F+I+G4 substitution model was determined as best fitting substitution model and used for the tree calculation.

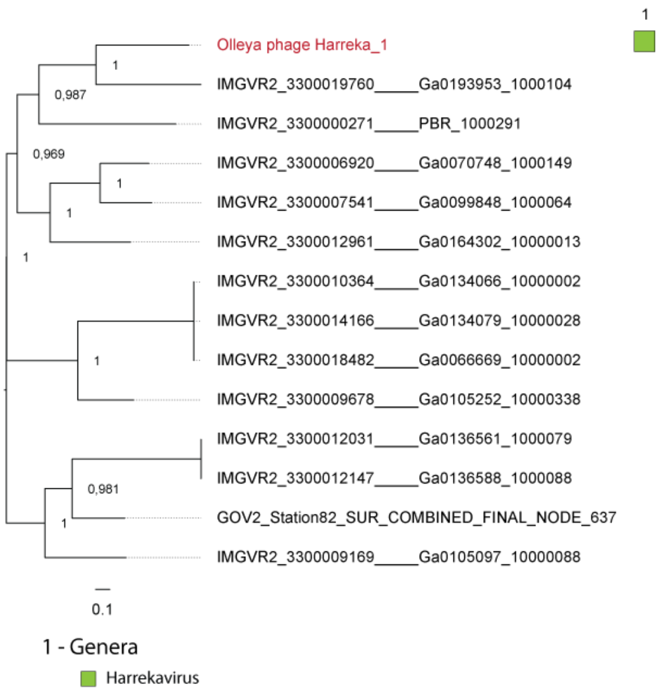


**Figure 19:** Core gene phylogeny of “Aggregaviridae”. This phylogeny is based on three core genes, which can be found in the annotation file of Harreka with the following protein clusters and annotations: 2 (tail tape measure protein), 6 (hp), 10 (hp).

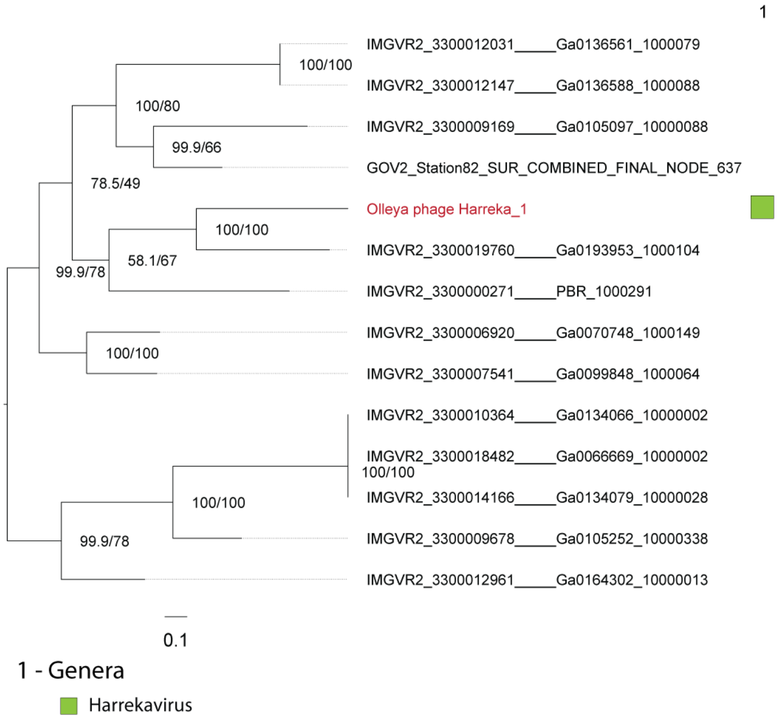


**Figure 20:** Determination of intra-family structure of the “Aggregaviridae” using MUSCLE aligned core proteins and IQ-Tree. The first branch support value is the SH-aLRT support in %, the second value is the ultrafast bootstrap support. Using the model finder the PMB+F+I+G4 substitution model was determined as best fitting substitution model and used for the tree calculation.

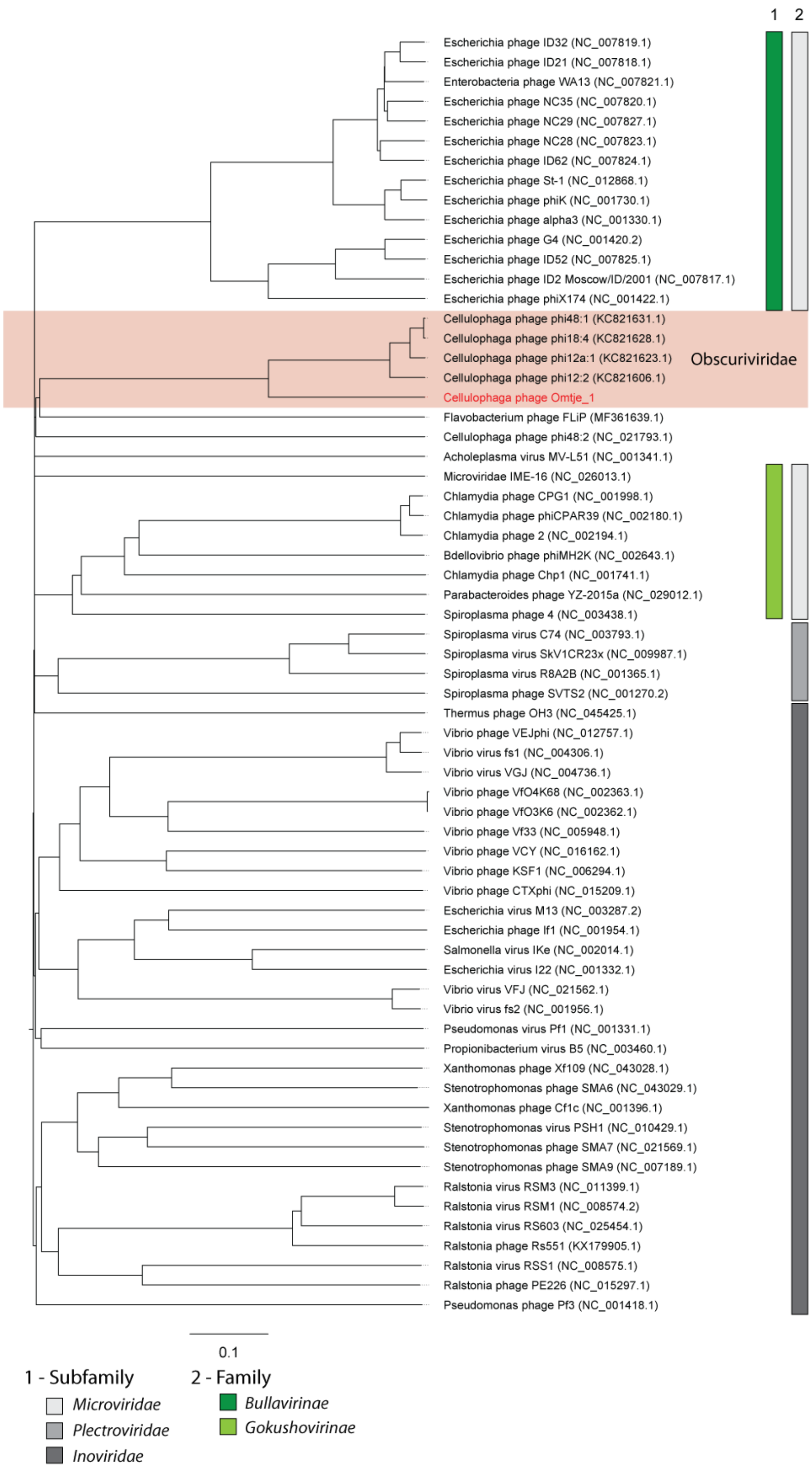


**Figure 21:** ViPTree of ssDNA viruses.

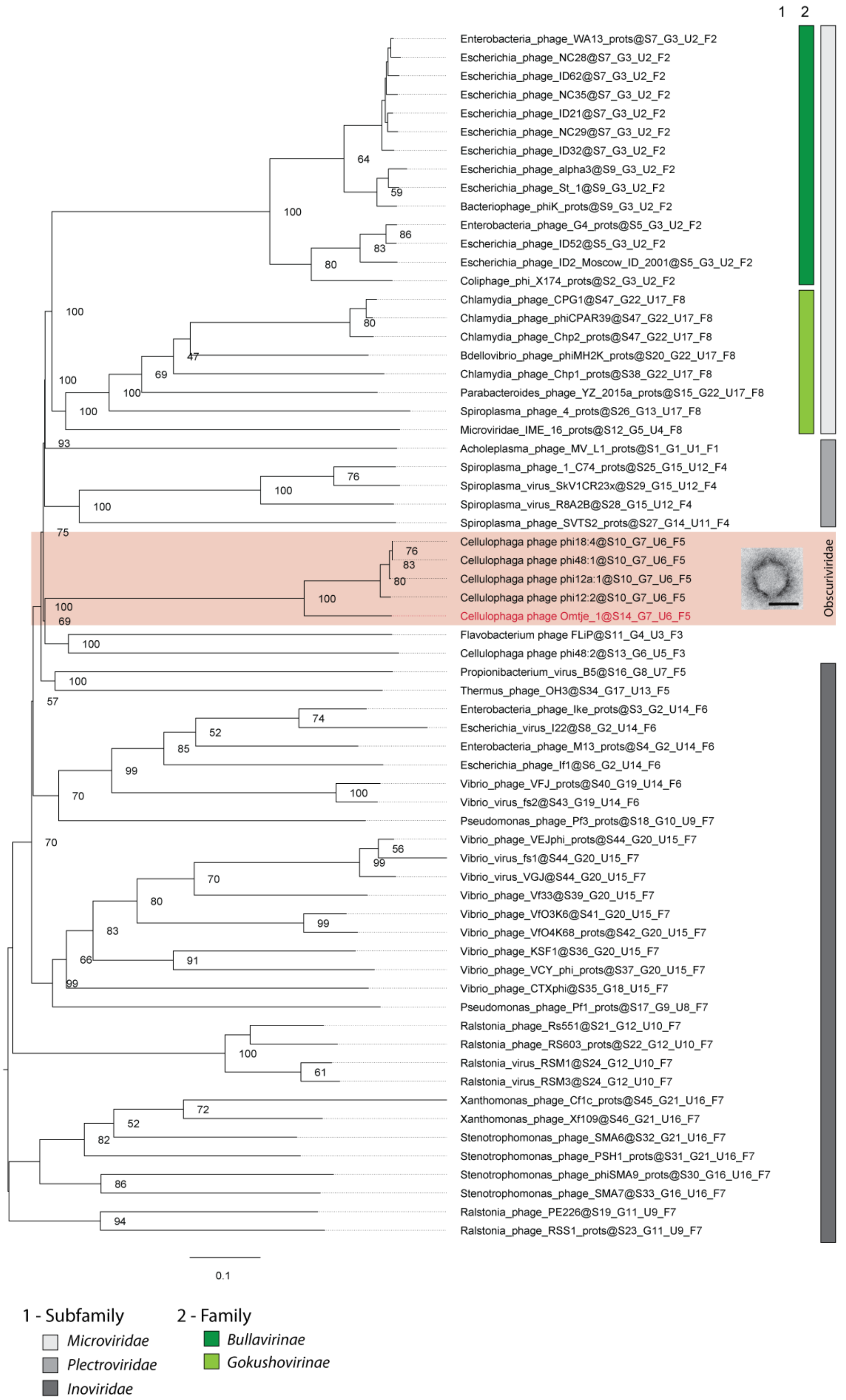


**Figure 22:** Whole genome phylogeny determined with VICTOR (amino-acid based) for the ssDNA phages. Our new phage isolate is depicted in red. Pseudo-bootstrap values are indicated at branches. TEM image of the negative stained new flavophage (scale bar in each TEM image 50 nm) and the new family name are depicted.Family and subfamily clustering is indicated at the end of the genome names.

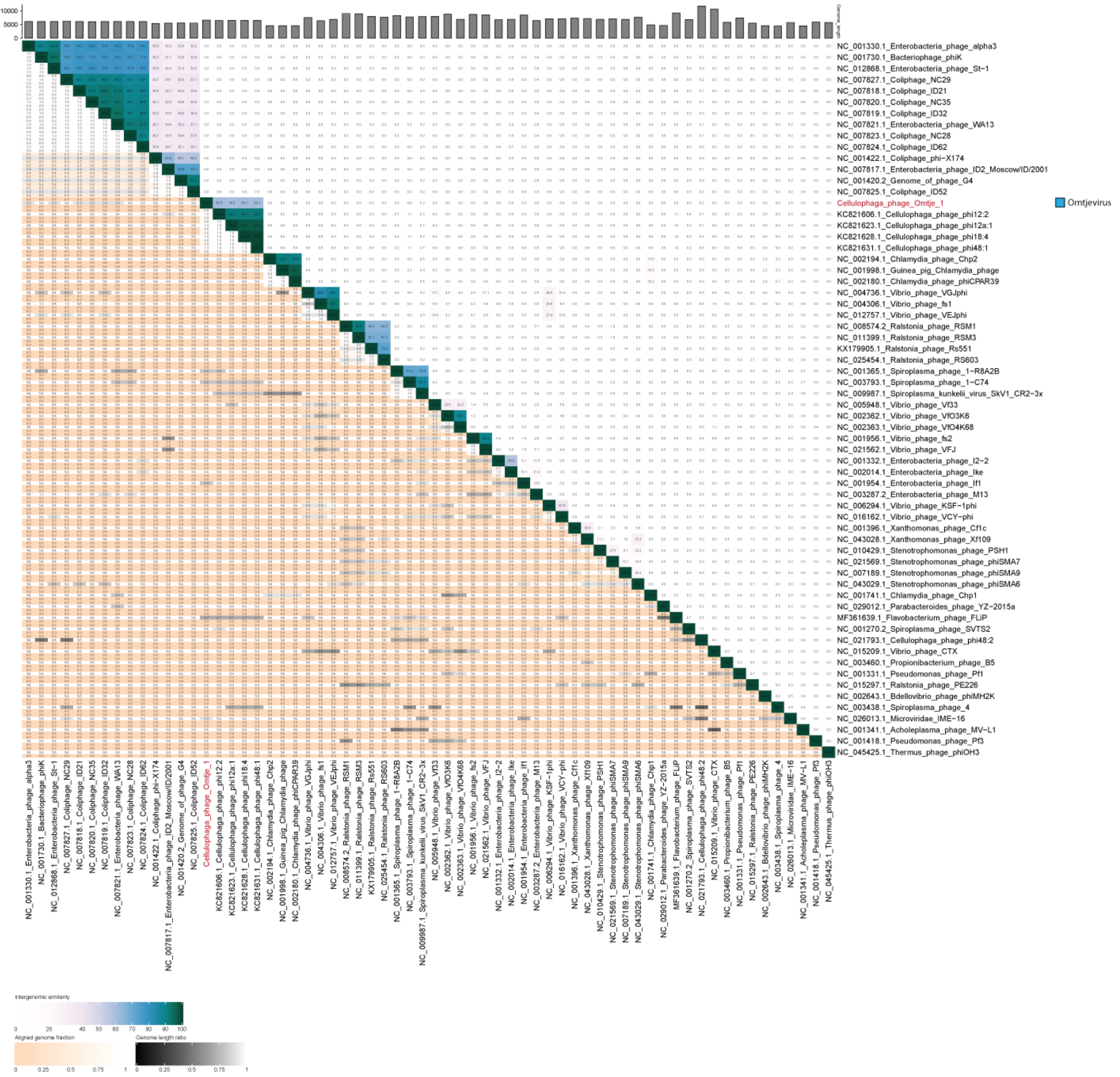


**Figure 24:** Intergenomic nucleotide similarity matrix of Omtje and known ssDNA phages.

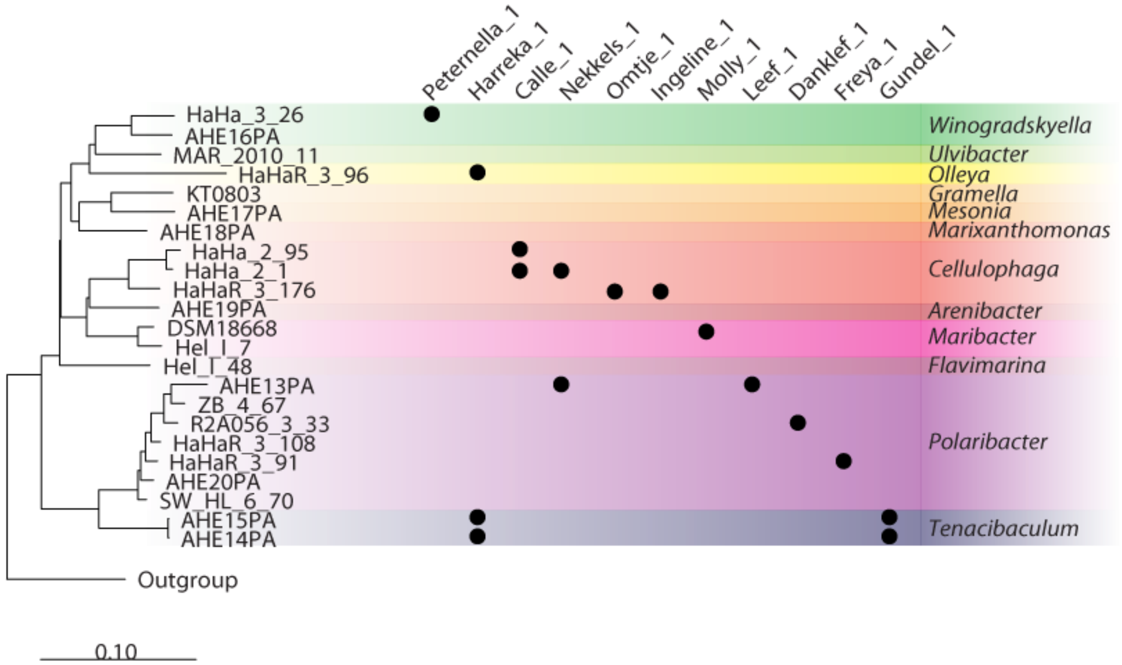


**Figure 25:** Determination of host range using all bacterial isolates, which were used as isolation source for phages in 2017 and 2018. Bacterial isolates are ordered in a 16S rRNA neighbor joining tree and on the top are the phages sorted by the host order in the tree. Colly_1 was not tested.

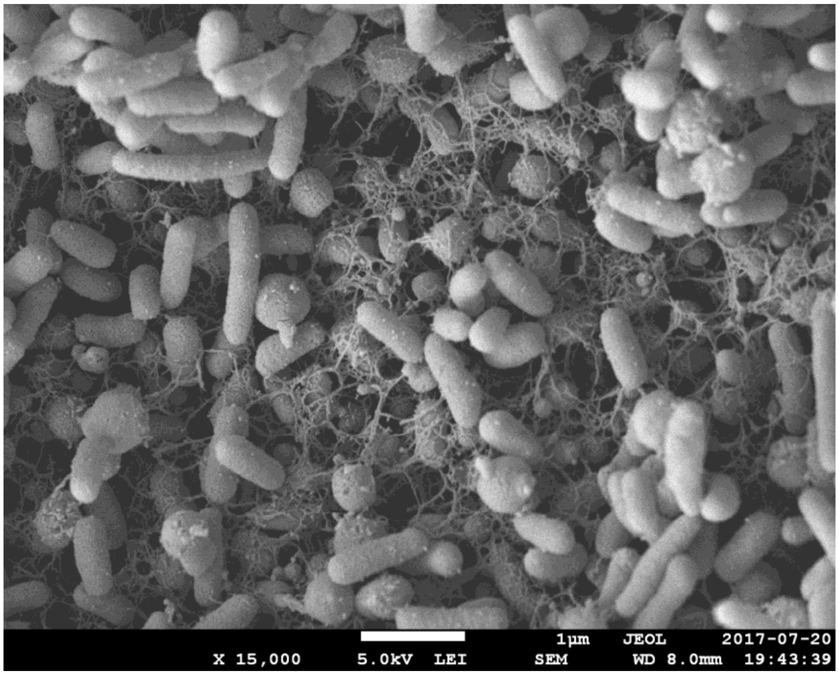


**Figure 26:** SEM image of Cellulophaga sp. HaHaR_3_176 colony grown on marine broth plates. The biofilm produced by the bacteria is visible as a fibrillar meshwork.

### Tables

**Table 1:** Bacteria used for flavophage isolation in 2017 and 2018.

| Genus | Strain | DSMZ accession number | year of isolation | Associated studies | GenBank accession number for 16S sequence | used in 2017 | used in 2018 |
|---|---|---|---|---|---|---|---|
| *Gramella* | KT0803 | DSM17595 | 1999 | (31) | AF235117.1 | + | + |
| *Cellulophaga* | HaHaR_3_176 | DSM111152 | 2016 | (32) | LT724228.1 | + | + |
| *Maribacter* | Hel_1_7 | - | 2010 | (23) | JX854136.1 | + | + |
| *Maribacter* | KT02ds 18-6 | DSM18668 | 1998 | (33) | AM712900.1 | + | + |
| *Leeuwenhoekiella* | Hel_1_48 | - | 2010 | (23) | JX854131.1 | + | - |
| *Ulvibacter* | Mar_2010_11 | - | 2010 | (23) | JX854389.1 | + | - |
| *Winogradskyella* | AHE16PA | - | 2017 | | MT667377 | - | + |
| *Mesonia* | AHE17PA | - | 2017 | | MT667380 | - | + |
| *Marixanthomonas* | AHE18PA | - | 2017 | | MT667381 | - | + |
| *Winogradskyella* | HaHa_3_26 | DSM111041 | 2016 | (32) | | - | + |
| *Cellulophaga* | HaHa_2_1 | DSM111038 | 2016 | (32) | | - | + |
| *Polaribacter* | HaHaR_3_91 | DSM111048 | 2016 | (32) | | - | + |
| *Cellulophaga* | HaHa_2_95 | DSM111037 | 2016 | (32) | | - | + |
| *Arenibacter* | AHE19PA | - | 2017 | | MT667378 | - | + |
| *Tenacibaculum* | AHE14PA | DSM111040 | 2017 | | | - | + |
| *Polaribacter* | AHE20PA | - | 2017 | | MT667376 | - | + |
| *Tenacibaculum* | AHE15PA | DSM111039 | 2017 | | | - | + |
| *Polaribacter* | HaHaR_3_108 | - | 2016 | (32) | MT667379 | - | + |
| *Olleya* | HaHaR_3_96 | DSM111044 | 2016 | (32) | | - | + |
| *Polaribacter* | R2A056_3_33 | DSM111047 | 2016 | (32) | | - | + |
| *Polaribacter* | SW_HL_6_70 | - | 2016 | (32) | MT667382 | - | + |
| *Polaribacter* | ZB_4_67 | - | 2016 | (32) | MT667383 | - | + |
| *Polaribacter* | AHE13PA | DSM111061 | 2017 | | | - | + |

**Table 2:** Detailed information of metagenomes used for read mapping

| julian day | date | size fraction | accession number |
|---|---|---|---|
| 76 | 16.03.2016 | 0.2 - 3 µm | PRJNA441607 |
| 81 | 21.03.2016 | 0.2 - 3 µm | PRJNA441608 |
| 91 | 31.03.2016 | 0.2 - 3 µm | PRJNA441609 |
| 103 | 12.04.2016 | 0.2 - 3 µm | PRJNA441610 |
| 110 | 19.04.2016 | 0.2 - 3 µm | PRJNA441611 |
| 117 | 26.04.2016 | 0.2 - 3 µm | PRJNA441612 |
| 123 | 02.05.2016 | 0.2 - 3 µm | PRJNA441613 |
| 133 | 12.05.2016 | 0.2 - 3 µm | PRJNA441614 |
| 138 | 17.05.2016 | 0.2 - 3 µm | PRJNA441615 |
| 78 | 19.03.2018 | 0.2 - 3 µm | PRJEB38290 |
| | | 3 - 10 µm | PRJEB38290 |
| | | > 10 µm | PRJEB38290 |
| 93 | 03.04.2018 | 0.2 - 3 µm | PRJEB38290 |
| 95 | 05.04.2018 | 0.2 - 3 µm | PRJEB38290 |
| 100 | 10.04.2018 | 0.2 - 3 µm | PRJEB38290 |
| 102 | 12.04.2018 | 0.2 - 3 µm | PRJEB38290 |
| | | 3 - 10 µm | PRJEB38290 |
| | | > 10 µm | PRJEB38290 |
| 107 | 17.04.2018 | 0.2 - 3 µm | PRJEB38290 |
| | | 3 - 10 µm | PRJEB38290 |
| | | > 10 µm | PRJEB38290 |
| 109 | 19.04.2018 | 0.2 - 3 µm | PRJEB38290 |
| 114 | 24.04.2018 | 0.2 - 3 µm | PRJEB38290 |
| 116 | 26.04.2018 | 0.2 - 3 µm | PRJEB38290 |
| | | 3 - 10 µm | PRJEB38290 |
| | | > 10 µm | PRJEB38290 |
| 122 | 02.05.2018 | 0.2 - 3 µm | PRJEB38290 |
| 123 | 03.05.2018 | 0.2 - 3 µm | PRJEB38290 |
| 128 | 08.05.2018 | 0.2 - 3 µm | PRJEB38290 |
| | | 3 - 10 µm | PRJEB38290 |
| | | > 10 µm | PRJEB38290 |
| 131 | 11.05.2018 | 0.2 - 3 µm | PRJEB38290 |
| | | 3 - 10 µm | PRJEB38290 |
| | | > 10 µm | PRJEB38290 |
| 135 | 15.05.2018 | 0.2 - 3 µm | PRJEB38290 |
| 137 | 17.05.2018 | 0.2 - 3 µm | PRJEB38290 |
| 142 | 22.05.2018 | 0.2 - 3 µm | PRJEB38290 |
| | | 3 - 10 µm | PRJEB38290 |
| | | > 10 µm | PRJEB38290 |
| 144 | 24.05.2018 | 0.2 - 3 µm | PRJEB38290 |
| 149 | 29.05.2018 | 0.2 - 3 µm | PRJEB38290 |
| | | 3 - 10 µm | |
| | | > 10 µm | |

**Table 3:** Virus to bacteria ratios for Julian day 102, 128, and 144 using transmission electron microscopy (TEM) and epifluorescence light microscopy (LM).

| Julian day | TEM | LM |
|---|---|---|
| 102 | 1.8 | 36.3 |
| 128 | 2.6 | 79.3 |
| 144 | 3.1 | 52.3 |

**Table 4:** List of obtained flavophage isolates.

| Phage isolation name | Culture accession | sampling date | host | host genus | phage subgroup | phage group | genome size |
|---|---|---|---|---|---|---|---|
| E30_2/2/2/1/F | DSM111231 | 03.04.2018 | HaHa_2_95 | *Cellulophaga* | Calle_1 | Calle | 72979 |
| E30_2/2/2/2/F | | 03.04.2018 | HaHa_2_95 | *Cellulophaga* | Calle_2 | Calle | 72979 |
| Re30_10/1s/1 final | | 03.05.2018 | HaHa_2_95 | *Cellulophaga* | Calle_3 | Calle | 72980 |
| E30_2/1/1/F | | 03.04.2018 | HaHa_2_95 | *Cellulophaga* | Calle_2 | Calle | 72979 |
| E47_12/2/1 final | DSM111232 | 15.05.2018 | R2A056_3_33 | *Polaribacter* | Danklef_1 | Danklef | 47186 |
| E47_12/1/1 final | | 15.05.2018 | R2A056_3_33 | *Polaribacter* | Danklef_2 | Danklef | 47302 |
| E47_12/3/1 final | | 15.05.2018 | R2A056_3_33 | *Polaribacter* | Danklef_2 | Danklef | 47302 |
| E47_14/2/1 final | | 24.05.2018 | R2A056_3_33 | *Polaribacter* | Danklef_3 | Danklef | 47396 |
| E47_14/3/1 final | | 24.05.2018 | R2A056_3_33 | *Polaribacter* | Danklef_4 | Danklef | 47426 |
| E47_14/1/1 final | | 24.05.2018 | R2A056_3_33 | *Polaribacter* | Danklef_5 | Danklef | 48177 |
| E27_2/2/1 final | DSM111233 | 03.04.2018 | HaHaR_3_91 | *Polaribacter* | Freya_1 | Freya | 43978 |
| E27_2/2/2 final | | 03.04.2018 | HaHaR_3_91 | *Polaribacter* | Freya_1 | Freya | 43978 |
| E27_4/1/2 final | DSM111234 | 10.04.2018 | HaHaR_3_91 | *Polaribacter* | Freya_10 | Freya | 48920 |
| E27_8/1s/1 final | | 24.04.2018 | HaHaR_3_91 | *Polaribacter* | Freya_2 | Freya | 44820 |
| E27_8/2s/1 final | | 24.04.2018 | HaHaR_3_91 | *Polaribacter* | Freya_3 | Freya | 44820 |
| E27_8/2s/2 final | | 24.04.2018 | HaHaR_3_91 | *Polaribacter* | Freya_3 | Freya | 44820 |
| E27_8/2s/3 final | | 24.04.2018 | HaHaR_3_91 | *Polaribacter* | Freya_3 | Freya | 44820 |
| E27_8/2s/4 final | | 24.04.2018 | HaHaR_3_91 | *Polaribacter* | Freya_3 | Freya | 44820 |
| E27_8/2s/5 final | | 24.04.2018 | HaHaR_3_91 | *Polaribacter* | Freya_3 | Freya | 44820 |
| E27_8/2s/6 final | | 24.04.2018 | HaHaR_3_91 | *Polaribacter* | Freya_3 | Freya | 44820 |
| E27_8/3s/2 final | | 24.04.2018 | HaHaR_3_91 | *Polaribacter* | Freya_3 | Freya | 44820 |
| E27_8/3s/1 final | | 24.04.2018 | HaHaR_3_91 | *Polaribacter* | Freya_4 | Freya | 44820 |
| E27_2/1/1 final | | 03.04.2018 | HaHaR_3_91 | *Polaribacter* | Freya_5 | Freya | 45722 |
| E27_2/1/2 final | | 03.04.2018 | HaHaR_3_91 | *Polaribacter* | Freya_5 | Freya | 45722 |
| E27_2/2/1/F | | 03.04.2018 | HaHaR_3_91 | *Polaribacter* | Freya_5 | Freya | 45722 |
| E27_2/3/1 final | | 03.04.2018 | HaHaR_3_91 | *Polaribacter* | Freya_5 | Freya | 45722 |
| E27_2/3/2 final | | 03.04.2018 | HaHaR_3_91 | *Polaribacter* | Freya_5 | Freya | 45722 |
| E27_2/3/3 final | | 03.04.2018 | HaHaR_3_91 | *Polaribacter* | Freya_5 | Freya | 45722 |
| E27_4/2/1 final | | 10.04.2018 | HaHaR_3_91 | *Polaribacter* | Freya_6 | Freya | 46194 |
| E27_10/1s/1 final | | 03.05.2018 | HaHaR_3_91 | *Polaribacter* | Freya_7 | Freya | 46194 |
| E27_4/1/3 final | | 10.04.2018 | HaHaR_3_91 | *Polaribacter* | Freya_8 | Freya | 48018 |
| E27_2/1/1/F | | 03.04.2018 | HaHaR_3_91 | *Polaribacter* | Freya_9 | Freya | 48613 |
| E27_4/1/1 final | | 10.04.2018 | HaHaR_3_91 | *Polaribacter* | Freya_9 | Freya | 48613 |
| E27_4/3/1 final | | 10.04.2018 | HaHaR_3_91 | *Polaribacter* | Freya_9 | Freya | 48613 |
| E37_7/1s/1 final | DSM111235 | 19.04.2018 | AHE14PA | *Tenacibaculum* | Gundel_1 | Gundel | 78511 |
| E37_7/1s/2 final | | 19.04.2018 | AHE14PA | *Tenacibaculum* | Gundel_1 | Gundel | 78511 |
| E37_7/1s/3 final | | 19.04.2018 | AHE14PA | *Tenacibaculum* | Gundel_1 | Gundel | 78511 |
| E42_11/1/1 final | | 08.05.2018 | AHE15PA | *Tenacibaculum* | Gundel_1 | Gundel | 78511 |
| E42_11/1s/1 final | | 08.05.2018 | AHE15PA | *Tenacibaculum* | Gundel_1 | Gundel | 78511 |
| E42_11/2/1 final | | 08.05.2018 | AHE15PA | *Tenacibaculum* | Gundel_1 | Gundel | 78511 |
| E42_11/2s/1 final | | 08.05.2018 | AHE15PA | *Tenacibaculum* | Gundel_1 | Gundel | 78511 |
| E42_11/3/1 final | | 08.05.2018 | AHE15PA | *Tenacibaculum* | Gundel_1 | Gundel | 78511 |
| E42_11/4/1 final | | 08.05.2018 | AHE15PA | *Tenacibaculum* | Gundel_1 | Gundel | 78511 |
| E42_11/5/1 final | | 08.05.2018 | AHE15PA | *Tenacibaculum* | Gundel_1 | Gundel | 78511 |
| E46_12/1/1 final | DSM111256 | 15.05.2018 | HaHaR_3_96 | *Olleya* | Harreka_1 | Harreka | 43175 |
| E46_12/2/1 final | | 15.05.2018 | HaHaR_3_96 | *Olleya* | Harreka_1 | Harreka | 43175 |
| E46_12/3/1 final | | 15.05.2018 | HaHaR_3_96 | *Olleya* | Harreka_1 | Harreka | 43175 |
| E46_14/1/1 final | | 24.05.2018 | HaHaR_3_96 | *Olleya* | Harreka_1 | Harreka | 43175 |
| E46_14/2/1 final | | 24.05.2018 | HaHaR_3_96 | *Olleya* | Harreka_1 | Harreka | 43175 |
| E46_14/3/1 final | | 24.05.2018 | HaHaR_3_96 | *Olleya* | Harreka_1 | Harreka | 43175 |
| E46_14/4/1 final | | 24.05.2018 | HaHaR_3_96 | *Olleya* | Harreka_1 | Harreka | 43175 |
| 4_2/1_II/2 final | DSM111236 | 21.03.2017 | HaHaR_3_176 | *Cellulophaga* | Ingeline_1 | Ingeline | 42624 |
| 4_2/1_II/4 final | | 21.03.2017 | HaHaR_3_176 | *Cellulophaga* | Ingeline_2 | Ingeline | 42624 |

| | | | | | | | |
|---|---|---|---|---|---|---|---|
| 4_2/1_II/1 final | | 21.03.2017 | HaHaR_3_176 | *Cellulophaga* | Ingeline_3 | Ingeline | 42624 |
| 4_2/2/2 final | | 21.03.2017 | HaHaR_3_176 | *Cellulophaga* | Ingeline_3 | Ingeline | 42624 |
| 4_2/2_II/1 final | | 21.03.2017 | HaHaR_3_176 | *Cellulophaga* | Ingeline_4 | Ingeline | 42624 |
| 4_2/3/3 final | | 21.03.2017 | HaHaR_3_176 | *Cellulophaga* | Ingeline_5 | Ingeline | 42625 |
| E4_10/2/1 final | | 03.05.2018 | HaHaR_3_176 | *Cellulophaga* | Ingeline_6 | Ingeline | 42625 |
| 4_2/5/1 final | | 21.03.2017 | HaHaR_3_176 | *Cellulophaga* | Ingeline_7 | Ingeline | 42625 |
| E4_10GS/2/1 final | | 03.05.2018 | HaHaR_3_176 | *Cellulophaga* | Ingeline_7 | Ingeline | 42625 |
| 4_2/2/3 final | | 21.03.2017 | HaHaR_3_176 | *Cellulophaga* | Ingeline_8 | Ingeline | 42775 |
| E4_10/3/1 final | | 03.05.2018 | HaHaR_3_176 | *Cellulophaga* | Ingeline_8 | Ingeline | 42776 |
| E4_10GS/3/1 final | | 03.05.2018 | HaHaR_3_176 | *Cellulophaga* | Ingeline_9 | Ingeline | 42797 |
| E50_14/1/1 final | DSM111238 | 24.05.2018 | AHE13PA | *Polaribacter* | Leef_1 | Leef | 37547 |
| E50_14/2/1 final | | 24.05.2018 | AHE13PA | *Polaribacter* | Leef_1 | Leef | 37547 |
| E50_14/3/1 final | | 24.05.2018 | AHE13PA | *Polaribacter* | Leef_1 | Leef | 37547 |
| 8_5/3/1 final | DSM111257 | 11.04.2017 | DSM18668 | *Maribacter* | Molly_1 | Molly | 124695 |
| 8_5/3/2 final | | 11.04.2017 | DSM18668 | *Maribacter* | Molly_1 | Molly | 124695 |
| 8_5/6/1 final | | 11.04.2017 | DSM18668 | *Maribacter* | Molly_1 | Molly | 124695 |
| 8_5/7/1 final | | 11.04.2017 | DSM18668 | *Maribacter* | Molly_1 | Molly | 124695 |
| 8_5/7/2 final | | 11.04.2017 | DSM18668 | *Maribacter* | Molly_1 | Molly | 124695 |
| 8_5/1/2 final | | 11.04.2017 | DSM18668 | *Maribacter* | Molly_2 | Molly | 124695 |
| 8_5/6/2 final | | 11.04.2017 | DSM18668 | *Maribacter* | Molly_3 | Molly | 124695 |
| 8_4/4/1 final | | 04.04.2017 | DSM18668 | *Maribacter* | Molly_4 | Molly | 125038 |
| 8_4/5/1 final | | 04.04.2017 | DSM18668 | *Maribacter* | Molly_5 | Molly | 124898 |
| 8_4/5/2 final | | 04.04.2017 | DSM18668 | *Maribacter* | Molly_5 | Molly | 124898 |
| 8_4/3/2 final | | 04.04.2017 | DSM18668 | *Maribacter* | Molly_7 | Molly | 125344 |
| 8_5/8/1 final | DSM111252 | 11.04.2017 | DSM18668 | *Maribacter* | Colly_1 | Molly | 124169 |
| 8_5/8/2 final | | 11.04.2017 | DSM18668 | *Maribacter* | Colly_1 | Molly | 124169 |
| SW26_10/1s/1 final | DSM111239 | 03.05.2018 | HaHa_2_1 | *Cellulophaga* | Nekkels_1 | Nekkels | 53385 |
| E26_2/1/1/F | | 03.04.2018 | HaHa_2_1 | *Cellulophaga* | Nekkels_2 | Nekkels | 54332 |
| E26_2/1/2/F | | 03.04.2018 | HaHa_2_1 | *Cellulophaga* | Nekkels_2 | Nekkels | 54332 |
| 4_1/1/1 final | DSM111240 | 14.03.2017 | HaHaR_3_176 | *Cellulophaga* | Omtje_1 | Omtje | 6558 |
| 4_1/1/2 final | | 14.03.2017 | HaHaR_3_176 | *Cellulophaga* | Omtje_1 | Omtje | 6558 |
| 4_1/2/1 final | | 14.03.2017 | HaHaR_3_176 | *Cellulophaga* | Omtje_1 | Omtje | 6558 |
| 4_1/2/2 final | | 14.03.2017 | HaHaR_3_176 | *Cellulophaga* | Omtje_1 | Omtje | 6558 |
| 4_1/3/1 final | | 14.03.2017 | HaHaR_3_176 | *Cellulophaga* | Omtje_1 | Omtje | 6558 |
| 4_1/3/2 final | | 14.03.2017 | HaHaR_3_176 | *Cellulophaga* | Omtje_1 | Omtje | 6558 |
| 4_1/4/1 final | | 14.03.2017 | HaHaR_3_176 | *Cellulophaga* | Omtje_1 | Omtje | 6558 |
| 4_1/4/2 final | | 14.03.2017 | HaHaR_3_176 | *Cellulophaga* | Omtje_1 | Omtje | 6558 |
| 4_2/2/1 final | | 21.03.2017 | HaHaR_3_176 | *Cellulophaga* | Omtje_2 | Omtje | 6558 |
| 4_2/1_II/3 final | | 21.03.2017 | HaHaR_3_176 | *Cellulophaga* | Omtje_3 | Omtje | 6558 |
| 4_2/3/1 final | | 21.03.2017 | HaHaR_3_176 | *Cellulophaga* | Omtje_3 | Omtje | 6558 |
| 4_2/3_II/2 final | | 21.03.2017 | HaHaR_3_176 | *Cellulophaga* | Omtje_3 | Omtje | 6558 |
| 4_2/4/1 final | | 21.03.2017 | HaHaR_3_176 | *Cellulophaga* | Omtje_3 | Omtje | 6558 |
| 4_2/4/2 final | | 21.03.2017 | HaHaR_3_176 | *Cellulophaga* | Omtje_3 | Omtje | 6558 |
| 4_2/6/1 final | | 21.03.2017 | HaHaR_3_176 | *Cellulophaga* | Omtje_3 | Omtje | 6558 |
| E4_10GS/1/1 final | | 03.05.2018 | HaHaR_3_176 | *Cellulophaga* | Omtje_3 | Omtje | 6558 |
| E4_11/1/1 final | | 08.05.2018 | HaHaR_3_176 | *Cellulophaga* | Omtje_3 | Omtje | 6558 |
| E4_11/2/1 final | | 08.05.2018 | HaHaR_3_176 | *Cellulophaga* | Omtje_3 | Omtje | 6558 |
| E4_11/3/1 final | | 08.05.2018 | HaHaR_3_176 | *Cellulophaga* | Omtje_3 | Omtje | 6558 |
| 4_2/6/2 final | | 21.03.2017 | HaHaR_3_176 | *Cellulophaga* | Omtje_4 | Omtje | 6558 |
| E4_10/1/1 final | | 03.05.2018 | HaHaR_3_176 | *Cellulophaga* | Omtje_5 | Omtje | 6559 |
| 4_2/3_II/1 final | | 21.03.2017 | HaHaR_3_176 | *Cellulophaga* | Omtje_6 | Omtje | 6558 |
| E25_12/1s/1 final | DSM111241 | 15.05.2018 | HaHa_3_26 | *Winogradskyella* | Peternella_1 | Peternella | 39649 |
| E25_12/1/1 final | | 15.05.2018 | HaHa_3_26 | *Winogradskyella* | Peternella_1 | Peternella | 39649 |
| E25_12/2/1 final | | 15.05.2018 | HaHa_3_26 | *Winogradskyella* | Peternella_1 | Peternella | 39649 |
| E25_12/3/1 final | | 15.05.2018 | HaHa_3_26 | *Winogradskyella* | Peternella_1 | Peternella | 39649 |

**Table 5:** Sampling site of metagenome derived contigs. * indicated multiple hits.

| Family | ID | source | viral spacer match | metagenomic spacer match | %identity | start | end | length | evalue | bacteria_taxonomic_affiliation | bacterium hit | annotation_of_hit | |
|---|---|---|---|---|---|---|---|---|---|---|---|---|---|
| **Aggregaviridae** | IMGVR2_3300010364_____Ga0134066_10000002 | soil | 0 | 0 | 66 | 10,351 | 11582 | 1,231 | 3.70E-62 | *Flavobacteriaceae* | Gillisia sp. Hel1_33_143 | hypothetical protein | * |
| **Aggregaviridae** | IMGVR2_3300018482_____Ga0066669_10000002 | soil | 0 | 0 | 66 | 32,502 | 33733 | 1,231 | 3.72E-62 | *Flavobacteriaceae* | Gillisia sp. Hel1_33_143 | hypothetical protein | * |
| **Aggregaviridae** | IMGVR2_3300014166_____Ga0134079_10000028 | soil | 0 | 0 | 70 | 26,517 | 27010 | 493 | 6.59E-52 | *Firmicutes* | Clostridium botulinum strain B609 | DNA modification methylase | |
| **Aggregaviridae** | IMGVR2_3300009678_____Ga0105252_10000338 | groundwater sediment | 0 | 0 | 72 | 14,796 | 15415 | 619 | 1.62E-78 | *Bacteroidetes* | Mucilaginibacter sp. BJC16-A31 | structural protein | |
| **Aggregaviridae** | IMGVR2_3300006920_____Ga0070748_1000149 | marine (Delaware Bay) | 0 | 4 | 76 | 5,936 | 6126 | 190 | 4.24E-29 | *Flavobacteriaceae* | Chryseobacterium sp. IHB B 17019, | hypothetical protein | |
| **Aggregaviridae** | IMGVR2_3300007541_____Ga0099848_1000064 | marine (Chesapeake Bay) | 0 | 2 | 75 | 26,396 | 26865 | 469 | 2.66E-76 | *Flavobacteriaceae* | Elizabethkingia anophelis strain 0422 | hypothetical protein | |
| **Aggregaviridae** | IMGVR2_3300012961_____Ga0164302_10000013 | soil amended with pyrogenic organic matter | 0 | 6 | 72 | 35664 | 36302 | 638 | 8.31E-83 | *Flavobacteriaceae* | Chryseobacterium gleum strain NCTC11432 | Type III effector pipB2 | |
| **Aggregaviridae** | IMGVR2_3300012031_____Ga0136561_1000079 | saline lake | 0 | 0 | 74 | 7,315 | 8013 | 698 | 3.04E-119 | *e-proteobacteria* | Campylobacter jejuni strain CJ017CCUA | DNA methylase N-4 | * |
| **Aggregaviridae** | IMGVR2_3300012147_____Ga0136588_1000088 | saline lake | 0 | 0 | 74 | 7,342 | 8040 | 698 | 3.03E-119 | *e-proteobacteria* | Campylobacter jejuni strain CJ017CCUA | DNA methylase N-4 | * |
| **Aggregaviridae** | IMGVR2_3300009169_____Ga0105097_10000088 | lake sediment | 0 | 0 | 74 | 19 | 1346 | 1,327 | 0 | *Enterobacteria* | Escherichia fergusonii strain EFCF056 plasmid pEF06 | hypothetical protein | * |
| **Aggregaviridae** | IMGVR2_3300000271_____PBR_1000291 | photobioreactor incubated | 0 | 0 | 68 | 34,771 | 35407 | 636 | 1.38E-48 | *Flavobacteriaceae* | Chryseobacterium sp. StRB126 | modification methylase | |
| **Aggregaviridae** | IMGVR2_3300019760_____Ga0193953_1000104 | river freshwater | 0 | 0 | 71 | 27,884 | 28504 | 620 | 8.57E-71 | *Bacteroidetes* | Sphingobacterium psychroaquaticum strain SJ-25 | hypothetical protein | |

| | | | | | | | | | | | | | |
|---|---|---|---|---|---|---|---|---|---|---|---|---|---|
| **Aggregaviridae** | GOV2_Station82_SUR_NODE_637 | marine (South Atlantic Ocean) | 1 | 7 | 67 | 33,644 | 34353 | 709 | 2.15E-52 | *Flavobacteriaceae* | Elizabethkingia anophelis strain 0422 | hypothetical protein | |
| **Forsetiviridae** | GOV2_Station158_MES_NODE_1201 | marine (Norwegian Sea) | 0 | 0 | 68 | 38,647 | 39569 | 922 | 2.76E-77 | *Flavobacteriaceae* | Tenacibaculum sp. DSM 106434 | integrase | * |
| **Forsetiviridae** | GOV2_Station158_SUR_NODE_1364 | marine (Norwegian Sea) | 0 | 1 | 83 | 1 | 2399 | 2,398 | 0 | *Flavobacteriaceae* | Polaribacter vadi strain LPB0003 | C4-dicarboxylate ABC transporter | * |
| **Forsetiviridae** | GOV2_Station76_MES_NODE_1002 | marine (South Atlantic Ocean) | 0 | 0 | 100 | 47,019 | 47708 | 689 | 0 | *Flavobacteriaceae* | Maribacter cobaltidurans strain B1 | Paa1 family thioesterase | * |
| **Forsetiviridae** | GOV2_Station138_MES_NODE_1133 | marine (North Pacific Ocean) | 0 | 0 | 68 | 43,953 | 45601 | 1,648 | 8.68E-154 | *Flavobacteriaceae* | Elizabethkingia anophelis strain E6809 | hypothetical protein | * |
| **Forsetiviridae** | IMGVR2_3300009550_____Ga0115013_10000204 | marine (South Atlantic Ocean) | 0 | 21 | 81 | 21,694 | 21826 | 132 | 6.30E-21 | *Flavobacteriaceae* | Cellulophaga baltica 18 | hypothetical protein | * |
| **Forsetiviridae** | IMGVR2_3300009684_____Ga0114958_10000210 | freshwater lake | 0 | 0 | 79 | 44,824 | 46035 | 1,211 | 0 | *Flavobacteriaceae* | Flavobacterium psychrophilum strain 5 | chromosome partitioning protein ParB | * |
| **Forsetiviridae** | IMGVR2_3300019755_____Ga0193954_1000159 | Freshwater microbial mat | 0 | 1 | 72 | 1,120 | 1692 | 572 | 2.27E-77 | *Bacteroidetes* | Bacteroides intestinalis strain APC919/174 | DNA cytosine methyltransferase | * |
| **Pachyvirinae** | IMGVR2_3300001278_____BBAY75_10000041 | marine (macroalgae surface) | 0 | 0 | 74 | 26,456 | 26848 | 392 | 4.86E-57 | *Flavobacteriaceae* | Flavivirga eckloniae strain ECD14 | hypothetical protein | |
| **Pachyvirinae** | IMGVR2_3300005056_____Ga0071102_1000080 | marine (Atlantic Ocean) | 0 | 0 | 75 | 54,907 | 55242 | 335 | 3.45E-53 | *Flavobacteriaceae* | Formosa sp. PS13 | DUF3127 | * |
| **Winoviridae** | IMGVR2_3300007093_____Ga0104055_1000085 | human, oral | 0 | 1 | 71 | 18,546 | 19837 | 1,291 | 1.10E-162 | *Flavobacteriaceae* | Capnocytophaga sp. H2931 | hypothetical protein | * |
| **Winoviridae** | IMGVR2_3300007713_____Ga0105659_1000065 | human, oral | 0 | 3 | 72 | 3 | 971 | 968 | 1.74E-147 | *Flavobacteriaceae* | Chryseobacterium balustinum strain KC_1863 | ribosome biogenesis GTPase Der | * |
| **Winoviridae** | IMGVR2_3300006459_____Ga0100222_100241 | human, oral | 0 | 1 | 70 | 33,833 | 35432 | 1,599 | 0 | *Flavobacteriaceae* | Chryseobacterium sp. NBC122 | hypothetical protein | * |
| **Winoviridae** | IMGVR2_3300007126_____Ga0102717_100 | human, oral | 0 | 1 | 72 | 7 | 800 | 793 | 0 | *Flavobacteriaceae* | Elizabethkingia anophelis | integrase | |

| | | | | | | | | | | | | | |
|---|---|---|---|---|---|---|---|---|---|---|---|---|---|
| | 371 | | | | | | | | | | | | |
| **Winoviridae** | IMGVR2_3300008130_____Ga0114850_100310 | human, oral | 0 | 5 | 70 | 6,522 | 10322 | 3,800 | 0 | *Flavobacteriaceae* | Chryseobacterium sp. F5649 | 2-oxoglutarate dehydrogenase complex dihydrolipoyllysine-residue succinyltransferase | * |
| **Winoviridae** | IMGVR2_3300006742_____Ga0101805_100074 | human, oral | 0 | 1 | 81 | 3,882 | 5124 | 1,242 | 0 | *Flavobacteriaceae* | Chryseobacterium carnipullorum strain F9942 | ADP-forming succinate--CoA ligase subunit beta | * |
| **Winoviridae** | IMGVR2_3300012252_____Ga0122200_100146 | city subway wood | 0 | 0 | 74 | 800 | 3867 | 3,067 | 0 | *Flavobacteriaceae* | Flavobacterium columnare ATCC 49512 | cytosine-specific methyltransferase | |
| **Winoviridae** | IMGVR2_3300010054_____Ga0098069_100157 | marine (Subarctic Pacific Ocean) | 0 | 0 | 68 | 195 | 1711 | 1,516 | 2.03E-128 | *Flavobacteriaceae* | Flavobacterium columnare strain 94-081 | ABC transporter ATP-binding protein | * |
| **Winoviridae** | IMGVR2_3300019758_____Ga0193951_1000082 | Freshwater microbial mat | 0 | 0 | 89 | 1 | 1869 | 1,868 | 0 | *Flavobacteriaceae* | Tenacibaculum mesophilum strain DSM 13764 | DUF4870 | * |
| **Winoviridae** | IMGVR2_3300014204_____Ga0172381_10001225 | landfill leachate | 2 | 9 | 66 | 660 | 1247 | 587 | 3.57E-29 | *Flavobacteriaceae* | Chryseobacterium lactis strain KC_1864 | DNA adenine methylase | |
| **Helgolandvirinae** | IMGVR2_3300001122_____JGI12148J13107_100002 | marine (South Atlantic Ocean) | 1 | 28 | 70 | 29,050 | 29967 | 917 | 5.05E-105 | *Flavobacteriaceae* | Gillisia sp. Hel1_33_143 | hypothetical protein | * |
| **Helgolandvirinae** | IMGVR2_3300012032_____Ga0136554_1000067 | saline lake | 0 | 44 | 91 | 29,643 | 30564 | 921 | 0 | *Flavobacteriaceae* | Psychroflexus torquis ATCC 700755 | EndoU-type ribonuclease | * |
| **Helgolandvirinae** | IMGVR2_3300009508_____Ga0115567_10000451 | marine (Kabeltonne, North Sea) | 0 | 0 | 94 | 28,021 | 33865 | 5,844 | 0 | *Flavobacteriaceae* | Gillisia sp. Hel1_33_143 | hypothetical protein | * |
| **Helgolandvirinae** | GOV2_Station168_DCM_NODE_1833 | marine (Barents Sea) | 0 | 2 | 93 | 40,662 | 41265 | 603 | 0 | *Flavobacteriaceae* | Polaribacter sp. SA4-12 | hypothetical protein | * |
| **Helgolandvirinae** | IMGVR2_3300001605_____Draft_10001254 | waste water | 0 | 0 | 67 | 32,641 | 34090 | 1,449 | 1.58E-116 | *Flavobacteriaceae* | Tenacibaculum dicentrarchi strain AY7486TD | hypothetical protein | * |
| **Helgolandvirinae** | IMGVR2_3300005080_____Ga0069611_10000122 | waste water | 0 | 0 | 72 | 23,590 | 24439 | 849 | 1.00E-119 | *Flavobacteriaceae* | Weeksella virosa strain NCTC11634 | site-specific DNA methylase | * |
| **Helgolandvirinae** | IMGVR2_3300005080_____Ga0069611_10000213 | waste water | 0 | 0 | 71 | 17,787 | 18276 | 489 | 6.43E-58 | *Flavobacteriaceae* | Chryseobacterium sp. IHB B 17019 | hypothetical protein | |

| | | | | | | | | | | | | | |
|---|---|---|---|---|---|---|---|---|---|---|---|---|---|
| **Helgolandvirinae** | IMGVR2_3300015214 _____Ga0172382_10001576 | landfill leachate | 0 | 0 | 69 | 31,915 | 3402 0 | 2,105 | 0 | *Flavobacteriaceae* | Chryseobacterium gleum strain 3012STDY6944375 | modification methylase DpnIIA | |
| **Dunevirinae** | IMGVR2_3300008250 _____Ga0105354_1000171 | marine (Gulf of Mexico) | 0 | 0 | 75 | 2,531 | 4121 | 1,590 | 0 | *Flavobacteriaceae* | Owenweeksia hongkongensis DSM 17368 | putative transcriptional regulator with HTH domain | * |

**Table 6:** Special genomic features of the Flosverisviridae.

| genome_names | integrase | LuxR gene | AI-E2 transporter | BACON domain | HicA/B |
|---|---|---|---|---|---|
| IMGVR2_3300001122_____JGI12148J13107_100002 | - | + | - | + | - |
| IMGVR2_3300012032_____Ga0136554_1000067 | + | + | - | + | - |
| IMGVR2_3300009508_____Ga0115567_10000451 | - | + | + | - | - |
| GOV2_Station168_DCM_ALL_assembly_NODE_1833_length_41265_cov_31.413807 | + | + | - | + | - |
| Leef_1 | + | + | - | + | - |
| IMGVR2_3300001605_____Draft_10001254 | + | + | - | - | - |
| IMGVR2_3300005080_____Ga0069611_10000122 | + | - | - | - | + |
| IMGVR2_3300015214_____Ga0172382_10001576 | + | + | - | - | - |
| IMGVR2_3300005080_____Ga0069611_10000213 | - | - | - | - | - |
| KU599887.1_Flavobacterium_phage_2A | + | + | - | - | - |
| KC959568.1_Flavobacterium_phage_6H | + | + | - | - | - |
| KU599889.1_Flavobacterium_phage_1H | + | + | - | - | - |
| IMGVR2_3300008250_____Ga0105354_1000171 | + | + | - | - | - |
| Ingeline_1 | + | + | - | + | - |

**Table 7:** CRISPR spacer mismatches to phage genomes. For Calle_1, Harreka_1, Ingeline_1, Molly_1, and Omtje_1 no match was found.

| Bacterium | length of spacer | Danklef_1 | Freya_1 | Gundel_1 | Leef_1 | Nekkels_1 | Peternella_1 |
|---|---|---|---|---|---|---|---|
| HaHaR_3_91 | 30 | | | | | | 9 |
| HaHaR_3_91 | 30 | | 3 | | | | |
| HaHaR_3_91 | 30 | 8 | 8 | | | | |
| HaHaR_3_91 | 30 | | 0 | | | | |
| HaHaR_3_91 | 30 | | 3 | | | | |
| HaHaR_3_91 | 30 | 9 | 9 | | | 9 | |
| HaHaR_3_91 | 30 | | | 8 | | | |
| R2A056_3_33 | 31 | 1 | 1 | | | | |
| R2A056_3_33 | 30 | 3 | | | | | |
| R2A056_3_33 | 30 | 1 | | | | | |
| R2A056_3_33 | 30 | 8 | | | 1 | | |
| R2A056_3_33 | 30 | | | 9 | | | |

**Table 8:** Spacer hits from IMG/VR blast against the viral spacer database and the metagenomic spacer database.

| Phage_name | Family_name | Spacer BLAST | Metagenome_SPACER_blast |
|---|---|---|---|
| Calle_1 | Pervagovirinae | 0 | 0 |
| Colly_1 | Molycolviridae | 0 | 1 |
| Danklef_1 | Forsetiviridae | 0 | 0 |
| Freya_1 | Forsetiviridae | 0 | 0 |
| Gundel_1 | Pachyvirinae | 0 | 0 |
| Harreka_1 | Aggregaviridae | 0 | 0 |
| Ingeline_1 | Dunevirinae | 0 | 0 |
| Leef_1 | Helgolandvirinae | 0 | 0 |
| Molly_1 | Molycolviridae | 0 | 0 |
| Nekkels_1 | Assiduviridae | 0 | 8* |
| Omtje_1 | Obscuriviridae | 0 | 0 |
| Peternella_1 | Winoviridae | 0 | 0 |
| GOV2_Station82_SUR_COMBINED_FINAL_NODE_637_length_40711_cov_68.812279 | Aggregaviridae | 1 | 7 |
| IMGVR2_3300000271_____PBR_1000291 | Aggregaviridae | 0 | 0 |
| IMGVR2_3300006920_____Ga0070748_1000149 | Aggregaviridae | 0 | 4 |
| IMGVR2_3300007541_____Ga0099848_1000064 | Aggregaviridae | 0 | 2 |
| IMGVR2_3300009169_____Ga0105097_10000088 | Aggregaviridae | 0 | 0 |
| IMGVR2_3300009678_____Ga0105252_10000338 | Aggregaviridae | 0 | 0 |
| IMGVR2_3300010364_____Ga0134066_10000002 | Aggregaviridae | 0 | 0 |
| IMGVR2_3300012031_____Ga0136561_1000079 | Aggregaviridae | 0 | 0 |
| IMGVR2_3300012147_____Ga0136588_1000088 | Aggregaviridae | 0 | 0 |
| IMGVR2_3300012961_____Ga0164302_10000013 | Aggregaviridae | 0 | 6 |
| IMGVR2_3300014166_____Ga0134079_10000028 | Aggregaviridae | 0 | 0 |
| IMGVR2_3300018482_____Ga0066669_10000002 | Aggregaviridae | 0 | 0 |
| IMGVR2_3300019760_____Ga0193953_1000104 | Aggregaviridae | 0 | 0 |
| IMGVR2_3300008250_____Ga0105354_1000171 | Dunevirinae | 0 | 0 |
| GOV2_Station158_MES_ALL_assembly_NODE_1201_length_49284_cov_8.598245 | Forsetiviridae | 0 | 0 |
| GOV2_Station158_SUR_ALL_assembly_NODE_1364_length_50267_cov_12.211563 | Forsetiviridae | 0 | 1 |
| IMGVR2_3300009550_____Ga0115013_10000204 | Forsetiviridae | 0 | 21 |
| IMGVR2_3300009684_____Ga0114958_10000210 | Forsetiviridae | 0 | 0 |
| IMGVR2_3300019755_____Ga0193954_1000159 | Forsetiviridae | 0 | 1 |
| GOV2_Station168_DCM_ALL_assembly_NODE_1833_length_41265_cov_31.413807 | Helgolandvirinae | 0 | 2 |
| IMGVR2_3300001122_____JGI12148J13107_100002 | Helgolandvirinae | 1 | 28 |
| IMGVR2_3300001605_____Draft_10001254 | Helgolandvirinae | 0 | 0 |
| IMGVR2_3300005080_____Ga0069611_10000122 | Helgolandvirinae | 0 | 0 |
| IMGVR2_3300005080_____Ga0069611_10000213 | Helgolandvirinae | 0 | 0 |
| IMGVR2_3300009508_____Ga0115567_10000451 | Helgolandvirinae | 0 | 0 |
| IMGVR2_3300012032_____Ga0136554_1000067 | Helgolandvirinae | 0 | 44 |
| IMGVR2_3300015214_____Ga0172382_10001576 | Helgolandvirinae | 0 | 0 |
| IMGVR2_3300001278_____BBAY75_10000041 | Pachyvirinae | 0 | 0 |
| IMGVR2_3300005056_____Ga0071102_1000080 | Pachyvirinae | 0 | 0 |
| IMGVR2_3300006459_____Ga0100222_100241 | Winoviridae | 0 | 1 |
| IMGVR2_3300006742_____Ga0101805_100074 | Winoviridae | 0 | 1 |
| IMGVR2_3300007093_____Ga0104055_1000085 | Winoviridae | 0 | 1 |
| IMGVR2_3300007126_____Ga0102717_100371 | Winoviridae | 0 | 1 |
| IMGVR2_3300007713_____Ga0105659_1000065 | Winoviridae | 0 | 3 |
| IMGVR2_3300008130_____Ga0114850_100310 | Winoviridae | 0 | 5 |
| IMGVR2_3300010054_____Ga0098069_100157 | Winoviridae | 0 | 0 |
| IMGVR2_3300012252_____Ga0122200_100146 | Winoviridae | 0 | 0 |
| IMGVR2_3300014204_____Ga0172381_10001225 | Winoviridae | 2 | 9 |
| IMGVR2_3300019758_____Ga0193951_1000082 | Winoviridae | 0 | 0 |

* the viral spacer IDs: 3300027009:Ga0209093_1001256:1:1555, 3300027262:Ga0209303_1000810:1:15572, 3300027498:Ga0209185_1193139:1:249, 3300009417:Ga0114953_1000198:1:2830, 3300009421:Ga0114952_1377632:1:220, 3300009415:Ga0115029_1000158:2:63503, 3300009072:Ga0115030_1000872:1:653, 3300027325:Ga0209186_1000073:1:10669

**Table 9:** Read mapping results of phage Danklef and its host.

| Danklef | | | | | | host 95% identity | | | | | phage 70% identity | | | | | phage 100% identity | | | | |
|---|---|---|---|---|---|---|---|---|---|---|---|---|---|---|---|---|---|---|---|---|
| metagenome | date | Julian Day | total reads | total bases | size fraction | genome coverage [%] | mapped reads | total bases reads | relative read abundance [%] | normalized coverage [%] | genome coverage [%] | mapped reads | total bases reads | relative read abundance [%] | normalized coverage [%] | genome coverage [%] | mapped reads | total bases reads | relative read abundance [%] | normalized coverage [%] |
| | 19.03.2018 | 78 | 132647062 | 3.216E+10 | 0.2 -3 µm | 9.43 | 4994 | 1188625 | 0.0038 | 0.0085 | 5.03 | 22 | 5334 | 0.0000 | 0.0035 | 0.84 | 2 | 499 | 0.0000 | 0.0003 |
| | 03.04.2018 | 93 | 130734852 | 3.094E+10 | 0.2 -3 µm | 54.99 | 24581 | 5897035 | 0.0188 | 0.0440 | 36.11 | 165 | 39487 | 0.0001 | 0.0270 | 6.69 | 21 | 4745 | 0.0000 | 0.0033 |
| | 05.04.2018 | 95 | 130836262 | 3.131E+10 | 0.2 -3 µm | 12.02 | 7086 | 1666568 | 0.0054 | 0.0123 | 5.75 | 20 | 4901 | 0.0000 | 0.0033 | 1.15 | 4 | 942 | 0.0000 | 0.0006 |
| | 10.04.2018 | 100 | 131511730 | 3.167E+10 | 0.2 -3 µm | 13.20 | 6380 | 1513284 | 0.0049 | 0.0110 | 4.80 | 17 | 4254 | 0.0000 | 0.0028 | 1.34 | 4 | 994 | 0.0000 | 0.0007 |
| | 12.04.2018 | 102 | 137437816 | 3.31E+10 | 0.2 -3 µm | 14.19 | 6571 | 1566388 | 0.0048 | 0.0109 | 5.68 | 35 | 8393 | 0.0000 | 0.0054 | NA | 0 | 0 | 0.0000 | 0.0000 |
| | 17.04.2018 | 108 | 134007214 | 3.216E+10 | 0.2 -3 µm | 23.07 | 11848 | 2800784 | 0.0088 | 0.0201 | 6.68 | 28 | 6956 | 0.0000 | 0.0046 | 0.67 | 2 | 500 | 0.0000 | 0.0003 |
| | 19.04.2018 | 110 | 145428602 | 3.516E+10 | 0.2 -3 µm | 2.49 | 5360 | 1262124 | 0.0037 | 0.0083 | 3.38 | 9 | 2210 | 0.0000 | 0.0013 | NA | 0 | 0 | 0.0000 | 0.0000 |
| | 24.04.2018 | 115 | 148226228 | 3.577E+10 | 0.2 -3 µm | 9.04 | 6485 | 1536719 | 0.0044 | 0.0099 | 4.14 | 27 | 6038 | 0.0000 | 0.0036 | 1.80 | 6 | 1199 | 0.0000 | 0.0007 |
| | 26.04.2018 | 117 | 186004950 | 4.485E+10 | 0.2 -3 µm | 17.38 | 14495 | 3445399 | 0.0078 | 0.0177 | 8.88 | 118 | 28607 | 0.0001 | 0.0135 | 0.53 | 1 | 251 | 0.0000 | 0.0001 |
| | 02.05.2018 | 123 | 172417734 | 4.171E+10 | 0.2 -3 µm | 69.26 | 50034 | 11884495 | 0.0290 | 0.0658 | 29.52 | 285 | 67909 | 0.0002 | 0.0345 | 3.67 | 8 | 1944 | 0.0000 | 0.0010 |
| | 03.05.2018 | 124 | 175408924 | 4.239E+10 | 0.2 -3 µm | 16.15 | 14667 | 3490138 | 0.0084 | 0.0190 | 5.90 | 81 | 19063 | 0.0000 | 0.0095 | NA | 0 | 0 | 0.0000 | 0.0000 |
| | 08.05.2018 | 129 | 157697368 | 3.823E+10 | 0.2 -3 µm | 4.09 | 9794 | 2315843 | 0.0062 | 0.0140 | 3.93 | 70 | 16889 | 0.0000 | 0.0094 | NA | 0 | 0 | 0.0000 | 0.0000 |
| | 11.05.2018 | 134 | 174367208 | 4.196E+10 | 0.2 -3 µm | 2.19 | 7828 | 1837712 | 0.0045 | 0.0101 | 2.19 | 40 | 9605 | 0.0000 | 0.0049 | NA | 0 | 0 | 0.0000 | 0.0000 |
| | 15.05.2018 | 136 | 167144802 | 4.036E+10 | 0.2 -3 µm | 1.68 | 6884 | 1614687 | 0.0041 | 0.0092 | 2.76 | 31 | 7690 | 0.0000 | 0.0040 | NA | 0 | 0 | 0.0000 | 0.0000 |
| | 17.05.2018 | 142 | 178481766 | 4.291E+10 | 0.2 -3 µm | 2.54 | 6735 | 1581558 | 0.0038 | 0.0085 | 2.46 | 39 | 9563 | 0.0000 | 0.0047 | NA | 0 | 0 | 0.0000 | 0.0000 |
| | 22.05.2018 | 143 | 162504646 | 3.904E+10 | 0.2 -3 µm | 45.15 | 23120 | 5492681 | 0.0142 | 0.0325 | 2.65 | 34 | 8124 | 0.0000 | 0.0044 | NA | 0 | 0 | 0.0000 | 0.0000 |
| | 24.05.2018 | 145 | 159197066 | 3.831E+10 | 0.2 -3 µm | 11.96 | 11956 | 2806414 | 0.0075 | 0.0169 | 2.35 | 20 | 4787 | 0.0000 | 0.0026 | NA | 0 | 0 | 0.0000 | 0.0000 |
| | 29.05.2018 | 150 | 171271466 | 4.143E+10 | 0.2 -3 µm | 9.28 | 8498 | 2006371 | 0.0050 | 0.0112 | 3.34 | 35 | 8514 | 0.0000 | 0.0044 | NA | 0 | 0 | 0.0000 | 0.0000 |
| | 19.03.2018 | 78 | 408373164 | 9.731E+10 | 3-10 µm | 27.12 | 12534 | 2990531 | 0.0031 | 0.0071 | 22.10 | 91 | 21556 | 0.0000 | 0.0047 | 1.90 | 7 | 1427 | 0.0000 | 0.0003 |
| | 12.04.2018 | 102 | 332059730 | 7.927E+10 | 3-10 µm | 23.33 | 11040 | 2641373 | 0.0033 | 0.0077 | 21.81 | 91 | 21294 | 0.0000 | 0.0057 | 4.00 | 13 | 2844 | 0.0000 | 0.0008 |
| | 17.04.2018 | 108 | 420871688 | 9.615E+10 | 3-10 µm | 24.59 | 16273 | 3816006 | 0.0039 | 0.0092 | 11.67 | 71 | 16859 | 0.0000 | 0.0037 | 2.35 | 11 | 2583 | 0.0000 | 0.0006 |
| | 26.04.2018 | 117 | 341246096 | 8.148E+10 | 3-10 µm | 29.41 | 15328 | 3672789 | 0.0045 | 0.0104 | 24.53 | 158 | 36915 | 0.0000 | 0.0096 | 3.45 | 8 | 1907 | 0.0000 | 0.0005 |
| | 08.05.2018 | 129 | 390032052 | 8.751E+10 | 3-10 µm | 2.66 | 4425 | 980248 | 0.0011 | 0.0026 | 9.37 | 50 | 11268 | 0.0000 | 0.0027 | 2.17 | 10 | 2089 | 0.0000 | 0.0005 |
| | 11.05.2018 | 134 | 346400002 | 8.256E+10 | 3-10 µm | 2.02 | 2851 | 678199 | 0.0008 | 0.0019 | 19.02 | 109 | 25778 | 0.0000 | 0.0066 | 3.90 | 11 | 2662 | 0.0000 | 0.0007 |
| | 22.05.2018 | 143 | 322898796 | 7.6E+10 | 3-10 µm | 50.62 | 78041 | 18478370 | 0.0242 | 0.0562 | 3.98 | 97 | 22638 | 0.0000 | 0.0063 | 0.53 | 5 | 1215 | 0.0000 | 0.0003 |

| | | | | | | | | | | | | | | | | | | | |
|---|---|---|---|---|---|---|---|---|---|---|---|---|---|---|---|---|---|---|---|
| 29.05.2018 | 150 | 361105792 | 8.634E+10 | 3-10 µm | 9.71 | 5784 | 1380526 | 0.0016 | 0.0037 | 16.49 | 66 | 14788 | 0.0000 | 0.0036 | 4.06 | 15 | 3227 | 0.0000 | 0.0008 |
| 19.03.2018 | 78 | 229237742 | 5.467E+10 | >10 µm | 2.94 | 7886 | 1838420 | 0.0034 | 0.0078 | 6.65 | 26 | 5165 | 0.0000 | 0.0020 | 4.17 | 16 | 2936 | 0.0000 | 0.0011 |
| 12.04.2018 | 102 | 366243118 | 8.751E+10 | >10 µm | 89.55 | 221392 | 53121088 | 0.0604 | 0.1402 | 98.42 | 790340 | 186175735 | 0.2158 | 45.0849 | 85.36 | 180645 | 42088011 | 0.0493 | 10.1922 |
| 17.04.2018 | 108 | 349480760 | 8.224E+10 | >10 µm | 7.10 | 19387 | 4643716 | 0.0055 | 0.0130 | 17.20 | 92 | 21466 | 0.0000 | 0.0055 | 4.38 | 14 | 3222 | 0.0000 | 0.0008 |
| 26.04.2018 | 117 | 357085094 | 8.282E+10 | >10 µm | 84.44 | 93975 | 21978405 | 0.0263 | 0.0613 | 11.02 | 492 | 114141 | 0.0001 | 0.0292 | 0.56 | 4 | 524 | 0.0000 | 0.0001 |
| 08.05.2018 | 129 | 332293300 | 7.447E+10 | >10 µm | 10.03 | 10802 | 2379436 | 0.0033 | 0.0074 | 9.18 | 156 | 34633 | 0.0000 | 0.0099 | 3.40 | 16 | 3250 | 0.0000 | 0.0009 |
| 11.05.2018 | 134 | 340848252 | 8.026E+10 | >10 µm | 3.83 | 3193 | 758983 | 0.0009 | 0.0022 | 14.13 | 68 | 15466 | 0.0000 | 0.0041 | 3.02 | 11 | 2241 | 0.0000 | 0.0006 |
| 22.05.2018 | 143 | 416648526 | 9.496E+10 | >10 µm | 97.97 | 780276 | 181559855 | 0.1873 | 0.4416 | 67.39 | 628 | 145880 | 0.0002 | 0.0326 | 15.34 | 51 | 11736 | 0.0000 | 0.0026 |
| 29.05.2018 | 150 | 369290526 | 7.388E+10 | >10 µm | 46.66 | 56043 | 11208341 | 0.0152 | 0.0350 | 9.53 | 144 | 28872 | 0.0000 | 0.0083 | 3.22 | 19 | 4129 | 0.0000 | 0.0012 |

**Table 10:** Read mapping results of phage Freya and its host

| Freya | | | | | | host 95% identity | | | | | phage 70% identity | | | | | phage 100% identity | | | | |
|---|---|---|---|---|---|---|---|---|---|---|---|---|---|---|---|---|---|---|---|---|
| metagenome | date | Julian Day | total reads | total bases | size fraction | genome coverage [%] | mapped reads | total bases reads | relative read abundance [%] | normalized coverage [%] | genome coverage [%] | mapped reads | total bases reads | relative read abundance [%] | normalized coverage [%] | genome coverage [%] | mapped reads | total bases reads | relative read abundance [%] | normalized coverage [%] |
| | 19.03.2018 | 78 | 132647062 | 3.216E+10 | 0.2 -3 µm | 9.77 | 5004 | 1190814 | 0.0038 | 0.0088 | 4.06 | 20 | 4802 | 0.0000 | 0.0034 | NA | 0 | 0 | 0.0000 | 0.0000 |
| | 03.04.2018 | 93 | 130734852 | 3.094E+10 | 0.2 -3 µm | 56.31 | 24590 | 5899122 | 0.0188 | 0.0453 | 41.04 | 179 | 42572 | 0.0001 | 0.0313 | 15.56 | 46 | 10810 | 0.0000 | 0.0079 |
| | 05.04.2018 | 95 | 130836262 | 3.131E+10 | 0.2 -3 µm | 12.44 | 7098 | 1668773 | 0.0054 | 0.0127 | 6.92 | 21 | 5163 | 0.0000 | 0.0037 | 1.81 | 4 | 954 | 0.0000 | 0.0007 |
| | 10.04.2018 | 100 | 131511730 | 3.167E+10 | 0.2 -3 µm | 13.63 | 6386 | 1515156 | 0.0049 | 0.0114 | 5.47 | 17 | 4253 | 0.0000 | 0.0031 | 2.01 | 5 | 1245 | 0.0000 | 0.0009 |
| | 12.04.2018 | 102 | 137437816 | 3.31E+10 | 0.2 -3 µm | 14.67 | 6574 | 1566772 | 0.0048 | 0.0112 | 5.69 | 34 | 8332 | 0.0000 | 0.0057 | NA | 0 | 0 | 0.0000 | 0.0000 |
| | 17.04.2018 | 108 | 134007214 | 3.216E+10 | 0.2 -3 µm | 23.74 | 11865 | 2805163 | 0.0089 | 0.0207 | 5.53 | 26 | 6463 | 0.0000 | 0.0046 | 1.14 | 2 | 501 | 0.0000 | 0.0004 |
| | 19.04.2018 | 110 | 145428602 | 3.516E+10 | 0.2 -3 µm | 2.52 | 5337 | 1256293 | 0.0037 | 0.0085 | 2.40 | 8 | 1974 | 0.0000 | 0.0013 | 0.55 | 1 | 243 | 0.0000 | 0.0002 |
| | 24.04.2018 | 115 | 148226228 | 3.577E+10 | 0.2 -3 µm | 9.41 | 6514 | 1543024 | 0.0044 | 0.0103 | 4.28 | 34 | 7607 | 0.0000 | 0.0048 | 0.95 | 3 | 587 | 0.0000 | 0.0004 |
| | 26.04.2018 | 117 | 186004950 | 4.485E+10 | 0.2 -3 µm | 17.92 | 14636 | 3480717 | 0.0079 | 0.0184 | 9.29 | 109 | 26337 | 0.0001 | 0.0134 | 3.19 | 7 | 1624 | 0.0000 | 0.0008 |
| | 02.05.2018 | 123 | 172417734 | 4.171E+10 | 0.2 -3 µm | 70.63 | 49858 | 11843314 | 0.0289 | 0.0675 | 27.45 | 272 | 64448 | 0.0002 | 0.0351 | 9.76 | 24 | 5764 | 0.0000 | 0.0031 |
| | 03.05.2018 | 124 | 175408924 | 4.239E+10 | 0.2 -3 µm | 16.87 | 14788 | 3519557 | 0.0084 | 0.0197 | 6.96 | 88 | 20730 | 0.0001 | 0.0111 | 0.86 | 2 | 498 | 0.0000 | 0.0003 |
| | 08.05.2018 | 129 | 157697368 | 3.823E+10 | 0.2 -3 µm | 4.28 | 9831 | 2324291 | 0.0062 | 0.0144 | 3.65 | 59 | 14136 | 0.0000 | 0.0084 | NA | 0 | 0 | 0.0000 | 0.0000 |
| | 11.05.2018 | 134 | 174367208 | 4.196E+10 | 0.2 -3 µm | 2.30 | 7838 | 1839997 | 0.0045 | 0.0104 | 2.36 | 45 | 10759 | 0.0000 | 0.0058 | NA | 0 | 0 | 0.0000 | 0.0000 |
| | 15.05.2018 | 136 | 167144802 | 4.036E+10 | 0.2 -3 µm | 1.76 | 6901 | 1617856 | 0.0041 | 0.0095 | 1.88 | 32 | 7889 | 0.0000 | 0.0044 | NA | 0 | 0 | 0.0000 | 0.0000 |
| | 17.05.2018 | 142 | 178481766 | 4.291E+10 | 0.2 -3 µm | 2.67 | 6766 | 1589226 | 0.0038 | 0.0088 | 2.64 | 36 | 8918 | 0.0000 | 0.0047 | NA | 0 | 0 | 0.0000 | 0.0000 |
| | 22.05.2018 | 143 | 162504646 | 3.904E+10 | 0.2 -3 µm | 46.23 | 23108 | 5488630 | 0.0142 | 0.0334 | 2.18 | 31 | 7265 | 0.0000 | 0.0042 | NA | 0 | 0 | 0.0000 | 0.0000 |

| | | | | | | | | | | | | | | | | | | | |
|---|---|---|---|---|---|---|---|---|---|---|---|---|---|---|---|---|---|---|---|
| 24.05.2018 | 145 | 159197066 | 3.831E+10 | 0.2 -3 µm | 12.19 | 11947 | 2803060 | 0.0075 | 0.0174 | 2.52 | 17 | 4035 | 0.0000 | 0.0024 | NA | 0 | 0 | 0.0000 | 0.0000 |
| 29.05.2018 | 150 | 171271466 | 4.143E+10 | 0.2 -3 µm | 9.68 | 8528 | 2014164 | 0.0050 | 0.0116 | 4.15 | 36 | 8765 | 0.0000 | 0.0048 | 0.56 | 1 | 248 | 0.0000 | 0.0001 |
| 19.03.2018 | 78 | 408373164 | 9.731E+10 | 3-10 µm | 28.03 | 12535 | 2992590 | 0.0031 | 0.0073 | 23.33 | 87 | 20723 | 0.0000 | 0.0048 | 6.23 | 19 | 4286 | 0.0000 | 0.0010 |
| 12.04.2018 | 102 | 332059730 | 7.927E+10 | 3-10 µm | 23.93 | 11013 | 2630933 | 0.0033 | 0.0079 | 30.07 | 113 | 26470 | 0.0000 | 0.0076 | 20.53 | 62 | 14457 | 0.0000 | 0.0041 |
| 17.04.2018 | 108 | 420871688 | 9.615E+10 | 3-10 µm | 25.34 | 16327 | 3823644 | 0.0039 | 0.0094 | 15.33 | 89 | 21284 | 0.0000 | 0.0050 | 6.77 | 23 | 5412 | 0.0000 | 0.0013 |
| 26.04.2018 | 117 | 341246096 | 8.148E+10 | 3-10 µm | 29.92 | 15261 | 3654928 | 0.0045 | 0.0107 | 31.34 | 176 | 40872 | 0.0001 | 0.0114 | 21.11 | 65 | 14571 | 0.0000 | 0.0041 |
| 08.05.2018 | 129 | 390032052 | 8.751E+10 | 3-10 µm | 2.79 | 4626 | 1017508 | 0.0012 | 0.0028 | 11.30 | 70 | 15650 | 0.0000 | 0.0041 | 7.94 | 38 | 8572 | 0.0000 | 0.0022 |
| 11.05.2018 | 134 | 346400002 | 8.256E+10 | 3-10 µm | 2.15 | 3015 | 704747 | 0.0009 | 0.0020 | 25.64 | 131 | 31052 | 0.0000 | 0.0086 | 21.64 | 74 | 17316 | 0.0000 | 0.0048 |
| 22.05.2018 | 143 | 322898796 | 7.6E+10 | 3-10 µm | 51.84 | 77576 | 18356831 | 0.0240 | 0.0574 | 3.79 | 135 | 31736 | 0.0000 | 0.0095 | 1.02 | 9 | 2219 | 0.0000 | 0.0007 |
| 29.05.2018 | 150 | 361105792 | 8.634E+10 | 3-10 µm | 9.94 | 5781 | 1377963 | 0.0016 | 0.0038 | 19.02 | 77 | 17065 | 0.0000 | 0.0045 | 15.86 | 52 | 11517 | 0.0000 | 0.0030 |
| 19.03.2018 | 78 | 229237742 | 5.467E+10 | >10 µm | 3.05 | 7871 | 1833101 | 0.0034 | 0.0080 | 5.03 | 44 | 9492 | 0.0000 | 0.0039 | 0.60 | 4 | 523 | 0.0000 | 0.0002 |
| 12.04.2018 | 102 | 366243118 | 8.751E+10 | >10 µm | 91.68 | 221532 | 53163739 | 0.0605 | 0.1444 | 100.00 | 1017402 | 239788612 | 0.2778 | 62.3038 | 100.00 | 862986 | 203769042 | 0.2356 | 52.9449 |
| 17.04.2018 | 108 | 349480760 | 8.224E+10 | >10 µm | 7.27 | 19439 | 4639441 | 0.0056 | 0.0134 | 25.67 | 128 | 29918 | 0.0000 | 0.0083 | 20.06 | 65 | 14670 | 0.0000 | 0.0041 |
| 26.04.2018 | 117 | 357085094 | 8.282E+10 | >10 µm | 85.92 | 93420 | 21843750 | 0.0262 | 0.0627 | 8.02 | 511 | 119352 | 0.0001 | 0.0328 | NA | 0 | 0 | 0.0000 | 0.0000 |
| 08.05.2018 | 129 | 332293300 | 7.447E+10 | >10 µm | 10.26 | 10703 | 2357850 | 0.0032 | 0.0075 | 10.20 | 151 | 33976 | 0.0000 | 0.0104 | 2.58 | 11 | 2309 | 0.0000 | 0.0007 |
| 11.05.2018 | 134 | 340848252 | 8.026E+10 | >10 µm | 3.77 | 3202 | 755997 | 0.0009 | 0.0022 | 18.26 | 83 | 18751 | 0.0000 | 0.0053 | 15.61 | 52 | 11565 | 0.0000 | 0.0033 |
| 22.05.2018 | 143 | 416648526 | 9.496E+10 | >10 µm | 97.83 | 777821 | 180995463 | 0.1867 | 0.4529 | 77.85 | 656 | 152642 | 0.0002 | 0.0365 | 38.06 | 137 | 31079 | 0.0000 | 0.0074 |
| 29.05.2018 | 150 | 369290526 | 7.388E+10 | >10 µm | 47.44 | 55357 | 11074665 | 0.0150 | 0.0356 | 9.31 | 139 | 28185 | 0.0000 | 0.0087 | 5.23 | 30 | 6314 | 0.0000 | 0.0019 |

**Table 11:** Read mapping results of phage Harreka and its host

| Harreka | | | | | | host 95% identity | | | | | phage 70% identity | | | | | phage 100% identity | | | | |
|---|---|---|---|---|---|---|---|---|---|---|---|---|---|---|---|---|---|---|---|---|
| metagenome | date | Julian Day | total reads | total bases | size fraction | genome coverage [%] | mapped reads | total bases reads | relative read abundance [%] | normalized coverage [%] | genome coverage [%] | mapped reads | total bases reads | relative read abundance [%] | normalized coverage [%] | genome coverage [%] | mapped reads | total bases reads | relative read abundance [%] | normalized coverage [%] |
| | 19.03.2018 | 78 | 132647062 | 3.216E+10 | 0.2 -3 µm | 5.69 | 4202 | 996396 | 0.0032 | 0.0070 | 4.50 | 12 | 2778 | 0.0000 | 0.0020 | 1.26 | 3 | 751 | 0.0000 | 0.0005 |
| | 03.04.2018 | 93 | 130734852 | 3.094E+10 | 0.2 -3 µm | 26.98 | 12212 | 2905325 | 0.0093 | 0.0213 | 36.91 | 127 | 30850 | 0.0001 | 0.0231 | 0.98 | 2 | 501 | 0.0000 | 0.0004 |
| | 05.04.2018 | 95 | 130836262 | 3.131E+10 | 0.2 -3 µm | 3.03 | 5394 | 1251525 | 0.0041 | 0.0091 | 0.79 | 2 | 501 | 0.0000 | 0.0004 | NA | 0 | 0 | 0.0000 | 0.0000 |
| | 10.04.2018 | 100 | 131511730 | 3.167E+10 | 0.2 -3 µm | 4.50 | 4365 | 1026578 | 0.0033 | 0.0074 | 0.61 | 2 | 496 | 0.0000 | 0.0004 | NA | 0 | 0 | 0.0000 | 0.0000 |
| | 12.04.2018 | 102 | 137437816 | 3.31E+10 | 0.2 -3 µm | 4.90 | 4379 | 1035734 | 0.0032 | 0.0071 | 1.61 | 4 | 1003 | 0.0000 | 0.0007 | NA | 0 | 0 | 0.0000 | 0.0000 |
| | 17.04.2018 | 108 | 134007214 | 3.216E+10 | 0.2 -3 µm | 8.91 | 7223 | 1694746 | 0.0054 | 0.0119 | 0.96 | 2 | 416 | 0.0000 | 0.0003 | NA | 0 | 0 | 0.0000 | 0.0000 |
| | 19.04.2018 | 110 | 145428602 | 3.516E+10 | 0.2 -3 µm | 0.94 | 5141 | 1208712 | 0.0035 | 0.0078 | 1.15 | 2 | 501 | 0.0000 | 0.0003 | NA | 0 | 0 | 0.0000 | 0.0000 |

| | | | | | | | | | | | | | | | | | | | |
|---|---|---|---|---|---|---|---|---|---|---|---|---|---|---|---|---|---|---|---|
| 24.04.2018 | 115 | 148226228 | 3.577E+10 | 0.2 -3 µm | 4.55 | 5030 | 1187657 | 0.0034 | 0.0075 | NA | 0 | 0 | 0.0000 | 0.0000 | NA | 0 | 0 | 0.0000 | 0.0000 |
| 26.04.2018 | 117 | 186004950 | 4.485E+10 | 0.2 -3 µm | 8.06 | 9685 | 2288183 | 0.0052 | 0.0116 | 2.22 | 4 | 957 | 0.0000 | 0.0005 | NA | 0 | 0 | 0.0000 | 0.0000 |
| 02.05.2018 | 123 | 172417734 | 4.171E+10 | 0.2 -3 µm | 59.11 | 34669 | 8195312 | 0.0201 | 0.0446 | 11.15 | 29 | 7107 | 0.0000 | 0.0039 | 1.81 | 4 | 954 | 0.0000 | 0.0005 |
| 03.05.2018 | 124 | 175408924 | 4.239E+10 | 0.2 -3 µm | 9.21 | 12262 | 2909762 | 0.0070 | 0.0156 | 2.25 | 8 | 1905 | 0.0000 | 0.0010 | NA | 0 | 0 | 0.0000 | 0.0000 |
| 08.05.2018 | 129 | 157697368 | 3.823E+10 | 0.2 -3 µm | 2.25 | 11830 | 2793998 | 0.0075 | 0.0166 | NA | 0 | 0 | 0.0000 | 0.0000 | NA | 0 | 0 | 0.0000 | 0.0000 |
| 11.05.2018 | 134 | 174367208 | 4.196E+10 | 0.2 -3 µm | 1.56 | 10251 | 2401621 | 0.0059 | 0.0130 | NA | 0 | 0 | 0.0000 | 0.0000 | NA | 0 | 0 | 0.0000 | 0.0000 |
| 15.05.2018 | 136 | 167144802 | 4.036E+10 | 0.2 -3 µm | 1.24 | 9985 | 2346676 | 0.0060 | 0.0132 | 0.58 | 1 | 251 | 0.0000 | 0.0001 | NA | 0 | 0 | 0.0000 | 0.0000 |
| 17.05.2018 | 142 | 178481766 | 4.291E+10 | 0.2 -3 µm | 1.74 | 8796 | 2062928 | 0.0049 | 0.0109 | 2.52 | 5 | 1236 | 0.0000 | 0.0007 | NA | 0 | 0 | 0.0000 | 0.0000 |
| 22.05.2018 | 143 | 162504646 | 3.904E+10 | 0.2 -3 µm | 3.67 | 9578 | 2236473 | 0.0059 | 0.0130 | 1.16 | 2 | 502 | 0.0000 | 0.0003 | NA | 0 | 0 | 0.0000 | 0.0000 |
| 24.05.2018 | 145 | 159197066 | 3.831E+10 | 0.2 -3 µm | 1.59 | 10564 | 2468194 | 0.0066 | 0.0146 | 1.41 | 5 | 974 | 0.0000 | 0.0006 | NA | 0 | 0 | 0.0000 | 0.0000 |
| 29.05.2018 | 150 | 171271466 | 4.143E+10 | 0.2 -3 µm | 1.47 | 7214 | 1694820 | 0.0042 | 0.0093 | 0.99 | 5 | 997 | 0.0000 | 0.0006 | NA | 0 | 0 | 0.0000 | 0.0000 |
| 19.03.2018 | 78 | 408373164 | 9.731E+10 | 3-10 µm | 11.63 | 7308 | 1740784 | 0.0018 | 0.0041 | 10.18 | 44 | 8804 | 0.0000 | 0.0021 | 1.98 | 6 | 1177 | 0.0000 | 0.0003 |
| 12.04.2018 | 102 | 332059730 | 7.927E+10 | 3-10 µm | 6.23 | 4417 | 1057338 | 0.0013 | 0.0030 | 9.01 | 24 | 5829 | 0.0000 | 0.0017 | 0.58 | 1 | 251 | 0.0000 | 0.0001 |
| 17.04.2018 | 108 | 420871688 | 9.615E+10 | 3-10 µm | 12.11 | 8315 | 1939600 | 0.0020 | 0.0046 | 8.15 | 36 | 7691 | 0.0000 | 0.0019 | 0.54 | 1 | 234 | 0.0000 | 0.0001 |
| 26.04.2018 | 117 | 341246096 | 8.148E+10 | 3-10 µm | 20.38 | 9099 | 2186167 | 0.0027 | 0.0061 | 7.10 | 23 | 5755 | 0.0000 | 0.0016 | NA | 0 | 0 | 0.0000 | 0.0000 |
| 08.05.2018 | 129 | 390032052 | 8.751E+10 | 3-10 µm | 1.62 | 4521 | 1015272 | 0.0012 | 0.0026 | 100.00 | 128332 | 21163852 | 0.0329 | 5.6013 | 100.00 | 41276 | 6833569 | 0.0106 | 1.8086 |
| 11.05.2018 | 134 | 346400002 | 8.256E+10 | 3-10 µm | 1.72 | 4522 | 1081727 | 0.0013 | 0.0030 | 0.77 | 3 | 751 | 0.0000 | 0.0002 | NA | 0 | 0 | 0.0000 | 0.0000 |
| 22.05.2018 | 143 | 322898796 | 7.6E+10 | 3-10 µm | 3.44 | 10539 | 2467150 | 0.0033 | 0.0074 | 1.98 | 9 | 2025 | 0.0000 | 0.0006 | NA | 0 | 0 | 0.0000 | 0.0000 |
| 29.05.2018 | 150 | 361105792 | 8.634E+10 | 3-10 µm | 1.71 | 4993 | 1193566 | 0.0014 | 0.0031 | 1.96 | 4 | 958 | 0.0000 | 0.0003 | NA | 0 | 0 | 0.0000 | 0.0000 |
| 19.03.2018 | 78 | 229237742 | 5.467E+10 | >10 µm | 1.31 | 4438 | 1026359 | 0.0019 | 0.0043 | 1.16 | 28 | 6675 | 0.0000 | 0.0028 | NA | 0 | 0 | 0.0000 | 0.0000 |
| 12.04.2018 | 102 | 366243118 | 8.751E+10 | >10 µm | 58.53 | 71613 | 17163734 | 0.0196 | 0.0445 | 3.25 | 11 | 2390 | 0.0000 | 0.0006 | 0.32 | 2 | 275 | 0.0000 | 0.0001 |
| 17.04.2018 | 108 | 349480760 | 8.224E+10 | >10 µm | 3.90 | 11635 | 2777313 | 0.0033 | 0.0077 | 2.62 | 58 | 14207 | 0.0000 | 0.0040 | NA | 0 | 0 | 0.0000 | 0.0000 |
| 26.04.2018 | 117 | 357085094 | 8.282E+10 | >10 µm | 63.36 | 43395 | 10135769 | 0.0122 | 0.0278 | 10.76 | 35 | 7517 | 0.0000 | 0.0021 | NA | 0 | 0 | 0.0000 | 0.0000 |
| 08.05.2018 | 129 | 332293300 | 7.447E+10 | >10 µm | 6.60 | 7271 | 1600647 | 0.0022 | 0.0049 | 4.22 | 17 | 3648 | 0.0000 | 0.0011 | NA | 0 | 0 | 0.0000 | 0.0000 |
| 11.05.2018 | 134 | 340848252 | 8.026E+10 | >10 µm | 4.39 | 3577 | 858813 | 0.0010 | 0.0024 | 2.31 | 9 | 1564 | 0.0000 | 0.0005 | 0.36 | 2 | 312 | 0.0000 | 0.0001 |
| 22.05.2018 | 143 | 416648526 | 9.496E+10 | >10 µm | 44.62 | 30459 | 6982053 | 0.0073 | 0.0167 | 3.29 | 8 | 1841 | 0.0000 | 0.0004 | NA | 0 | 0 | 0.0000 | 0.0000 |
| 29.05.2018 | 150 | 369290526 | 7.388E+10 | >10 µm | 3.82 | 11398 | 2253368 | 0.0031 | 0.0069 | 2.66 | 35 | 6355 | 0.0000 | 0.0020 | NA | 0 | 0 | 0.0000 | 0.0000 |

**Table 12:** Read mapping results of phage Ingeline and its host

| Ingeline | | | | | | host 95% identity | | | | | phage 70% identity | | | | | phage 100% identity | | | | |
|---|---|---|---|---|---|---|---|---|---|---|---|---|---|---|---|---|---|---|---|---|
| metagenome | date | Julian Day | total reads | total bases | size fraction | genome coverage [%] | mapped reads | total bases reads | relative read abundance [%] | normalized coverage [%] | genome coverage [%] | mapped reads | total bases reads | relative read abundance [%] | normalized coverage [%] | genome coverage [%] | mapped reads | total bases reads | relative read abundance [%] | normalized coverage [%] |
| | 19.03.2018 | 78 | 132647062 | 3.216E+10 | 0.2 -3 µm | 0.75 | 1806 | 421932 | 0.0014 | 0.0034 | NA | 0 | 0 | 0.0000 | 0.0000 | NA | 0 | 0 | 0.0000 | 0.0000 |
| | 03.04.2018 | 93 | 130734852 | 3.094E+10 | 0.2 -3 µm | 6.63 | 3860 | 907066 | 0.0030 | 0.0077 | 2.26 | 5 | 1192 | 0.0000 | 0.0009 | 0.91 | 2 | 496 | 0.0000 | 0.0004 |
| | 05.04.2018 | 95 | 130836262 | 3.131E+10 | 0.2 -3 µm | 0.61 | 2580 | 592503 | 0.0020 | 0.0050 | NA | 0 | 0 | 0.0000 | 0.0000 | NA | 0 | 0 | 0.0000 | 0.0000 |
| | 10.04.2018 | 100 | 131511730 | 3.167E+10 | 0.2 -3 µm | 0.91 | 2373 | 551176 | 0.0018 | 0.0046 | NA | 0 | 0 | 0.0000 | 0.0000 | NA | 0 | 0 | 0.0000 | 0.0000 |
| | 12.04.2018 | 102 | 137437816 | 3.31E+10 | 0.2 -3 µm | 0.72 | 2511 | 585454 | 0.0018 | 0.0046 | NA | 0 | 0 | 0.0000 | 0.0000 | NA | 0 | 0 | 0.0000 | 0.0000 |
| | 17.04.2018 | 108 | 134007214 | 3.216E+10 | 0.2 -3 µm | 2.10 | 4162 | 959789 | 0.0031 | 0.0078 | NA | 0 | 0 | 0.0000 | 0.0000 | NA | 0 | 0 | 0.0000 | 0.0000 |
| | 19.04.2018 | 110 | 145428602 | 3.516E+10 | 0.2 -3 µm | 0.32 | 3307 | 772074 | 0.0023 | 0.0058 | NA | 0 | 0 | 0.0000 | 0.0000 | NA | 0 | 0 | 0.0000 | 0.0000 |
| | 24.04.2018 | 115 | 148226228 | 3.577E+10 | 0.2 -3 µm | 0.70 | 3271 | 760049 | 0.0022 | 0.0056 | NA | 0 | 0 | 0.0000 | 0.0000 | NA | 0 | 0 | 0.0000 | 0.0000 |
| | 26.04.2018 | 117 | 186004950 | 4.485E+10 | 0.2 -3 µm | 0.93 | 5333 | 1250110 | 0.0029 | 0.0073 | 0.59 | 1 | 250 | 0.0000 | 0.0001 | NA | 0 | 0 | 0.0000 | 0.0000 |
| | 02.05.2018 | 123 | 172417734 | 4.171E+10 | 0.2 -3 µm | 9.43 | 9073 | 2102765 | 0.0053 | 0.0133 | 3.76 | 7 | 1732 | 0.0000 | 0.0010 | NA | 0 | 0 | 0.0000 | 0.0000 |
| | 03.05.2018 | 124 | 175408924 | 4.239E+10 | 0.2 -3 µm | 1.45 | 6294 | 1481953 | 0.0036 | 0.0092 | 0.36 | 1 | 155 | 0.0000 | 0.0001 | NA | 0 | 0 | 0.0000 | 0.0000 |
| | 08.05.2018 | 129 | 157697368 | 3.823E+10 | 0.2 -3 µm | 0.62 | 6111 | 1442289 | 0.0039 | 0.0099 | NA | 0 | 0 | 0.0000 | 0.0000 | NA | 0 | 0 | 0.0000 | 0.0000 |
| | 11.05.2018 | 134 | 174367208 | 4.196E+10 | 0.2 -3 µm | 0.29 | 5321 | 1237808 | 0.0031 | 0.0078 | NA | 0 | 0 | 0.0000 | 0.0000 | NA | 0 | 0 | 0.0000 | 0.0000 |
| | 15.05.2018 | 136 | 167144802 | 4.036E+10 | 0.2 -3 µm | 0.30 | 4826 | 1124904 | 0.0029 | 0.0073 | NA | 0 | 0 | 0.0000 | 0.0000 | NA | 0 | 0 | 0.0000 | 0.0000 |
| | 17.05.2018 | 142 | 178481766 | 4.291E+10 | 0.2 -3 µm | 0.39 | 4426 | 1034588 | 0.0025 | 0.0063 | NA | 0 | 0 | 0.0000 | 0.0000 | NA | 0 | 0 | 0.0000 | 0.0000 |
| | 22.05.2018 | 143 | 162504646 | 3.904E+10 | 0.2 -3 µm | 0.79 | 5674 | 1314451 | 0.0035 | 0.0089 | NA | 0 | 0 | 0.0000 | 0.0000 | NA | 0 | 0 | 0.0000 | 0.0000 |
| | 24.05.2018 | 145 | 159197066 | 3.831E+10 | 0.2 -3 µm | 0.43 | 7626 | 1773688 | 0.0048 | 0.0122 | NA | 0 | 0 | 0.0000 | 0.0000 | NA | 0 | 0 | 0.0000 | 0.0000 |
| | 29.05.2018 | 150 | 171271466 | 4.143E+10 | 0.2 -3 µm | 0.39 | 5813 | 1358236 | 0.0034 | 0.0086 | NA | 0 | 0 | 0.0000 | 0.0000 | NA | 0 | 0 | 0.0000 | 0.0000 |
| | 19.03.2018 | 78 | 408373164 | 9.731E+10 | 3-10 µm | 2.09 | 2374 | 553524 | 0.0006 | 0.0015 | NA | 0 | 0 | 0.0000 | 0.0000 | NA | 0 | 0 | 0.0000 | 0.0000 |
| | 12.04.2018 | 102 | 332059730 | 7.927E+10 | 3-10 µm | 1.02 | 1452 | 343084 | 0.0004 | 0.0011 | 2.49 | 7 | 1603 | 0.0000 | 0.0005 | 1.42 | 3 | 720 | 0.0000 | 0.0002 |
| | 17.04.2018 | 108 | 420871688 | 9.615E+10 | 3-10 µm | 1.78 | 1866 | 419786 | 0.0004 | 0.0011 | 0.59 | 5 | 1212 | 0.0000 | 0.0003 | NA | 0 | 0 | 0.0000 | 0.0000 |
| | 26.04.2018 | 117 | 341246096 | 8.148E+10 | 3-10 µm | 1.95 | 1629 | 384084 | 0.0005 | 0.0012 | 1.06 | 7 | 1124 | 0.0000 | 0.0003 | 1.06 | 6 | 947 | 0.0000 | 0.0003 |
| | 08.05.2018 | 129 | 390032052 | 8.751E+10 | 3-10 µm | 0.38 | 1730 | 375951 | 0.0004 | 0.0011 | 0.61 | 5 | 1177 | 0.0000 | 0.0003 | NA | 0 | 0 | 0.0000 | 0.0000 |
| | 11.05.2018 | 134 | 346400002 | 8.256E+10 | 3-10 µm | 0.21 | 1616 | 383164 | 0.0005 | 0.0012 | NA | 0 | 0 | 0.0000 | 0.0000 | NA | 0 | 0 | 0.0000 | 0.0000 |
| | 22.05.2018 | 143 | 322898796 | 7.6E+10 | 3-10 µm | 0.51 | 3232 | 740108 | 0.0010 | 0.0026 | 0.38 | 10 | 1840 | 0.0000 | 0.0006 | NA | 0 | 0 | 0.0000 | 0.0000 |
| | 29.05.2018 | 150 | 361105792 | 8.634E+10 | 3-10 µm | 0.44 | 2210 | 520325 | 0.0006 | 0.0016 | 0.48 | 1 | 206 | 0.0000 | 0.0001 | 0.48 | 1 | 206 | 0.0000 | 0.0001 |

| | | | | | | | | | | | | | | | | | | | |
|---|---|---|---|---|---|---|---|---|---|---|---|---|---|---|---|---|---|---|---|
| 19.03.2018 | 78 | 229237742 | 5.467E+10 | >10 µm | 0.22 | 1449 | 327028 | 0.0006 | 0.0016 | NA | 0 | 0 | 0.0000 | 0.0000 | NA | 0 | 0 | 0.0000 | 0.0000 |
| 12.04.2018 | 102 | 366243118 | 8.751E+10 | >10 µm | 7.56 | 11438 | 2699344 | 0.0031 | 0.0081 | 99.99 | 12452 | 2975865 | 0.0034 | 0.7978 | 99.98 | 10893 | 2607306 | 0.0030 | 0.6990 |
| 17.04.2018 | 108 | 349480760 | 8.224E+10 | >10 µm | 0.72 | 3032 | 721762 | 0.0009 | 0.0023 | NA | 0 | 0 | 0.0000 | 0.0000 | NA | 0 | 0 | 0.0000 | 0.0000 |
| 26.04.2018 | 117 | 357085094 | 8.282E+10 | >10 µm | 6.39 | 6043 | 1370596 | 0.0017 | 0.0044 | NA | 0 | 0 | 0.0000 | 0.0000 | NA | 0 | 0 | 0.0000 | 0.0000 |
| 08.05.2018 | 129 | 332293300 | 7.447E+10 | >10 µm | 1.36 | 2474 | 526918 | 0.0007 | 0.0019 | 0.42 | 1 | 177 | 0.0000 | 0.0001 | NA | 0 | 0 | 0.0000 | 0.0000 |
| 11.05.2018 | 134 | 340848252 | 8.026E+10 | >10 µm | 0.41 | 1056 | 248855 | 0.0003 | 0.0008 | 1.52 | 4 | 1004 | 0.0000 | 0.0003 | NA | 0 | 0 | 0.0000 | 0.0000 |
| 22.05.2018 | 143 | 416648526 | 9.496E+10 | >10 µm | 4.35 | 5792 | 1292102 | 0.0014 | 0.0036 | 0.62 | 4 | 525 | 0.0000 | 0.0001 | 0.32 | 2 | 267 | 0.0000 | 0.0001 |
| 29.05.2018 | 150 | 369290526 | 7.388E+10 | >10 µm | 1.30 | 4502 | 872547 | 0.0012 | 0.0031 | 0.73 | 5 | 885 | 0.0000 | 0.0003 | NA | 0 | 0 | 0.0000 | 0.0000 |

**Table 13:** Read mapping results of phage Leef and its host

| | | | | | | phage 70% identity | | | | | phage 100% identity | | | | |
|---|---|---|---|---|---|---|---|---|---|---|---|---|---|---|---|
| Leef metagenome | date | Julian Day | total reads | total bases | size fraction | genome coverage [%] | mapped reads | total bases reads | relative read abundance [%] | normalized coverage [%] | genome coverage [%] | mapped reads | total bases reads | relative read abundance [%] | normalized coverage [%] |
| | 19.03.2018 | 78 | 132647062 | 3.216E+10 | 0.2 -3 µm | 1.90 | 3 | 749 | 0.0000 | 0.0006 | NA | 0 | 0 | 0.0000 | 0.0000 |
| | 03.04.2018 | 93 | 130734852 | 3.094E+10 | 0.2 -3 µm | 17.57 | 42 | 10062 | 0.0000 | 0.0087 | 3.33 | 8 | 1964 | 0.0000 | 0.0017 |
| | 05.04.2018 | 95 | 130836262 | 3.131E+10 | 0.2 -3 µm | 2.04 | 4 | 1003 | 0.0000 | 0.0009 | NA | 0 | 0 | 0.0000 | 0.0000 |
| | 10.04.2018 | 100 | 131511730 | 3.167E+10 | 0.2 -3 µm | 2.00 | 3 | 753 | 0.0000 | 0.0006 | NA | 0 | 0 | 0.0000 | 0.0000 |
| | 12.04.2018 | 102 | 137437816 | 3.31E+10 | 0.2 -3 µm | 2.78 | 7 | 1508 | 0.0000 | 0.0012 | 1.11 | 2 | 455 | 0.0000 | 0.0004 |
| | 17.04.2018 | 108 | 134007214 | 3.216E+10 | 0.2 -3 µm | 3.86 | 6 | 1471 | 0.0000 | 0.0012 | 2.59 | 4 | 973 | 0.0000 | 0.0008 |
| | 19.04.2018 | 110 | 145428602 | 3.516E+10 | 0.2 -3 µm | NA | 0 | 0 | 0.0000 | 0.0000 | | 0 | 0 | 0.0000 | 0.0000 |
| | 24.04.2018 | 115 | 148226228 | 3.577E+10 | 0.2 -3 µm | 2.47 | 4 | 939 | 0.0000 | 0.0007 | NA | 0 | 0 | 0.0000 | 0.0000 |
| | 26.04.2018 | 117 | 186004950 | 4.485E+10 | 0.2 -3 µm | 9.61 | 28 | 6592 | 0.0000 | 0.0039 | NA | 0 | 0 | 0.0000 | 0.0000 |
| | 02.05.2018 | 123 | 172417734 | 4.171E+10 | 0.2 -3 µm | 18.37 | 48 | 11750 | 0.0000 | 0.0075 | 2.36 | 5 | 1236 | 0.0000 | 0.0008 |
| | 03.05.2018 | 124 | 175408924 | 4.239E+10 | 0.2 -3 µm | 5.68 | 13 | 2945 | 0.0000 | 0.0019 | 1.19 | 3 | 641 | 0.0000 | 0.0004 |
| | 08.05.2018 | 129 | 157697368 | 3.823E+10 | 0.2 -3 µm | 6.67 | 14 | 3451 | 0.0000 | 0.0024 | NA | 0 | 0 | 0.0000 | 0.0000 |
| | 11.05.2018 | 134 | 174367208 | 4.196E+10 | 0.2 -3 µm | 2.17 | 4 | 955 | 0.0000 | 0.0006 | 0.54 | 1 | 204 | 0.0000 | 0.0001 |
| | 15.05.2018 | 136 | 167144802 | 4.036E+10 | 0.2 -3 µm | 2.90 | 5 | 1199 | 0.0000 | 0.0008 | NA | 0 | 0 | 0.0000 | 0.0000 |
| | 17.05.2018 | 142 | 178481766 | 4.291E+10 | 0.2 -3 µm | 1.99 | 3 | 749 | 0.0000 | 0.0005 | NA | 0 | 0 | 0.0000 | 0.0000 |
| | 22.05.2018 | 143 | 162504646 | 3.904E+10 | 0.2 -3 µm | 6.50 | 14 | 3489 | 0.0000 | 0.0024 | 1.33 | 3 | 750 | 0.0000 | 0.0005 |
| | 24.05.2018 | 145 | 159197066 | 3.831E+10 | 0.2 -3 µm | 0.60 | 2 | 445 | 0.0000 | 0.0003 | NA | 0 | 0 | 0.0000 | 0.0000 |
| | 29.05.2018 | 150 | 171271466 | 4.143E+10 | 0.2 -3 µm | 4.70 | 12 | 2757 | 0.0000 | 0.0018 | NA | 0 | 0 | 0.0000 | 0.0000 |

| | | | | | | | | | | | | | | | | |
|---|---|---|---|---|---|---|---|---|---|---|---|---|---|---|---|---|
| 19.03.2018 | 78 | 408373164 | 9.731E+10 | 3-10 µm | | 14.71 | 61 | 14854 | 0.0000 | 0.0041 | | 1.34 | 5 | 1252 | 0.0000 | 0.0003 |
| 12.04.2018 | 102 | 332059730 | 7.927E+10 | 3-10 µm | | 7.97 | 23 | 5501 | 0.0000 | 0.0018 | NA | | 0 | 0 | 0.0000 | 0.0000 |
| 17.04.2018 | 108 | 420871688 | 9.615E+10 | 3-10 µm | | 9.07 | 36 | 9030 | 0.0000 | 0.0025 | | 1.20 | 4 | 1004 | 0.0000 | 0.0003 |
| 26.04.2018 | 117 | 341246096 | 8.148E+10 | 3-10 µm | | 17.39 | 54 | 12787 | 0.0000 | 0.0042 | | 1.58 | 4 | 1002 | 0.0000 | 0.0003 |
| 08.05.2018 | 129 | 390032052 | 8.751E+10 | 3-10 µm | | 2.67 | 6 | 1506 | 0.0000 | 0.0005 | NA | | 0 | 0 | 0.0000 | 0.0000 |
| 11.05.2018 | 134 | 346400002 | 8.256E+10 | 3-10 µm | | 1.26 | 4 | 856 | 0.0000 | 0.0003 | NA | | 0 | 0 | 0.0000 | 0.0000 |
| 22.05.2018 | 143 | 322898796 | 7.6E+10 | 3-10 µm | | 1.33 | 8 | 2003 | 0.0000 | 0.0007 | NA | | 0 | 0 | 0.0000 | 0.0000 |
| 29.05.2018 | 150 | 361105792 | 8.634E+10 | 3-10 µm | | 8.94 | 19 | 4492 | 0.0000 | 0.0014 | NA | | 0 | 0 | 0.0000 | 0.0000 |
| 19.03.2018 | 78 | 229237742 | 5.467E+10 | >10 µm | NA | | 0 | 0 | 0.0000 | 0.0000 | | | 0 | 0 | 0.0000 | 0.0000 |
| 12.04.2018 | 102 | 366243118 | 8.751E+10 | >10 µm | | 31.54 | 12562 | 2964733 | 0.0034 | 0.9023 | | 9.73 | 47 | 11267 | 0.0000 | 0.0034 |
| 17.04.2018 | 108 | 349480760 | 8.224E+10 | >10 µm | | 1.81 | 10 | 2361 | 0.0000 | 0.0008 | NA | | 0 | 0 | 0.0000 | 0.0000 |
| 26.04.2018 | 117 | 357085094 | 8.282E+10 | >10 µm | | 28.24 | 123 | 28151 | 0.0000 | 0.0091 | | 3.04 | 9 | 1940 | 0.0000 | 0.0006 |
| 08.05.2018 | 129 | 332293300 | 7.447E+10 | >10 µm | | 10.89 | 30 | 6873 | 0.0000 | 0.0025 | | 1.50 | 3 | 748 | 0.0000 | 0.0003 |
| 11.05.2018 | 134 | 340848252 | 8.026E+10 | >10 µm | | 1.06 | 2 | 501 | 0.0000 | 0.0002 | NA | | 0 | 0 | 0.0000 | 0.0000 |
| 22.05.2018 | 143 | 416648526 | 9.496E+10 | >10 µm | | 61.98 | 301 | 70233 | 0.0001 | 0.0197 | | 33.33 | 110 | 26002 | 0.0000 | 0.0073 |
| 29.05.2018 | 150 | 369290526 | 7.388E+10 | >10 µm | | 5.65 | 42 | 7238 | 0.0000 | 0.0026 | NA | | 0 | 0 | 0.0000 | 0.0000 |

**Table 14:** Isolation and cultivation specifics of eight strains obtained from two particle fractions at Helgoland Roads (54°11'03"N, 7°54'00"E) during mid-March and mid-May 2017.

| Genus | Strain | DSMZ accession number | Sampling date | Medium |
|---|---|---|---|---|
| ***Polaribacter*** | AHE13PA | DSM111061 | 15.03.2017 | Laminarin |
| ***Tenacibaculum*** | AHE14PA | DSM111040 | 15.03.2017 | Laminarin |
| ***Tenacibaculum*** | AHE15PA | DSM111039 | 15.03.2017 | Laminarin |
| ***Winogradskyella*** | AHE16PA | | 15.05.2017 | 2216 |
| ***Mesonia*** | AHE17PA | | 15.03.2017 | 2216 |
| ***Marixanthomonas*** | AHE18PA | | 15.03.2017 | Laminarin |
| ***Arenibacter*** | AHE19PA | | 15.03.2017 | 2216 |
| ***Polaribacter*** | AHE20PA | | 15.05.2017 | Laminarin |

20. Laslett D, Canback B. ARAGORN, a program to detect tRNA genes and tmRNA genes in nucleotide sequences. Nucleic Acids Research. 2004;32(1):11-6.
21. Jones P, Binns D, Chang H-Y, Fraser M, Li W, McAnulla C, et al. InterProScan 5: genome-scale protein function classification. Bioinformatics (Oxford, England). 2014;30(9):1236-40. Epub 2014/01/21.
22. Kearse M, Moir R, Wilson A, Stones-Havas S, Cheung M, Sturrock S, et al. Geneious Basic: an integrated and extendable desktop software platform for the organization and analysis of sequence data. Bioinformatics (Oxford, England). 2012;28(12):1647-9. Epub 2012/04/27.
23. Hahnke RL, Harder J. Phylogenetic diversity of Flavobacteria isolated from the North Sea on solid media. Systematic and Applied Microbiology. 2013;36(7):497-504.
24. Chin C-S, Alexander DH, Marks P, Klammer AA, Drake J, Heiner C, et al. Nonhybrid, finished microbial genome assemblies from long-read SMRT sequencing data. Nature Methods. 2013;10(6):563-9.
25. Koren S, Walenz BP, Berlin K, Miller JR, Bergman NH, Phillippy AM. Canu: scalable and accurate long-read assembly via adaptive k-mer weighting and repeat separation. Genome Res. 2017.
26. Rodriguez-R LM, Gunturu S, Harvey WT, Rosselló-Mora R, Tiedje JM, Cole JR, et al. The Microbial Genomes Atlas (MiGA) webserver: taxonomic and gene diversity analysis of Archaea and Bacteria at the whole genome level. Nucleic Acids Research. 2018;46(W1):W282-W8.
27. Rodriguez-R LM, Konstantinidis KT. The enveomics collection: a toolbox for specialized analyses of microbial genomes and metagenomes. PeerJ Preprints. 2016;4:e1900v1.
28. Holmfeldt K, Middelboe M, Nybroe O, Riemann L. Large Variabilities in Host Strain Susceptibility and Phage Host Range Govern Interactions between Lytic Marine Phages and Their Flavobacterium Hosts. Applied and Environmental Microbiology. 2007;73(21):6730-9.
29. Holmfeldt K, Odić D, Sullivan MB, Middelboe M, Riemann L. Cultivated Single-Stranded DNA Phages That Infect Marine Bacteroidetes Prove Difficult To Detect with DNA-Binding Stains. Applied and Environmental Microbiology. 2012;78(3):892-4.
30. Roux S, Brum JR, Dutilh BE, Sunagawa S, Duhaime MB, Loy A, et al. Ecogenomics and potential biogeochemical impacts of globally abundant ocean viruses. Nature. 2016;537(7622):689-93.
31. Panschin I, Becher M, Verbarg S, Spröer C, Rohde M, Schüler M, et al. Description of Gramella forsetii sp. nov., a marine Flavobacteriaceae isolated from North Sea water, and emended description of Gramella gaetbulicola Cho et al. 2011. International Journal of Systematic and Evolutionary Microbiology. 2017;67(3):697-703.
32. Alejandre-Colomo C, Harder J, Fuchs BM, Rosselló-Móra R, Amann R. High-throughput cultivation of heterotrophic bacteria during a spring phytoplankton bloom in the North Sea. Systematic and Applied Microbiology. 2020;43(2):126066.
33. Barbeyron T, Carpentier F, apos, Haridon S, Schüler M, Michel G, et al. Description of Maribacter forsetii sp. nov., a marine Flavobacteriaceae isolated from North Sea water, and emended description of the genus Maribacter. International Journal of Systematic and Evolutionary Microbiology. 2008;58(4):790-7.
